# Complex Polyploids: Origins, Genomic Composition, and Role of Introgressed Alleles

**DOI:** 10.1101/2023.08.23.553805

**Authors:** J. Luis Leal, Pascal Milesi, Eva Hodková, Qiujie Zhou, Jennifer James, D. Magnus Eklund, Tanja Pyhäjärvi, Jarkko Salojärvi, Martin Lascoux

**Affiliations:** Plant Ecology and Evolution, Department of Ecology and Genetics, Uppsala University, Norbyvägen 18D, 75236 Uppsala, Sweden; Science for Life Laboratory (SciLifeLab), Uppsala University, 75237 Uppsala, Sweden; Faculty of Environmental Sciences, Czech University of Life Sciences Prague, Kamýcká 129, 16521 Prague, Czech Republic; Physiological Botany, Department of Organismal Biology, Linnean Centre for Plant Biology in Uppsala, Uppsala University, Ulls väg 24E, 75651 Uppsala, Sweden; Organismal and Evolutionary Biology Research Program, Faculty of Biological and Environmental Sciences, and Viikki Plant Science Centre, University of Helsinki, P.O. Box 65 (Viikinkaari 1), 00014 Helsinki, Finland; Department of Forest Sciences, University of Helsinki, 00014 Helsinki, Finland; School of Biological Sciences, Nanyang Technological University, 60 Nanyang Drive, Singapore 637551, Singapore

**Author notes:** Correspondence should be sent to: Plant Ecology and Evolution, Department of Ecology and Genetics, Uppsala University, Norbyvägen 18D, 75236 Uppsala, Sweden.

**Keywords:** allopolyploidy, autopolyploidy, Betula, gene flow, genomic polarization, homoeologs, interploidal, introgressive hybridization, polyploid phylogenetics, polyploidization simulation, reticulate evolution

## Abstract

Introgression allows polyploid species to acquire new genomic content from diploid progenitors or from other unrelated diploid or polyploid lineages, contributing to genetic diversity and facilitating adaptive allele discovery. In some cases, high levels of introgression elicit the replacement of large numbers of alleles inherited from the polyploid’s ancestral species, profoundly reshaping the polyploid’s genomic composition. In such complex polyploids it is often difficult to determine which taxa were the progenitor species and which taxa provided additional introgressive blocks through subsequent hybridization. Here, we use population-level genomic data to reconstruct the phylogenetic history of *Betula pubescens* (downy birch), a tetraploid species often assumed to be of allopolyploid origin and which is known to hybridize with at least four other birch species. This was achieved by modeling of polyploidization and introgression events under the multispecies coalescent and then using an approximate Bayesian computation (ABC) rejection algorithm to evaluate and compare competing polyploidization models. We provide evidence that *B. pubescens* is the outcome of an autoploid genome doubling event in the common ancestor of *B. pendula* and its extant sister species, *B. platyphylla*, that took place approximately 178,000-188,000 generations ago. Extensive hybridization with *B. pendula*, *B. nana*, and *B. humilis* followed in the aftermath of autopolyploidization, with the relative contribution of each of these species to the *B. pubescens* genome varying markedly across the species’ range. Functional analysis of *B. pubescens* loci containing alleles introgressed from *B. nana* identified multiple genes involved in climate adaptation, while loci containing alleles derived from *B. humilis* revealed several genes involved in the regulation of meiotic stability and pollen viability in plant species.

## Introduction

Hybridization and whole genome duplication (WGD) shape evolution in many different ways, facilitating gene flow across distinct lineages (Albertin and Marullo 2012; Dasmahapatra et al. 2012; Ellstrand 2014; Mallet et al. 2016; Schmickl and Yant 2021), giving rise to new species (Dowling and Secor 1997; Ramsey and Schemske 1998; Mallet 2007; Rieseberg and Willis 2007; Soltis and Soltis 2009; Baack et al. 2015; Van de Peer et al. 2017), and fueling adaptive radiations (Seehausen 2006; Keller et al. 2013; Fontaine et al. 2015; Lamichhaney et al. 2015; Meier et al. 2017; Robertson et al. 2017). Homoploid hybridization, the formation of interspecific hybrids without a change in ploidy level, is widespread across many eukaryotic groups (Grant and Grant 1992; Mallet 2005; Meier et al. 2017), having been estimated to affect 25% of all vascular plants and 10% of animal species (Mallet 2005). Polyploids, species whose genome contains more than two complete sets of chromosomes, can arise via whole genome duplication within a single ancestral species (autopolyploids), or from hybridization between two different species concomitant with genome doubling (allopolyploids). Polyploidy is particularly prevalent in plants (Bowers et al. 2003; De Bodt et al. 2005; Wood et al. 2009; Amborella Genome Project 2013; Barker et al. 2016; Van de Peer et al. 2017) but evidence of ancient polyploidization events has been found in many other eukaryotic lineages, including fungi (Wolfe and Shields 1997; Kellis et al. 2004; Ma et al. 2009), vertebrates (Dehal and Boore 2005; Braasch and Postlethwait 2012; Berthelot et al. 2014; Macqueen and Johnston 2014), arthropods (Nossa et al. 2014; Schwager 2017; Li et al. 2018a; Cerca et al. 2021), and ciliates (Aury et al. 2006).

While introgression and polyploidization are generally addressed in the literature as separate biological phenomena, the two processes are not mutually exclusive and often coexist in nature. The occurrence of gene flow from a diploid species to its polyploid descendant, mediated by viable triploid hybrids, was first hypothesized more than sixty years ago by Stebbins (1956), who further noted that interploidal gene flow was expected to be largely unidirectional (Stebbins 1971) with the polyploid species acting as an “allelic sponge” (Schmickl and Yant 2021). A growing body of evidence has emerged suggesting that polyploids can accumulate new alleles not only through introgression with diploid progenitors but also from other, more distant, diploid and polyploid lineages (Marhold and Lihová 2006; Kim et al. 2008; Arnold et al. 2016; Spoelhof et al. 2017; Baduel et al. 2018; Cheng et al. 2019; Marburger et al. 2019; Nieto Feliner et al. 2020; Seear et al. 2020; Schmickl and Yant 2021; Nibau et al. 2022). Introgression via triploid hybrids allows incipient outcrossing polyploids to alleviate the dearth of intraspecific sexual partners (Felber 1991; Ramsey and Schemske 1998; Husband 2004), and contributes to genetic diversity and adaptation (Kim et al. 2008; Slotte et al. 2008; Arnold et al. 2016; Hardigan et al. 2017; Schmickl et al. 2017; Spoelhof et al. 2017; Schmickl and Yant 2021). Recent studies show that introgression can also play an important role in the establishment of meiotic stability in nascent polyploids (Marburger et al. 2019; Seear et al. 2020; Nibau et al. 2022). Introgression involving non-parental species in the aftermath of polyploidization is well-documented in the *Arabidopsis* genus, where it mediates bidirectional gene flow between tetraploid *A. arenosa* and tetraploid *A. lyrata* (Jørgensen et al. 2011; Arnold et al. 2016; Hohmann and Koch 2017; Lafon-Placette et al. 2017; Marburger et al. 2019; Monnahan et al. 2019; Seear et al. 2020). Interploidal gene flow from (non-parental) diploids into tetraploid species has been reported to occur, for example, in bread wheat (*Triticum aestivum*; Cheng et al. 2019), in the *Neobatrachus* genus of Australian burrowing frogs (Novikova et al. 2020), among *Miscanthus* perennial grasses (Clark et al. 2015), and in the *Rorippa* genus of flowering plants (Bleeker and Hurka 2001; Bleeker 2003). Gene flow across species of different ploidy levels has also been observed in the *Betula* (Ashburner and McAllister 2013, pp. 65-75), *Senecio* (Kim et al. 2008), and *Cardamine* (Lihová et al. 2007) genera of flowering plants, but as their reticulate phylogenies remain partially unresolved it is currently not clear whether observed interploidal patterns of gene flow involve any diploid species other than the polyploid’s parental species. This situation in turn calls attention to one unintended consequence of the coexistence of polyploidization and introgression in a single species: although levels of introgression are frequently too low to mask the evolutionary origin of a polyploid, there are instances where its presence may severely interfere with attempts at clarifying its reticulate history.

Few taxa illustrate this predicament better than the birch genus (*Betula* spp.), a group containing several polyploids, high levels of interspecific hybridization, and whose phylogeny has long been marred by uncertainty (Järvinen et al. 2004; Li et al. 2005; Li et al. 2007; Schenk et al. 2008; Wang et al. 2016; Wang et al. 2021). One tetraploid in particular, *B. pubescens*, a monoecious, self-incompatible tree that is widespread across northern Eurasia has been a subject of considerable interest. The high levels of genetic diversity and allelic richness observed in *B. pubescens* are often presented as indirect evidence of allopolyploidization (Howland et al. 1995; Tsuda et al. 2017) since species originating via WGD are generally expected to have levels of genetic diversity in line with values observed in their diploid progenitors (Howland et al. 1995; Palmé et al. 2004; Truong et al. 2007; Tsuda et al. 2017, but see Truong et al. (2007) for arguments for an autopolyploid origin). While most authors tend to agree that *B. pendula* is most likely one of the ancestral parents (Walters 1968; Howland et al. 1995; Järvinen et al. 2004; Tsuda et al. 2017; Wang et al. 2021), the identity of the putative second parent has remained elusive, with different studies providing support for disparate hypotheses. These include *B. nana* (Järvinen et al. 2004; Tsuda et al. 2017), *B. humilis* or a closely related, possibly extinct, species (Walters 1968; Järvinen et al. 2004; Salojärvi et al. 2017; Tsuda et al. 2017), and *B. platyphylla* (Wang et al. 2021). As *B. pubescens* also hybridizes with most of these same species (Anamthawat-Jónsson and Tomasson 1990; Atkinson 1992; Thórsson et al. 2001; Palmé et al. 2004; Jadwiszczak et al. 2012; Wang et al. 2014; Eidesen et al. 2015; Zohren et al. 2016; Tsuda et al. 2017), this combination of polyploidization and extensive introgression makes *B. pubescens* a particularly difficult species to characterize phylogenetically.

In part, this problem stems from a lack of adequate phylogenetic inference tools. The inference of phylogenetic relationships among lineages that contain species of hybrid origin can be performed using specialized tools that explicitly take into consideration the reticulate nature of their phylogenies. However, most currently available tools have been developed *either* to quantify levels of interspecific hybridization (Bryant and Moulton 2004, Huson and Bryant 2006; Than et al. 2008; Pickrell and Pritchard 2012; Solís-Lemus et al. 2017; Wen et al. 2018; Zhang et al. 2018) *or* to identify the parental species of a polyploid (Jones et al. 2013; Bertrand et al. 2015; Oberprieler et al. 2017; Oxelman et al. 2017; Lautenschlager et al. 2020; Rothfels 2021; Yan et al. 2022; Freyman et al. 2023; Leal et al. 2023) − but not both simultaneously. There are also methods that can be used to determine whether a polyploid originated via autopolyploidization or allopolyploidization (Chenuil et al. 1999; Roux and Pannell 2015; Winkler et al. 2017; Muñoz-Rodríguez et al. 2018), but again these assume relatively low levels of gene flow from additional species. Despite their success in unveiling the evolutionary history of many polyploids and homoploid hybrids, the methods listed above would struggle to disentangle polyploidization from introgression if faced with highly hybridized polyploids. In fact, to our knowledge, no method has yet been specifically developed to identify and quantify the genomic components of species that are the outcome of complex polyploidization: auto- or allopolyploids that were subsequently subjected to extensive introgression with one or more additional species (that is, with species other than the polyploid’s parental species).

Here, we use a novel approach to disentangle introgression from polyploidization in order to determine the mode of origin and reconstruct the reticulate phylogeny of the highly hybridized polyploid *B. pubescens*. This is achieved by employing genomic polarization, a phasing algorithm that sorts gene copies in polyploids according to their evolutionary history (Leal et al. 2023), and by modeling polyploidization and introgression events under the multispecies coalescent. An approximate Bayesian computation (ABC) rejection algorithm is then used to evaluate and compare competing polyploidization models. We carried out an analysis of 1,603 exonic sequences in 269 *B. pubescens* specimens collected from twenty-three locations along the species’ range. Our results provide new insights into the evolutionary origin of *B. pubescens*, its genomic composition, and the probable role of alleles introgressed from *B. nana* and *B. humilis*.

## Materials and Methods

### Conceptual Framework

Inference of the evolutionary history of the tetraploid *B. pubescens* was based on the phylogenetic analysis of multiple sequence alignments (MSAs), one for each locus, where the nucleotide sequence associated to *B. pubescens* has been polarized prior to tree inference. Genomic polarization is a recently developed iterative subgenome phasing algorithm for tetraploid species (Leal et al. 2023) that captures allelic variants in a polyploid species *not* present in a reference sequence, usually one of the other sequences present in the MSA. If the reference sequence coincides with one of the polyploid’s parental species, the polarized polyploid sequence shows high sequence identity with the second parental species, pairing with it during subsequent phylogenetic analysis.

Of particular interest to us, cumulative polyploid pairing patterns measured across hundreds of loci are expected to vary not only according to the reference sequence selected, but also to depend on the mode of polyploidization and levels of introgression (Fig. 1). For example, in the absence of introgression, a PPHH allopolyploidization model (where *B. pendula* and *B. humilis* are assumed to be the parental species) predicts pairing patterns where the tetraploid overwhelmingly pairs with *B. humilis* when *B. pendula* is used as the reference sequence, and vice-versa (model 1 in Figure 1). Occasional pairing with non-parental species (peaks shown in light blue in Figure 1) may also occur as a consequence of incomplete lineage sorting (ILS).

**Figure 1.**
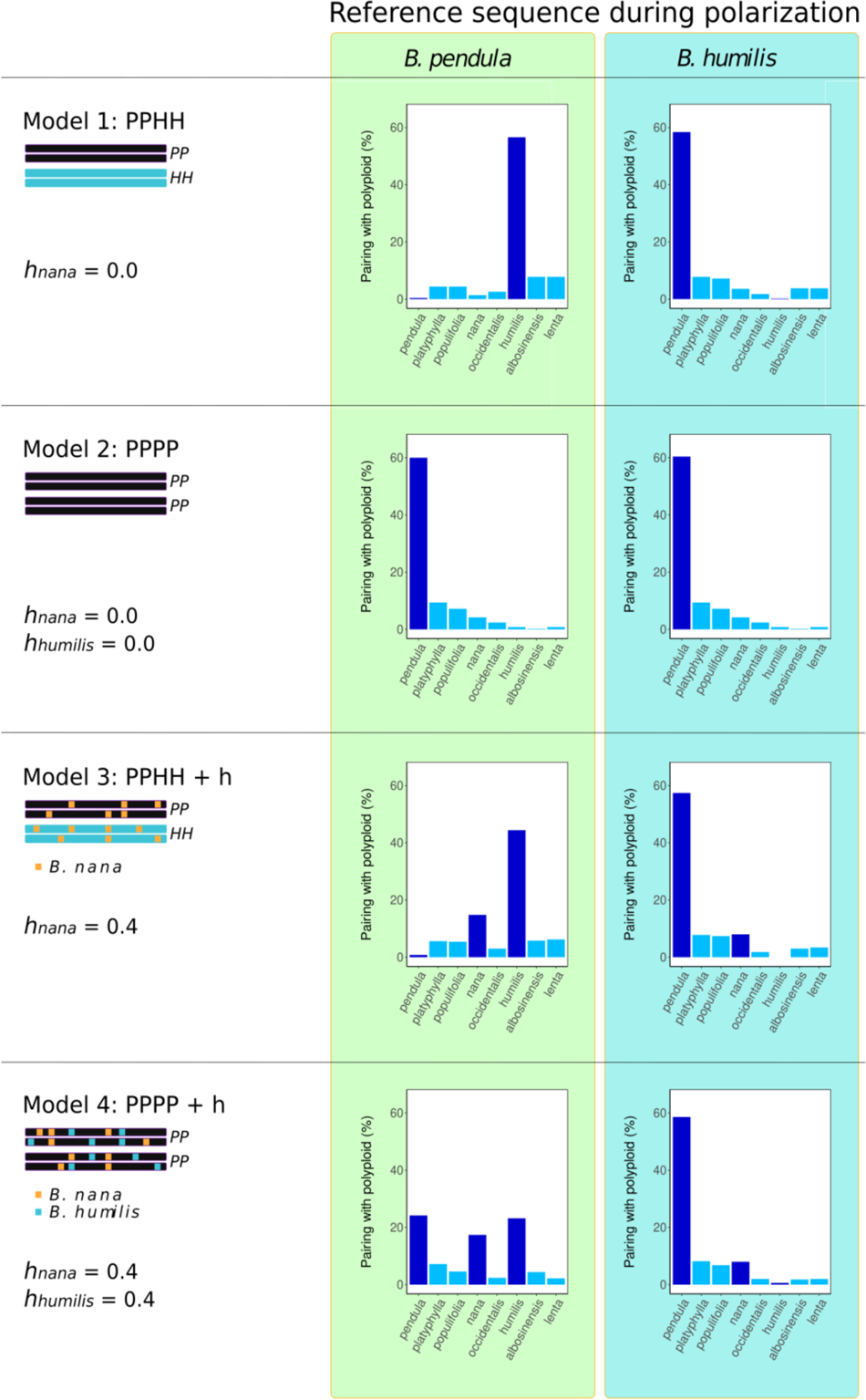
Schematic representation of polyploid pairing patterns for different polyploidization and hybridization models. Barplots show the frequency with which *B. pubescens* is expected to pair with other birch species, across multiple loci, for different evolutionary models and polarization settings. PP and HH indicate the hypothetical evolutionary origin of each subgenome component in the tetraploid (*B. pendula* and *B. humilis*, respectively, in the examples provided). Levels of introgressive hybridization are indicated by *h*.

Instead, the PPPP autopolyploid model (*B. pendula* as the sole parental species; no hybridization) predicts pairing patterns with a single peak observed for *B. pendula*, a pattern which remains invariant irrespective of the species used during polarization (model 2 in Figure 1). The presence of introgression is expected to affect pairing patterns in both allo- and autopolyploids (models 3 and 4 in Figure 1). The location of the extra peaks and their intensity provide clues as to the identity of the species with which the polyploid hybridizes, and the degree of introgression. By comparing observed polyploid pairing patterns with those obtained through polyploidization and introgression modeling, it should therefore become possible to not only source the different components of a polyploid’s genome but also to determine its evolutionary origin, that is, to establish which taxa were the original progenitor species, and which taxa provided additional introgressed genome blocks through subsequent hybridization.

### Plant Material, DNA Isolation, Library Preparation and Sequencing

Bud and leaf tissues were collected from natural populations for 300 birch individuals, 269 of which were subsequently identified as *B. pubescens,* 25 as *B. pendula* autotetraploids (4X), and 2 as triploid hybrids of different origin (see *Classification of Polyploid Specimens* below). Species and ploidy could not be conclusively determined for four samples. Some of these samples (44) were collected and sequenced as part of a previous study (Opgenoorth et al. 2021; Milesi et al. 2023). *B. pubescens* specimens were sampled from twenty-three locations across the species range, with a particular focus on Scandinavia (Supplementary Fig. S1). Leaf material was also obtained from *B. albosinensis*, *B. humilis*, *B. nana*, and *B. platyphylla* specimens. See Supplementary Table S1 and *Supplementary Materials 2* for further details.

Total DNA was extracted from bud and leaf tissues using a DNeasy Plant Mini Kit (Qiagen AG, Düsseldorf, Germany). Individual libraries were generated using the SeqCap EZ HyperCapWorkflow (Roche AG, Basel, Switzerland) using custom probes designed for targeted exome capture (TEC) and sequenced on paired-end mode (150 bp) on an Illumina NovaSeq 6000 platform (Illumina, San Diego, CA). Target resequencing was based on a custom probe design covering 1,609 nuclear genes with a 3.2M bp cumulative target size and reduced sequence redundancy (Milesi et al. 2023). See *Supplementary Materials 1* for further details. Additional accessions were obtained from The European Nucleotide Archive (ENA, https://www.ebi.ac.uk/ena) for *B. lenta*, *B. nana*, *B. occidentalis*, *B. pendula*, *B. platyphylla*, *B. populifolia*, *B. pubescens*, *Alnus glutinosa*, *Alnus incana*, and a further twenty *B. pendula* specimens, which were part of previous studies (Salojärvi et al., 2017; Milesi et al. 2023). Further details about each accession can be found in Supplementary Table S1 and *Supplementary Materials 1 & 2*.

### Processing of Sequencing Libraries and Generation of Multiple Sequence Alignments (MSAs)

#### Read mapping and variant calling

Sequenced reads were mapped to the *B. pendula* genome assembly, v Bpev01 (Salojärvi et al. 2017), using BWA-MEM v 0.7.17 (Li and Durbin 2009). Chromosomal assignment of loci was performed on the basis of the *B. pendula* chromosomal map, v 1.4c (https://genomevolution.org/CoGe/GenomeInfo.pl?gid=35080, accessed 2023-01-26). Genotyping was performed with GATK 4.2.0.0 (McKenna et al. 2010). For all birch and alder accessions, including those obtained through WGS, only sites located within the 1,603 nuclear loci covered by the exome probes were considered (six of the targeted 1,609 loci had very low coverage and were excluded). Sites located outside the targeted regions were excluded during downstream analysis. See *Supplementary Materials 1 & 3* for further details.

#### Classification of polyploid specimens

*B. pubescens* and *B. pendula* specimens collected in the wild are often difficult to tell apart based solely on morphology (Ashburner and McAllister 2013, p. 232). Autotriploid and autotetraploid *B. pendula* variants have also been reported to occur under natural conditions (Johnsson 1944; Zohren et al. 2016). We classified each sample according to its ploidy level (2X, 3X, or 4X) by computing the ratio between the number of reads supporting the reference and alternate alleles at biallelic variant sites (distribution of read counts), as described previously in Yoshida et al. (2013) and Zohren et al. (2016); see *Supplementary Materials 1* for further details. Results are shown in Supplementary Figure S2 and *Supplementary Materials 3*.

#### Generation of consensus sequences and MSAs

Consensus sequences were generated for each accession using bcftools consensus v 1.12 (Danecek et al. 2021) by applying single-nucleotide polymorphisms (SNPs) specific to each individual to the *B. pendula* reference assembly. Heterozygous sites were coded following the IUPAC nomenclature (Cornish-Bowden 1985). As only invariant sites and point mutations were used when producing each consensus sequence, consensus sequences associated to different samples are aligned by default: they all have the same length and retain identical start positions for each gene, and therefore require no further alignment (Leal et al. 2023). Separate MSAs were produced for each of the 1,603 loci included in the analysis by splitting each consensus sequence using getfasta from the bedtools suite, based on the gff3 annotation file associated to the *B. pendula* reference assembly (Salojärvi et al. 2017). Each MSA contains eight birch species (*B. albosinensis, B. humilis, B. lenta*, *B. nana*, *B. occidentalis*, *B. pendula*, *B. platyphylla*, and *B. populifolia*) and two alder species (*A. glutinosa* and *A. incana*). Additionally, each MSA contains a nucleotide sequence associated to one of the 269 *B. pubescens* samples included in this study, with each *pubescens* specimen and locus therefore being the subject of a separate phylogenetic analysis. See *Supplementary Materials 1* for further details.

### Data Analysis

#### Genomic polarization

Genomic polarization (Leal et al. 2023) of the focal *B. pubescens* nucleotide sequence was performed directly on each MSA using the PolyAncestor pipeline (https://github.com/LLN273/PolyAncestor) before carrying out phylogenetic analysis (Supplementary Figure S3). We performed independent polarizations based on four reference species: *B. pendula*, *B. platyphylla*, *B. humilis*, and *B. nana*. During polarization, the nucleotide sequence associated to the focal *B. pubescens* sample is scanned nucleotide by nucleotide, and variants at polymorphic sites found to also be present in the reference sequence are filtered out. This does not apply to sites in the focal *B. pubescens* sequence that are fixed, masked, or whose allelic composition exactly matches that observed in the reference sequence (in all three cases, the polarized sequence remains identical to the unpolarized *B. pubescens* sequence). Sites in the polarized sequence were recoded using the IUPAC nomenclature. Note that the terms *reference sequence* and *reference genome* have distinct meanings in the context of this paper. The former refers to the sequence in the MSA used as a reference during genomic polarization, while the latter refers to the *B. pendula* reference assembly used during read mapping, variant calling, and generation of consensus sequences.

#### Phylogenetic analysis

Phylogenetic reconstruction of individual gene families was performed using maximum-likelihood as implemented in IQ-TREE2 v 2.0-rc2-omp-mpi (Minh et al. 2020). Polyploid pairing profiles were produced by counting the number of times the polarized polyploid species pairs with any of the other birch species included in the phylogeny, based on the analysis of individual gene trees (IQ-TREE2 output). This analysis was carried out independently for each of the 269 *B. pubescens* samples collected. Next, polyploid pairing profiles were generated for each population/location by averaging across all individuals in the population (Supplementary Figure S3). See *Supplementary Materials 1* for further details.

#### Population structure analysis

Bayesian statistical inference of admixture levels between *B. pubescens* populations and different birch species was performed with STRUCTURE v 2.3.4 (Pritchard et al. 2000), based on the analysis of 10,000 variant loci in 126 birch accessions. The presence of population genetic structure was also investigated using discriminant analysis of principal components (DAPC; Jombart et al. 2010) with adegenet v 2.1.7, based on 50,000 variant loci belonging to 49 birch accessions. See Supplementary Materials 1 for further details.

### Model Testing

In order to evaluate competing polyploidization models, we generated synthetic MSAs based on different polyploidization modes and levels of hybridization, and compared their associated pairing profiles to those observed experimentally using an approximate Bayesian computation (ABC) rejection algorithm.

#### Polyploidization and hybridization simulations

We produced synthetic MSAs where the focal polyploid, representing *B. pubescens*, was generated based on different polyploidization and hybridization models (Supplementary Figure S3). We considered the following base polyploidization scenarios: PPPP, AAAA, PPNN, PPHH, PPNH, PPP_y_P_y_, AANN, AAHH, and AANH, where character pairs indicate the origin of each of the tetraploid’s subgenomes (PP: *B. pendula*; AA: the common ancestor of *B. pendula* and *B. platyphylla*; NN: *B. nana*; HH: *B. humilis*; P_y_P_y_: *B. platyphylla*; NH: *B. nana* x *B. humilis* hybrid). The first two cases (PPPP and AAAA) indicate an autopolyploid origin, while all other cases assume *B. pubescens* to be an allopolyploid. For each of these nine models, we then simulate different scenarios where the polyploid gains new alleles through introgressive hybridization with one or more third species.

We used SimPhy v 1.0.2 (Mallo et al. 2016), a phylogeny simulator based on the multispecies coalescent model, to generate gene family trees evolving under ILS (based on a species tree provided by the user). Initial simulation conditions were similar to those used by Zhang et al. (2018) and Leal et al. (2023) to simulate a moderate ILS scenario. A homemade script was used to further adjust (reduce) genus-wide levels of ILS. When running SimPhy, the initial species-wide phylogeny was kept fixed and mimicked the observed *Betulaceae* tree for the species of interest, to which five extra branches were added: the allopolyploid’s putative ancestral species, and the species with which the allopolyploid can potentially hybridize (Supplementary Fig. S5). Two haploid individuals were created per species. Twenty-five independent replicates were simulated for each of the five polyploidization scenarios, each containing fifty gene families, producing a grand total of 1,250 gene families per model.

We generated synthetic MSAs for each gene family using INDELible v 1.03 (Fletcher and Yang 2009) based on the gene trees created with SimPhy. Mean nucleotide sequence length and standard deviation were set to 1,500 and 100 bp, respectively. For exact simulations conditions, see Leal et al. (2023).

Once the MSAs were generated, the two haploid individuals belonging to the same species were used to create diploid individuals, and the allopolyploid was created using the nucleotide sequences associated to the two hypothetical parental species (e.g., ‘ANCESTRAL N’ and ‘ANCESTRAL PEN’ in Supplementary Figure S5). For each gene in the tetraploid assumed to have undergone introgressive hybridization (such loci were selected randomly), one of the putative ancestral species was replaced by the sequence associated to the species with which the tetraploid hybridizes (e.g. ‘ANCESTRAL H’). Diverse evolutionary scenarios can therefore be simulated by selecting different parental pairs, the identity of one or more hybridizing species, and hybridization levels. All putative ancestral sequences (*H*, *N*, *PLA*, *PEN*, and *ANC*, see Supplementary Figure S5) were dropped from the MSA prior to phylogenetic inference. Genomic polarization of the synthetic *B. pubescens* sequence and the subsequent phylogenetic analysis of individual gene families with IQ-TREE2 were performed as described above. All genes from all replicates (1,250 genes) were used during the analysis of polyploid pairing patterns.

When modeling the allopolyploidization process, it was assumed that alleles obtained through introgression can be shared across subgenomes (as opposed to being confined to a particular subgenome). While bivalent pairing is predominant during meiotic prophase I in *B. pubescens* (Brown and Al-Dawoody 1979), observation of segregation patterns revealed that this species exhibits polysomic inheritance (Stern 1965), suggesting that genetic recombination can occur between homoeologous chromosomes associated to different subgenomes.

### Model evaluation

Evaluation of each of the nine base polyploidization models was done using an approximate Bayesian computation (ABC) rejection algorithm (Tavaré et al. 1997; Beaumont et al. 2002; Beaumont 2019). Polyploid pairing profiles observed experimentally for four polarization geometries (polarization performed using *B. pendula*, *B. platyphylla*, *B. nana*, and *B. humilis* as reference sequences) and across different sampling locations were compared with those produced using simulated data, with the aim of identifying the model that best reproduces the experimental observations across the species range (Supplementary Figure S3). Polyploid pairing frequencies obtained for the four polarizing geometries used therefore constitute our summary statistics. The number of model parameters varied between seven and eleven, depending on the polyploidization model (Supplementary Fig. S6). Most model parameters (H_i_) represent fractions of the genome associated to a specific evolutionary history. Two extra parameters were used to model genus-wide levels of ILS and the level of secondary gene flow and/or ancestral shared polymorphisms between *B. pendula* and *B. platyphylla*. Allopolyploid systems (models 3 to 9) were simulated under two different conditions, either precluding or allowing for the possibility of homoeologous exchange. Initial optimization of model parameters, performed separately for each of the nine polyploidization models and for different populations, was carried out using a simulated annealing algorithm with four independent chains and randomized start model parameters (Supplementary Figure S3). Final optimization was performed using ABC. Batches of 1,000 simulation runs were generated for each model/location with priors sampled from a truncated normal distribution centered on the best model parameters determined using simulated annealing. The weighted *L2* norm distance was used to evaluate the degree of agreement between simulated (y_sim_) and observed (y_obs_) polyploid pairing frequencies:

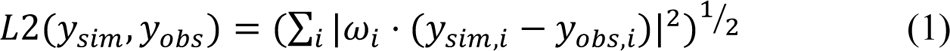

with weights *ω*_i_ set to 5 for the highly variable genomic components (pairing with *B. pendula*, *B. platyphylla*, the *B. pendula/B. platyphylla* clade*, B. nana*, and *B. humilis*), and to 1 in all other cases. Weights were chosen in order to increase L2 sensibility to the highly variable genomic components while accounting as well for background polyploid-pairing values associated to other birch species. The best polyploidization model was selected by identifying the model that consistently best reproduces observed pairing patterns across the *B. pubescens* species range by incurring only plausible local adjustments on the level of hybridization with one or more third species. For each model and location, posterior distributions for each model parameter were estimated based on the top 5% (50) simulation runs with lowest L2 distance. Finally, estimates for the levels of introgression (*h_nana_*, *h_humilis_*) were produced for each model on the basis of the optimized *H_i_* model parameters. Link to scripts used to perform simulated annealing and ABC is provided at the end of the article.

### Demographic Modeling

The speciation time for *B. pubescens* was estimated using coalescent simulations with fastsimcoal2, v 2.7.0.9 (Excoffier et al. 2013), based on *B. pendula*, *B. platyphylla*, and *B. pubescens* SNP data. Only *B. pubescens* loci containing four gene copies of either *B. pendula* or *B. platyphylla* ancestry were included in the analysis (any locus containing one or more gene copies of *B. nana* or *B. humilis* ancestry in any of the *B. pubescens* samples included was filtered out). This ensured that *B. pubescens* loci included in the coalescent analysis all derived from *B. pendula*, *B. platyphylla*, or their common ancestor. In order to minimize data loss due to locus filtering, we sampled only four *B. pubescens* tetraploids and eight diploids for each of *B. pendula* and *B. platyphylla*. The unfolded multidimensional site frequency spectrum (SFS) contained both intronic and four-fold degenerate biallelic sites with a minimum sequencing depth of 8X observed in all samples. Four-fold degenerate sites were identified using SnpEff, v 5.0 (Cingolani et al. 2012). Maximum-likelihood inference of the ancestral allele used when building the SFS was performed with est-sfs, v 2.04 (Keightley and Jackson 2018), using *B. pendula* as the focal species and *B. populifolia*, *B. nana*, and *B. occidentalis* as outgroups. Separate multidimensional SFS were created based on different *B. pubescens* populations (Central Europe, Southern Sweden, Central Asia, and Spain). The speciation time for *B. pubescens* was estimated separately for each *B. pubescens* population. See *Supplementary Materials 1* for further details on model selection and optimization.

### Geographic Distribution and Functional Analysis of Alleles of B. nana and B. humilis Ancestry

The fraction of *B. pubescens* individuals in a population containing alleles of *B. nana* or *B. humilis* origin was computed separately for each of the 1,603 genes included in the analysis. Such allelic variants were identified using the genomic polarization protocol (IQ-TREE2 output), as detailed above. Only MSAs containing at least 100 parsimony-informative sites were included in the analysis (see “Phylogenetic Analysis: Detailed Description” in *Supplementary Materials I*). For this particular analysis, and in order to minimize local genetic bias (family effects), each (meta-)population includes all individuals from two or more sampling sites within a certain geographic area but located hundreds of kilometers apart. The only exception is the Southwestern European population (ES), which includes samples from a single location. K-means cluster analysis was used to partition *B. pubescens* loci containing alleles of *B. nana* or *B. humilis* ancestry in groups with a similar abundance profile across geographic locations. See *Supplementary Materials 1* for further details.

As it is common practice when working with non-model plant species, functional annotation of *B. pubescens*/*B. pendula* genes was based on the annotation of homologous genes in the *Arabidopsis thaliana* model species. We used the association file between *A. thaliana* and *B. pendula* genes (based on sequence similarity) released with the *B. pendula* reference assembly (Salojärvi et al. 2017). Functional annotation for *A. thaliana* genes was obtained from the Swiss-Prot collection of manually curated (reviewed) genes listed in the Uniprot database (https://www.uniprot.org/uniprotkb?facets=reviewed%3Atrue&query=%28taxonomy_id%3A3701%29, downloaded October 27, 2022).

## Results

### The Proportion of the Humilis and Nana Genomic Components in B. pubescens Varies Across the Species Range

Analysis of the genomic composition of tetraploid *B. pubescens* specimens was performed using the genomic polarization framework (see *Materials and Methods* and Leal et al. (2023)). Observed pairing patterns for the polarized polyploid, evaluated across hundreds of loci, suggest that at least three species have contributed to the *B. pubescens* genome either as parental species during polyploidization or as a consequence of subsequent introgression: *B. pendula*, *B. nana*, and *B. humilis* (Fig. 2a). The contribution of each of these species varies widely across the *B. pubescens* distribution range (Figs. 2a, 2b and Supplementary Fig. S7).

**Figure 2.**
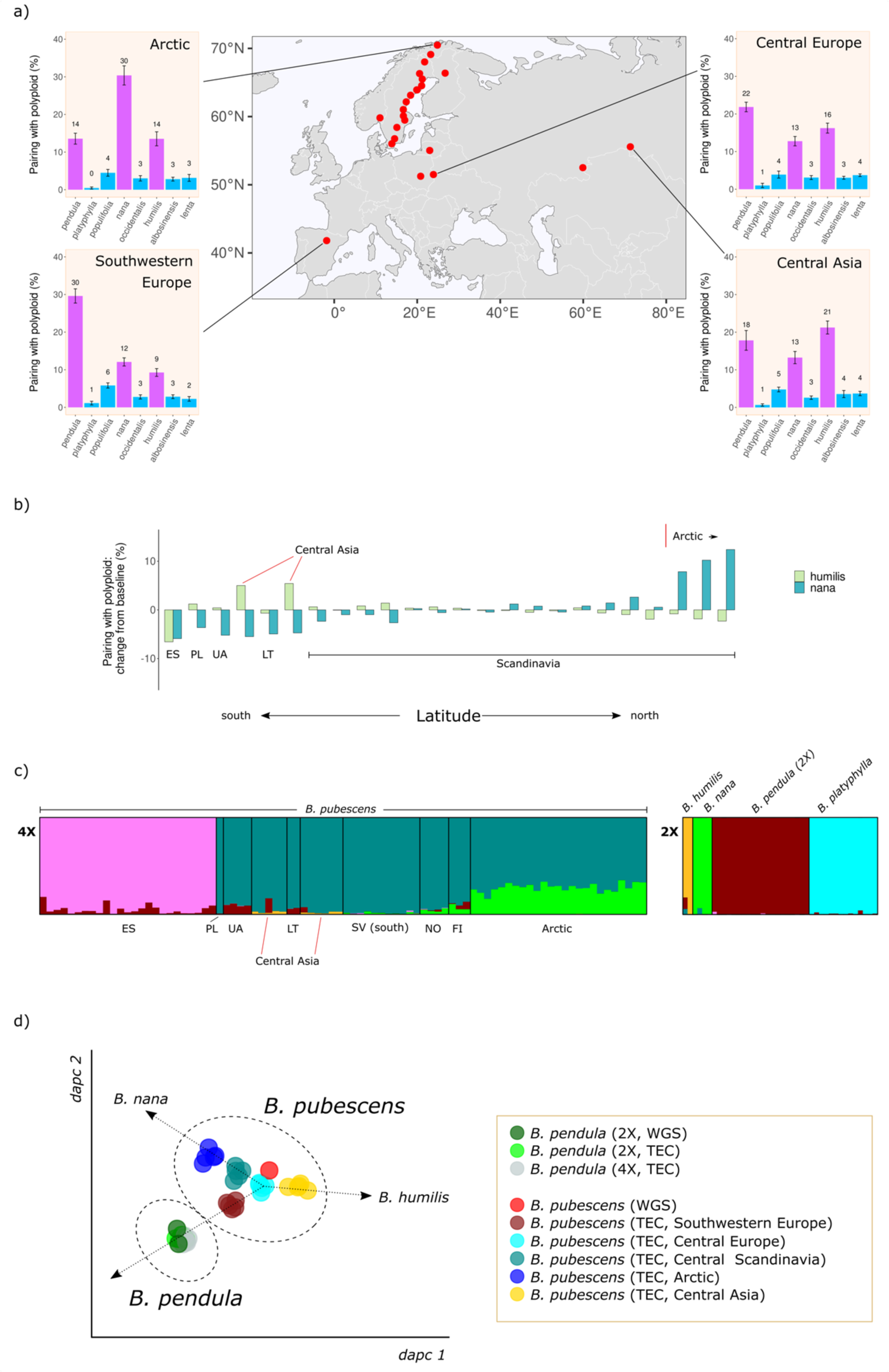
*B. pubescens* genomic composition and population structure –. **(a)** Barplots show the frequency with which the polarized *B. pubescens* sequence pairs with other birch species during phylogenetic inference at four different locations across the *B. pubescens* species range, based on phylogenetic analysis of individual gene families using IQ-TREE2. *B. platyphylla* was used as the reference sequence during genomic polarization. Red dots indicate *B. pubescens* sampling locations. **(b)** Variation in polyploid pairing frequencies between polarized *B. pubescens* and *B. humilis* (light green) or *B. nana* (dark green) measured across sampling locations, compared to the baseline value, the average pairing frequency across all locations. *B. platyphylla* was used as the reference sequence during genomic polarization. **(c)** Estimated genetic admixture of 126 individuals belonging to five *Betula* species according to STRUCTURE analysis based on 10,000 variant loci and K=6. Each individual is represented by a vertical column. Data for diploid (2X) and tetraploid samples (4X) is shown separately to facilitate interpretation. Data for tetraploid samples is sorted by latitude from left (south) to right (north). Population identifiers (based on Supplementary Figure S1): ES (site 1), PL (2), UA (3), CA-west (22), LT (4), CA-east (23), SV-south (7), NO (5), FI (21), Arctic (20). The number of optimal clusters, as determined using the ln Pr(X|K) method, is 6 (Supplementary Figure S10). **(d)** Discriminant analysis of principal components (DAPC) plot for 49 birch accessions based on 50,000 variant loci. *B. pendula* and *B. pubescens* are represented by both whole genome sequencing (WGS) and targeted exome capture (TEC) accessions. *B. pubescens* groups are as follows: Southwestern Europe (site 1.), Central Europe (2, 3, 4), Central Scandinavia (17), Arctic (20), Central Asia (22, 23).

The nana genomic component is highest in Arctic regions, decreasing progressively as we move towards lower latitudes (Fig. 2b and Supplementary Fig. S7). The humilis genomic component is more uniform across the *B. pubescens* range, with a maximum value in Central Asia, and below average values in Southern Europe (Fig. 2b and Supplementary Fig. S7). While the nana and humilis components are both reduced in specimens collected in Southern Europe (Fig. 2b and Supplementary Fig. S7d), they are nonetheless still clearly observed, unlike in tetraploid *B. pendula* samples where their presence is vestigial and mostly in line with background ILS levels (Supplementary Fig. S8). This suggests that the Southern European population, while unique among the populations included in this study, retains genomic characteristics generally found in other *B. pubescens* populations. Using the *B. pendula* chromosome map as a reference, we also checked whether *B. pubescens* loci containing only alleles of *B. pendula* origin varied across samples and whether they were concentrated in specific genomic regions. The location of *B. pubescens* loci fixed for the pendula allele varied from sample to sample and were generally distributed across the whole genome (*Supplementary Materials 4*).

Complementing the results presented above, we also performed the analysis of a subset of *B. pubescens* specimens using STRUCTURE, an approach used in previous studies to evaluate the level of admixture between this polyploid and other birch species (Wang et al. 2014; Eidesen et al. 2015; Zohren et al. 2016; Tsuda et al. 2017). The STRUCTURE analysis differentiates different *B. pubescens* populations (Fig. 2c and Supplementary Figs. S9 and S10) but, unlike the phylogenetic analysis based on polarized genomic sequences (Figure 2a and Supplementary Fig. S7), it provides no hint that all *B. pubescens* specimens across the species range contain large genomic components of *B. nana* and *B. humilis* origin. Population structure was also investigated using DAPC. In the resulting plot, individual *B. pubescens* populations are distributed along a tri-axis (Figs. 2d and Supplementary Fig. S11), with the Scandinavian and Arctic populations showing a heightened nana contribution, the Central Asia population being shifted towards *B. humilis*, and the Southwestern European samples being closer to the *B. pendula* cluster.

### Polyploidization and Hybridization Modeling Gives Support to an Autopolyploid Origin for B. Pubescens in the Absence of Homoeologous Exchange

We appraised the ability of nine hypothetical *B. pubescens* polyploidization models (Supplementary Fig. S6) to explain observed polyploid pairing patterns across the species’ range (shown in Figure 2a and Supplementary Figure S7) using an approximate Bayesian computation rejection algorithm (Supplementary Fig. S3). We proceeded in two steps. First, we excluded the possibility of full homoeologous replacement in any of the allopolyploid models, namely that gene copies initially found in one of the subgenomes could attain fixation. We then run a second set of simulations where we allowed for homoeologous replacement.

Upon comparison with the observed data for a *B. pubescens* population in Southern Sweden (Figs. 3 and Supplementary Fig. S12), the best fit (weighted *L2* norm distance = 0.49 ± 0.01) was obtained when *B. pubescens* was assumed to be an autopolyploid which originated via WGD from the common ancestor of *B. pendula* and its sister species, *B. platyphyll*a (Model 2: AAAA). Optimized levels of introgression for Model 2 were *h_nana_* = 0.41 ± 0.03 and *h_humilis_* = 0.34 ± 0.02, suggesting that, post-polyploidization, gene flow from *B. nana* and *B. humilis* have replaced at least one gene copy in 41% and 34% of all loci, respectively, in *B. pubescens* specimens in Southern Sweden. Levels of introgression are predicted to be even higher for *B. pendula*, whose alleles are present on average in 79% of all loci in the *B. pubescens* genome.

**Figure 3.**
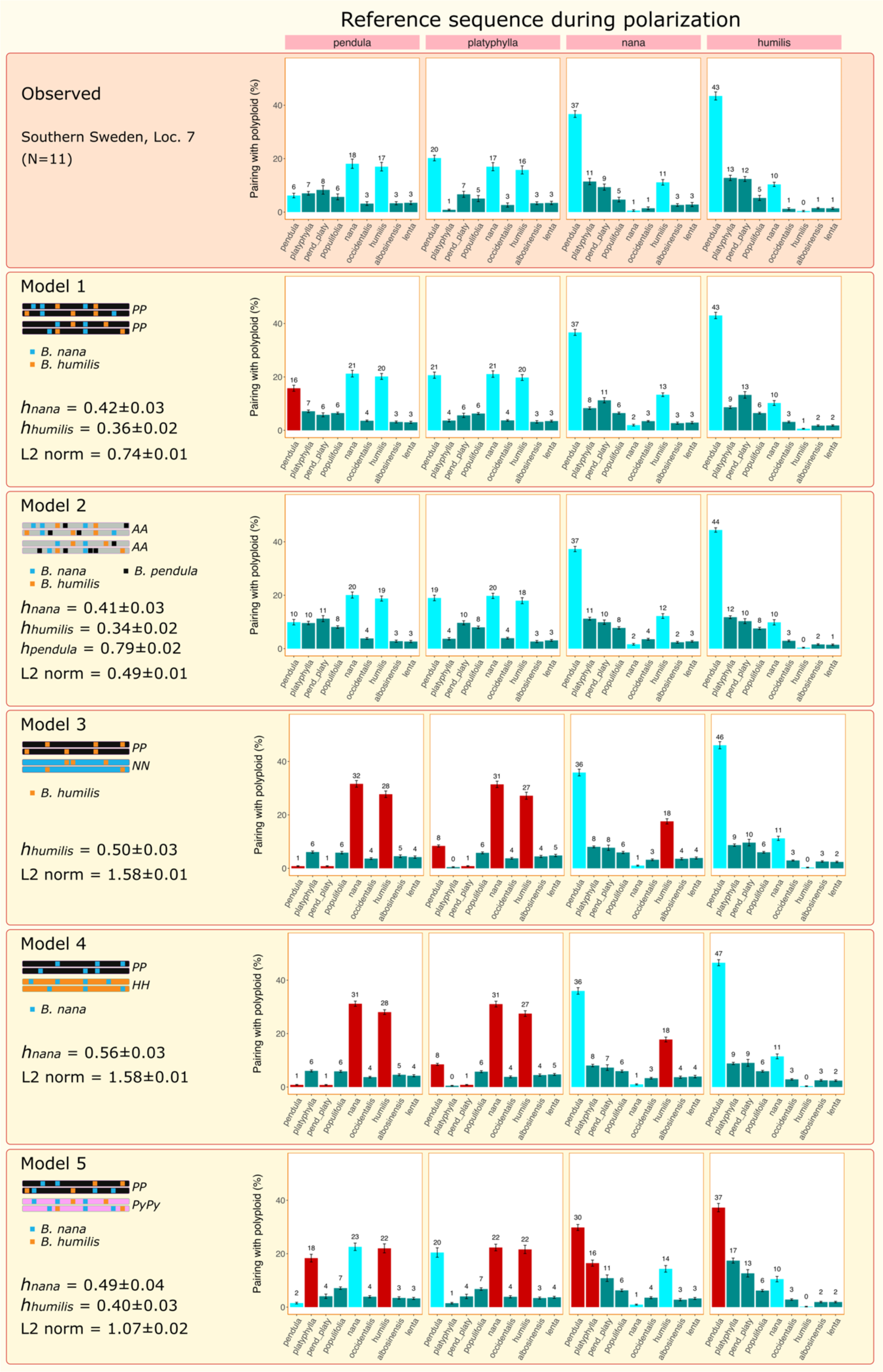
Comparison of *B. pubescens* evolutionary models. Barplots shows the frequency with which the polarized *B. pubescens* sequence pairs with other birch species, for different evolutionary models and polarization settings, based on phylogenetic analysis of individual gene families using IQ-TREE2. P, N, H, P_y_, and A indicate the hypothetical evolutionary origin of each subgenome component in the tetraploid (*B. pendula*, *B. nana*, *B. humilis*, *B. platyphylla*, and the common ancestor of *B. pendula* and *B. platyphylla*, respectively). Modeling data summarizes optimized results [top 5% simulation runs (50 out of 1000 runs)], obtained using approximate Bayesian computation using the observed values in a population from Southern Sweden as a reference (top row). Each simulation run averages results over 25 independent replicates. Cyan bars indicate level of polyploid-pairing for the three key species (see text for details). Bars in red indicate discrepancies between predicted and experimental results higher than 5 percentage points. The weighted *L2* norm distance measures the goodness of fit and includes data for all four polarizations (lower *L2* values denote a better fit). *L2* values are based on extended pairing spectra shown in Supplementary Figure S12. *pend_platy* indicates instances when the polarized polyploid sequence is an outgroup to the pendula-platyphylla clade. See Supplementary Figure S12 for modeling results for Models 6-9.

The pendula autopolyploid model (Model 1: PPPP) provided the second-best overall fit (weighted *L2* norm distance = 0.74 ± 0.01). The deterioration in the *L2* error observed when using this model was due to its inability to properly predict the pendula genomic contribution, which was inflated for one of the polarization geometries (peak highlighted in red in Figures 3). All the allopolyploid models provided much worse fits, with predicted polyploid-pairing profiles for the three key species (*B. pendula*, *B. nana*, and *B. humilis*) showing major discrepancies when contrasted against the observed data (Figs. 3, 4a, and 4b). Unlike the autopolyploidization models, where the nana and humilis genomic contributions can be adjusted independently, in the *B. pendula*/*B. nana* and *B. pendula*/*B. humilis* allopolyploid models the phylogenetic signals from nana and humilis genomic components show an inversely proportional dependency: a progressive reduction in the *B. nana* peak was observed as *h_humilis_* values are increased, and vice-versa (dashed line in Fig. 4a). A similar dynamic was observed in the remaining allopolyploid models.

**Figure 4.**
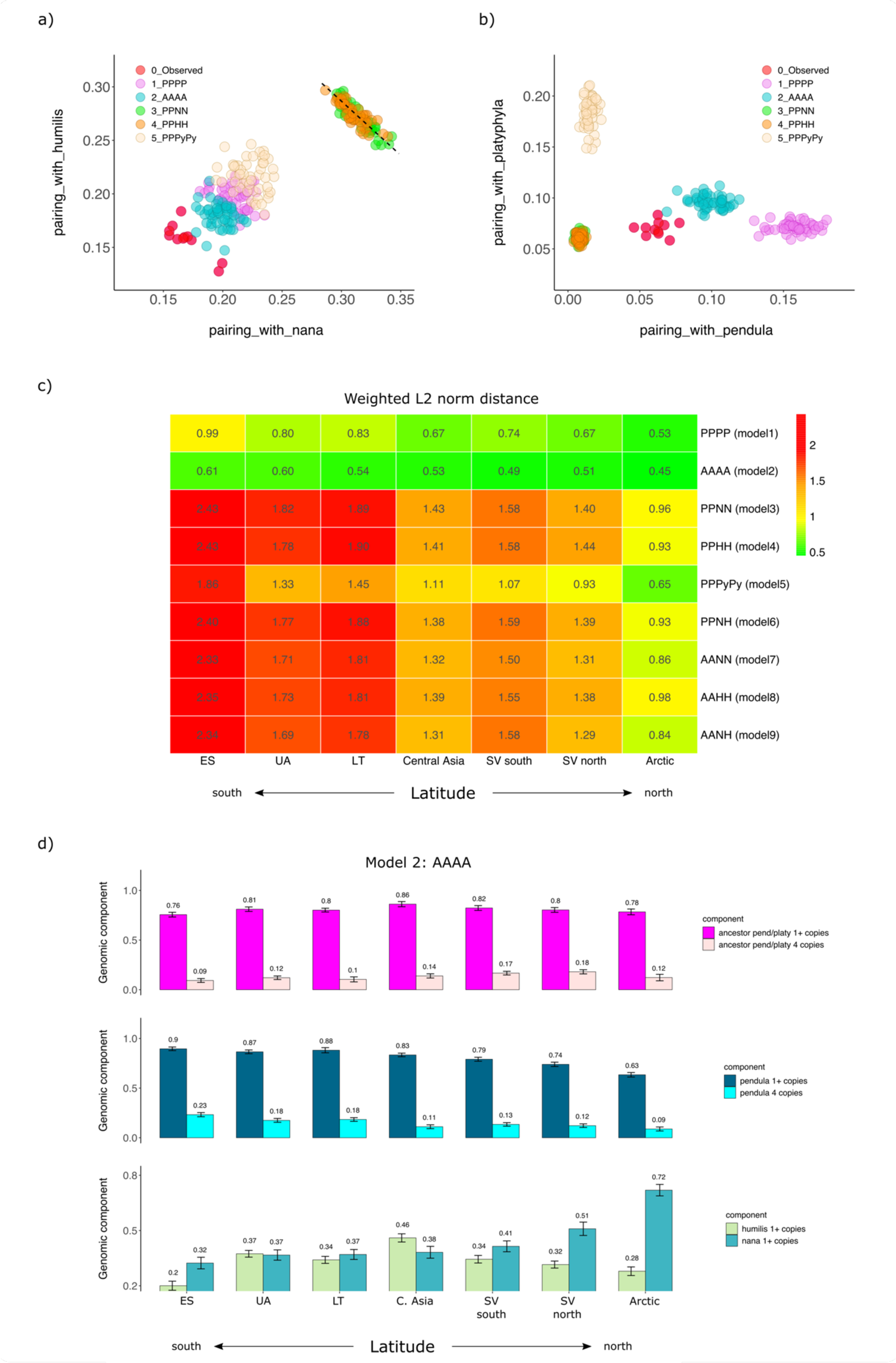
*B. pubescens* evolutionary models’ predictive power. Frequency with which the polarized *B. pubescens* sequence pairs with **(a)** *B. nana* and *B. humilis* and **(b)** *B. pendula* and *B. platyphylla*, as predicted by different evolutionary models, based on phylogenetic analysis of individual gene families using IQ-TREE2. Values observed experimentally for a population in Southern Sweden are shown in red. For each model, data is shown for the top 5% simulation runs with lowest L2 distance. Values are based on phylogenies obtained when *B. platyphylla* (a) or *B. pendula* (b) was used as the reference sequence during polarization. Each simulated data point summarizes runs for 25 independent replicates. The dashed line is a guide to the eye. **(c)** Weighted *L2* norm distance, averaged over the top 5% simulation run, for different models and sampling locations. Lower *L2* values denote a better fit. *B. pubescens* population groups are as follows (based on Supplementary Figure S1): ES (site 1), UA (3), LT (4), Central Asia (23), SV south (7), SV North (17), and Arctic (20). (**d**) *B. pubescens* genomic composition as a function of sampling location estimated using the AAAA autopolyploid model. “1+ copies”: percentage of loci with at least one gene copy originating from the common ancestor of *B. pendula* and *B. platyphylla* (top), *B. pendula* (center), *B. humilis* or *B. nana* (bottom). “4 copies”: percentage of loci where all four gene copies originate from the common ancestor of *B. pendula* and *B. platyphylla* (top), or *B. pendula* (center).

The AAAA autopolyploid model outperforms all other models not only when applied to the Southern Sweden *B. pubescens* population, but also in all other populations across the species range (Fig. 4c). Polyploid pairing predictions based on the AAAA autopolyploid model for seven geographic locations covering the *B. pubescens* range reveal a high degree of accuracy when compared with observed values (Supplementary Fig. S13). The *B. pubescens* genomic composition estimated using the AAAA model suggests that, depending on geographic location, 63 to 90% of all loci contain at least one gene copy of *B. pendula* origin, and that in 9 to 23% of the genome all four gene copies are of *B. pendula* ancestry (Fig. 4d). Rampant gene flow from parental diploids has been previously observed in other diploid-autotetraploid complexes such as in *A. arenosa* (Arnold et al. 2015, Monnahan et al. 2019). The simulation results also confirmed that the nana genomic component follows a latitude cline (Fig. 4d), while the percentage of loci in *B. pubescens* harboring alleles of *B. humilis* origin is relatively stable along the species range and reaches its highest value in Central Asia (Fig. 4d). In spite of the high levels of gene flow from multiple birch species, modeling results suggest that the *B. pubescens* genome retains a large fraction of ancestral alleles, with 76 to 86% of all loci containing alleles inherited from the common ancestor of *B. pendula* and *B. platyphylla* (Fig. 4d).

### Allotetraploid Models that Include Homoeologous Exchange All Converge Towards the Autopolyploid Hypothesis

To establish whether the presence of homoeologous replacement could allow any of the allopolyploid models to match or surpass the autopolyploid models in their ability to fit the data, we ran a new set of simulations where homoeologous replacement was permitted. When homoeologous replacement was allowed, all seven allopolyploid models converged towards genomic configurations similar to the ones obtained for the autopolyploid models (Supplementary Table S2). In Models 3-6, the non-pendula subgenome was greatly abridged, first due to homoeologous replacement by the pendula subgenome and, in Models 3-5, by further introgression with a third species. A similar pattern was observed in allopolyploid models where one of the two subgenomes originated from the common ancestor of *B. pendula* and *B. platyphylla* (Models 7-9). Under this hypothetical scenario, the extent of homoeologous replacement was estimated to affect between 29% (PPPyPy model) and 39% (PPNH model) of the *B. pubescens* genome. Predicted levels of homoeologous replacement by the nana, humilis, or platyphylla subgenomes in allopolyploid models were only vestigial and never exceed 3% (Supplementary Table S2).

### Dating of B. pubescens origin

To date the origin of *B. pubescens*, we used fastsimcoal2 to fit demographic models to SNP data for {*B. pubescens*, *B. pendula*, *B. platyphylla*} population trios. The analysis was carried out separately for four different *B. pubescens* populations in Central Europe, Southern Sweden, Central Asia, and Southwestern Europe. We started by testing simple evolutionary models based on all possible phylogenetic relationships among the three species included. Model comparison based on Akaike weights unanimously identified the ((*B. pendula*, *B. platyphylla*), *B. pubescens*) model as the most likely phylogeny (Supplementary Table S3), providing further evidence that *B. pubescens* first emerged before *B. pendula* and *B. platyphylla* diverged and evolved as distinct species. Next, we optimized the best phylogenetic topology by testing all possible migration combinations (129 models). As before, this analysis was done independently for different *B. pubescens* populations and Akaike weights were used to identify the model that best fits the data. The same exact migration model was deemed to best fit the data independently of the dataset used (*Supplementary Materials 5*), with modeling based on the Central European *B. pubescens* population providing the best overall fit (highest estimated maximum likelihood). Values estimated based on this dataset indicate that *B. pendula* and *B. platyphylla* diverged about 100,000 generations ago (Fig. 5), that is, about 2.5 Ma based on a 25-year generation time, in agreement with previous estimates (2.6 Ma; Chen et al. 2021). Gene flow from *B. pendula* into *B. pubescens* was estimated to be at least one order of magnitude higher than gene flow from *B. platyphylla* (Fig. 5). *B. pubescens* was estimated to have diverged from the common ancestor of *B. pendula* and *B. platyphylla* about 180,000 generations ago (Fig. 5), with parametric bootstrapping indicating a 95% confidence interval between 178,659 and 188,529 generations (Supplementary Table S4).

**Figure 5.**
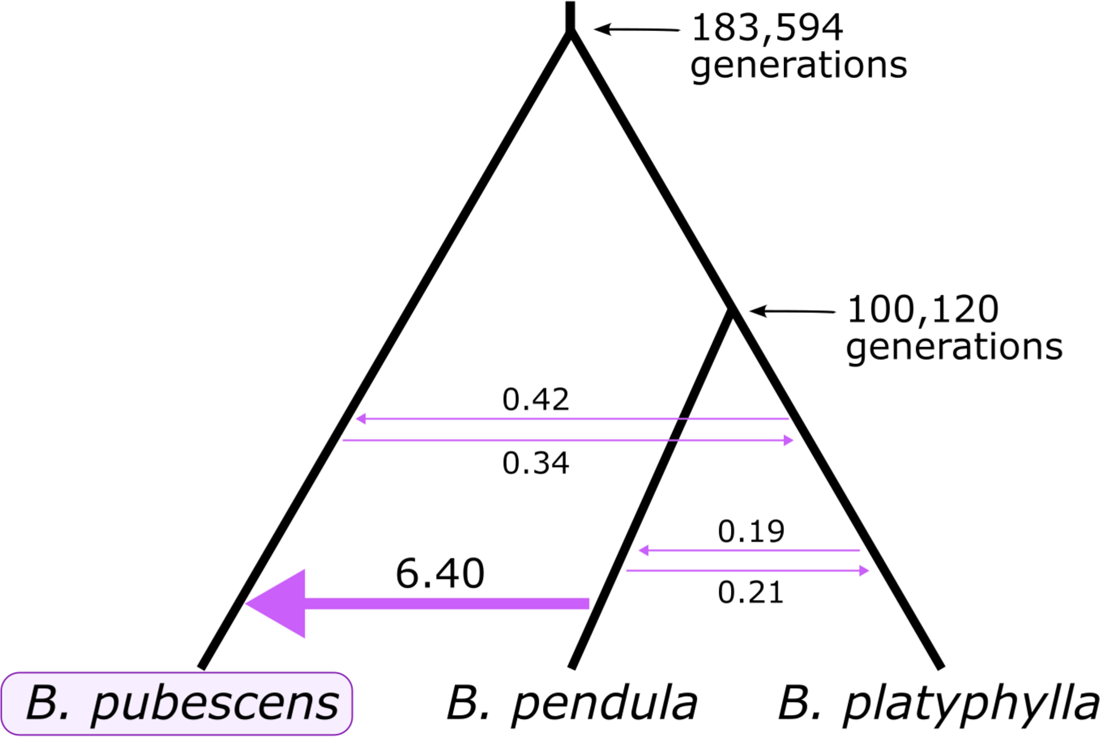
Phylogenetic topology and maximum likelihood estimates (MLE) of model parameters obtained using fastsimcoal2. Mean divergence times (in generations) and number of introgression events (per generation) estimated for migration model with highest Akaike weight. Only *B. pubescens* loci containing four gene copies of *B. pendula* and/or *B. platyphylla* ancestry were included in the analysis (loci containing one or more gene copies originating from *B. nana* or *B. humilis* were excluded).

### Identification and Functional Analysis of introgressed alleles of B. Nana and B. Humilis Ancestry

The geographic distribution and potential biological role of alleles of *B. nana* and *B. humilis* origin was examined separately for each gene. Of the 524 loci included in this study whose associated MSAs contain at least 100 parsimony-informative sites, 425 different genes were tagged as having an allele of *B. nana* ancestry in at least one *B. pubescens* individual (457 and 470 genes were tagged as containing alleles of *B. humilis* and *B. pendula* origin, respectively). For many of these genes, the fraction of individuals within a population possessing introgressed alleles remained in the low single digits across the *B. pubescens* geographic range (Fig. 6 and *Supplementary Materials 6*). On the other hand, there were 105 genes where the fraction of individuals with an allele of *B. nana* parentage reached 65% or more in at least one population (’highly retained alleles’ hereafter). There were 61 and 290 genes with highly retained alleles of *B. humilis* and *B. pendula* ancestry, respectively. Among the loci studied only one contained solely alleles inherited from the common ancestor of *B. pendula* and *B. platyphylla* in all the individuals sampled, and only two loci contained exclusively alleles of *B. pendula* origin (Supplementary Table S5). Many other loci contain alleles originating from *B. pendula* (*Supplementary Materials 6*), which have progressively replaced those of ancestral origin, but it is unclear whether their pervasiveness is due to selective pressure or simply a consequence of incessant gene flow between two species with largely overlapping geographic ranges.

**Figure 6.**
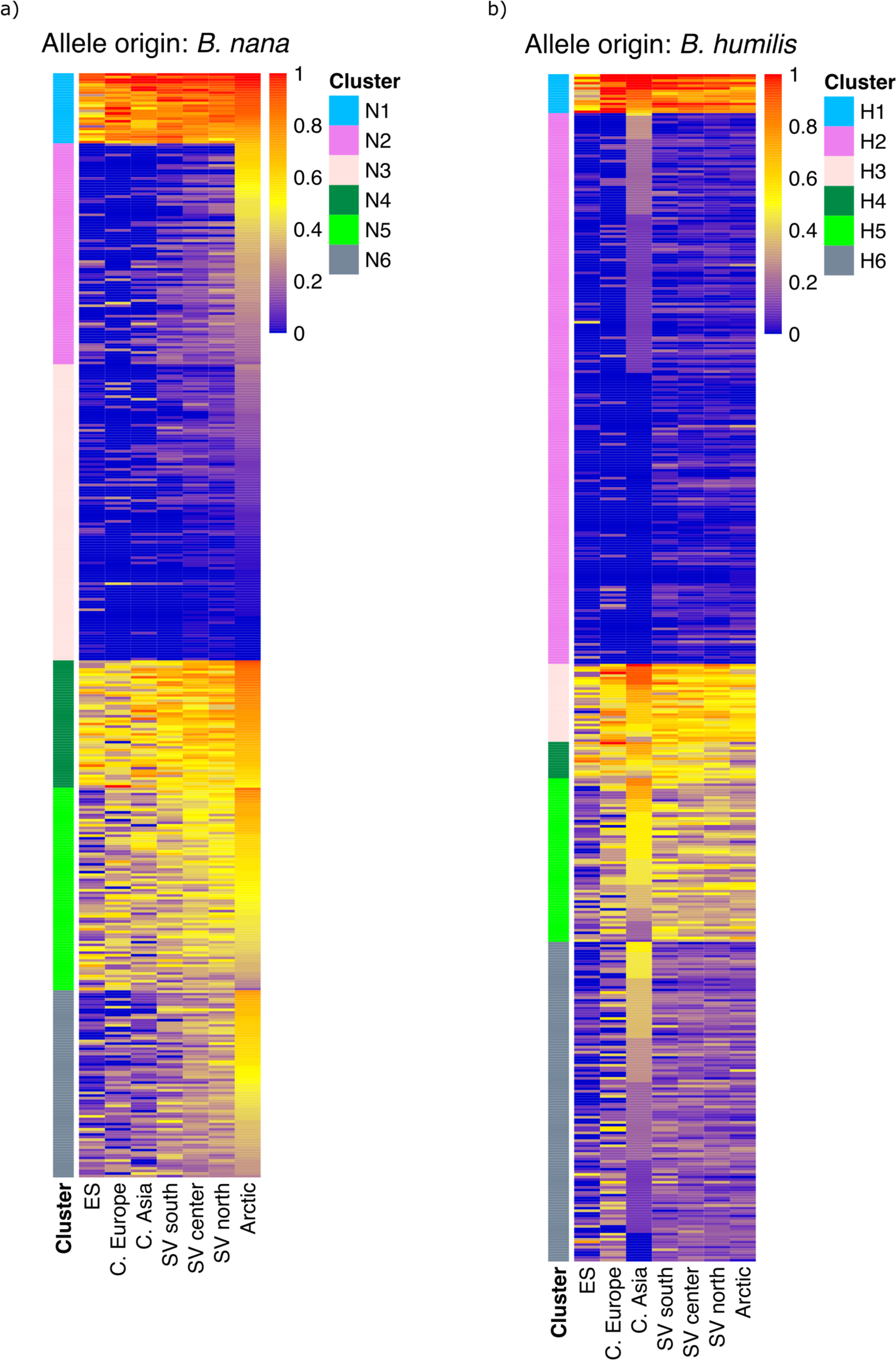
Abundance profiles for *B. pubescens* allelic variants of *B. nana* and *B. humilis* origin. Heatmaps show fraction of *B. pubescens* individuals in different populations (columns) containing an allele of **(a)** *B. nana* or **(b)** *B. humilis* origin, for different genes (rows). K-means cluster analysis was used to partition genes in groups with similar abundance profiles across sampling locations. The analysis identified six clusters (N1 to N6) for alleles of *B. nana* origin, and six clusters (H1 to H6) for alleles of *B. humilis* origin.

Gene flow from *B. nana* and *B. humilis* into *B. pubescen*s is generally more geographically localized. Alleles of *B. nana* or *B. humilis* origin preserved across the *B. pubescens* geographic range are therefore of particular interest as they are likely candidates for gene variants that provide a fitness advantage. As expected, geographic distributions of introgressed *B. nana* and *B. humilis* alleles vary depending on the species of origin. Many highly-retained alleles of *B. nana* ancestry show a clinal or quasi-clinal dependency with the highest prevalence being observed in Arctic regions, but there is a second group of genes where nana alleles are found in most individuals in most populations (Figs. 6a, 7a and 7c). As for alleles of *B. humilis* origin, while the largest group consists of alleles mostly found in Central Asia, one can find a sizable number of alleles highly retained across all populations, as well as some showing a clinal dependency (Figs. 6b, 7b and 7c).

**Figure 7.**
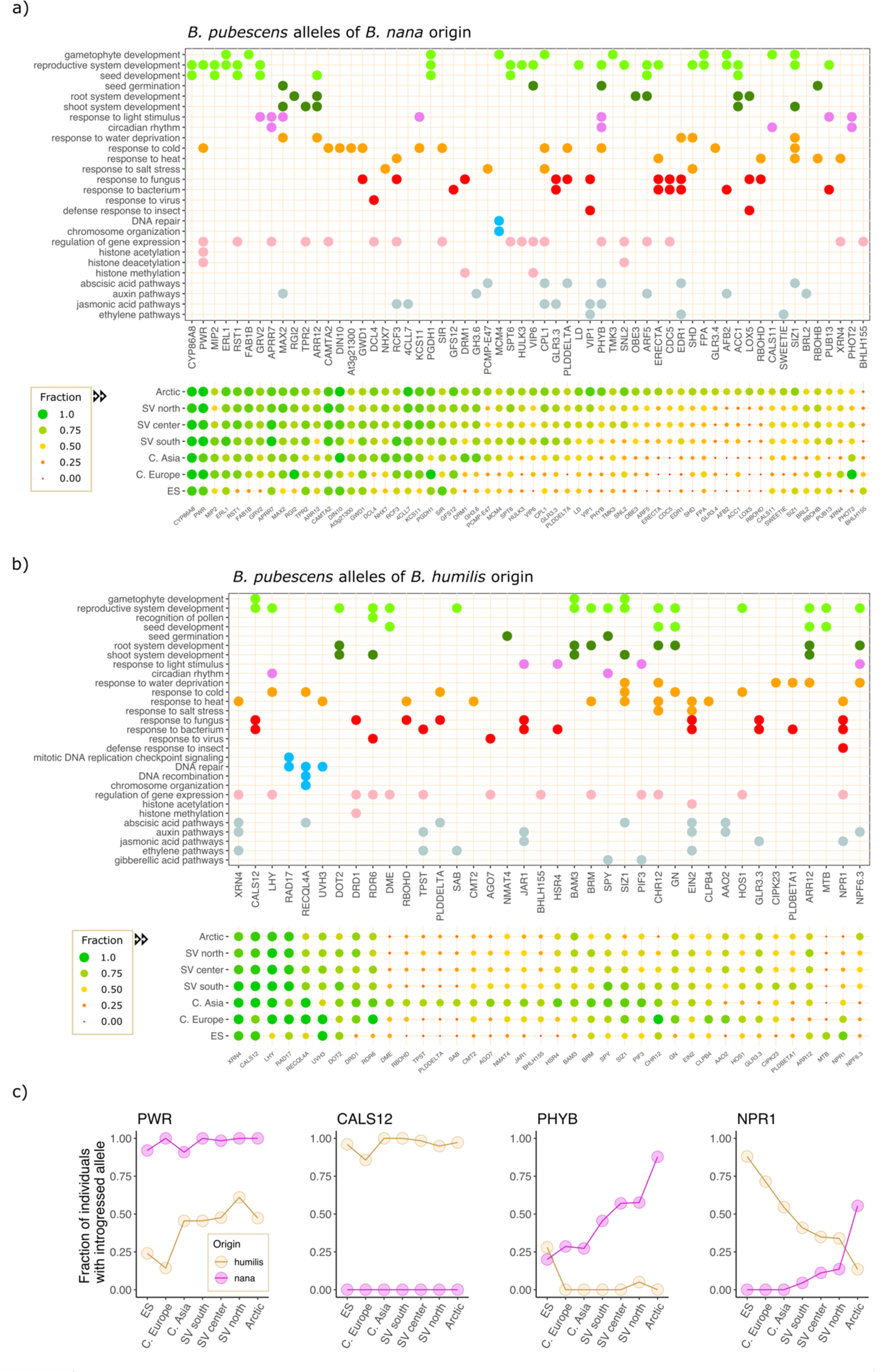
Functional analysis of *B. pubescens* allelic variants of *B. nana* and *B. humilis* origin. Gene ontology (GO) annotation of genes associated to retained allelic variants of **(a)** *B. nana* and **(b)** *B. humilis* origin. Symbol colors represent broad GO categories (e.g. biotic defense responses are shown in red). The fraction of individuals in a population carrying an introgressed allele are also shown for each gene and geographic location. Data are shown only for genes with a nana and/or humilis allelic fraction ≥ 0.65 observed in at least one population and for which a GO annotation is available. **(c)** Fraction of individuals with introgressed alleles observed across different populations for homologs of the following *Arabidopsis* genes: *POWERDRESS* (*PWR*), *CALLOSE SYNTHASE 12* (*CALS12*), *Phytochrome B* (*PhyB*), and *REGULATORY PROTEIN NPR1* (*NPR1*). *B. pubescens* population identifiers (based on Supplementary Figure S1): Arctic (sites18, 19, 20); SV north (14, 15, 16, 17); SV center (9, 10, 11, 12, 13); SV south (6, 7, 8); Central Asia (22, 23); Central Europe (2, 3, 4), ES (1).

A more striking difference between alleles of *B. nana* and *B. humilis* ancestry concerns their purported functional role. Many of the loci containing highly preserved alleles derived from *B. nana* are believed to code for genes involved in response to cold and/or light stimulus in plants, including homologs of *PWR*, *PRR7*, *MAX2*, *CAMT2*, *DIN10*, *GWD1*, and *KCS11* (Fig. 7a and Table 1). Additionally, an allele of *B. nana* origin in *Phytochrome B* (*PhyB*), the primary red-light photoreceptor for circadian control in *Arabidopsis* and which also functions as a temperature sensor (Somers et al. 1998; Jung et al. 2016; Legris et al. 2016), shows a marked clinal distribution, with its prevalence varying from 20% in Southern Europe to 88% in the Arctic (Fig. 7c). Among highly-retained alleles of *B. humilis* ancestry, we found a gene known to be involved in gametophytic fertility in plants, *CALS12* (Fig. 7b and Table 1). A humilis *CALS12* allele was detected in 97.0% of all *B. pubescens* specimens sampled (Fig. 7c), and no other allele either of *B. pendula* or *B. nana* origin was detected for this locus in any of the samples studied, suggesting that this gene is under strong selective pressure. Highly preserved alleles obtained from *B. humilis* are also found within three genes involved in DNA repair and maintenance of meiotic stability: *RECQL4A*, *UVH3*, and *RAD17* (Fig. 7b and Table 1).

**Table 1.** Selected *B. pubescens* loci containing highly retained alleles of *B. nana* or *B. humilis* origin.

| Gene / protein<br>(*A. thaliana*<br>nomenclature) | Overall prevalence<br>(% of all *B. pubescens*<br>specimens sampled) | Function |
| --- | --- | --- |
| **Selected alleles of *B. nana* origin** |  |  |
| CYP86A8<br>(Cytochrome P450) | 99.3 | Involved in synthesis of lipid-polyester used in the seed coat to prevent embryo aging and ensure seed longevity in *A. thaliana* (Renard et al. 2020) |
| PWR<br>(Powerdress) | 98.5 | Co-regulates both flowering time and leaf senescence in *A. thaliana* (Chen et al. 2016; Kim et al. 2016); part of a histone-modifying complex that epigenetically controls freezing tolerance (Lim et al. 2020). |
| CAMTA2<br>(Calmodulin-binding<br>transcription activator 2) | 92.9 | Regulation of cellular response to cold in Arabidopsis (Kim et al. 2013). |
| KCS11<br>(3-ketoacyl-CoA synthase 11) | 85.5 | Regulation of cellular response to cold in Arabidopsis (Joubès et al. 2008). |
| APRR7<br>(Two-component response<br>regulator-like APRR7) | 91.4 | Involved in phasing of the circadian clock and control of seedling deetiolation (Kaczorowski et al. 2003). |
| MAX2<br>(F-box protein MAX2) | 83.3 | Positive regulator of photomorphogenesis (Shen et al. 2007). |
| DIN10<br>(Dark inducible 10) | 93.7 | Regulation of cellular response to cold in Arabidopsis (Maruyama et al. 2009). |
| GWD1<br>(Starch excess 1) | 78.1 | Regulation of cellular response to cold in Arabidopsis (Yano et al. 2005). |
| PhyB<br>(Phytochrome B) | Clinal dependency;<br>highly retained in Arctic<br>population (88%) | Primary red-light photoreceptor for circadian control in Arabidopsis (Somers et al. 1998), functions as a temperature sensor (Kim et al. 2002, Jung et al. 2016; Legris et al. 2016), and regulates flowering time and seed germination (Seo et al. 2006; Hajdu et al. 2015). |
| **Selected alleles of *B. humilis* origin** |  |  |
| CALS12<br>(Callose synthase 12) | 97.0 | Plays role in pollen development and fertility in Arabidopsis, being involved in the synthesis of the callose wall separating microspores after meiosis (Enns et al. 2005). |
| XRN4<br>(Exoribonuclease4) | 94.4 | Part of a mRNA decay pathway involved in plant acclimation to heat stress (Merret et al. 2013) |
| RECQL4A<br>(ATP-dependent DNA<br>helicase Q-like 4A) | 74.5<br>(100% in Central Asia and<br>Central Europe) | Contributes to the maintenance of genomic stability in *A. thaliana* by modulating the response to DNA damage, suppressing spontaneous somatic crossovers, and limiting the number of crossovers during meiotic recombination (Bagherieh-Najjar et al. 2005; Hartung et al. 2008; Séguéla-Arnaud et al. 2015). Is also required for dissolving telomeric bridges between non-homologous chromosomes during meiosis (Higgins et al. 2011). |
| RAD17<br>(Cell cycle checkpoint protein) | 95.5 | Encodes a checkpoint protein that is part of a DNA damage response pathway induced by genotoxic stress in yeast, mammals, and Arabidopsis (Zhou and Elledge 2000; Bao et al. 2001; Heitzeberg et al. 2004). |
| UVH3<br>(DNA repair protein) | 81.7<br>(100% in Central and<br>Southwestern Europe) | Encodes an endonuclease involved in the repair of UV-damaged DNA in Arabidopsis (Liu et al. 2001). |
| LHY<br>(Late elongated hypocotyl) | 95.2 | Circadian clock gene involved in the regulation of flowering time and freezing tolerance in Arabidopsis (LHY; Fujiwara et al. 2008; Dong et al. 2011). In *Picea abies* specimens sampled in Sweden, there is a strong clinal preference for different LHY alleles which matches changes in environmental conditions (Li et al. 2022). |
| NPR1<br>(Nonexpresser of PR genes 1) | Clinal dependency;<br>highly retained in Southern<br>European population (88%) | Salicylic acid (SA) transducer and a key regulatory gene of the systemic acquired resistance (SAR) pathway activated in plants upon local infection by microbial pathogens (Cao et al. 1994). There is evidence of balancing selection acting on NPR1 to maintain two allelic classes in *A. thaliana* (Caldwell and Michelmore 2009) and in *Capsella grandiflora* (Sicard et al. 2015), likely due to functional differences between alleles. |

## Discussion

### Evidence for an Autopolyploid Origin for B. pubescens

Here, we present a Bayesian framework that can be used to establish the mode of origin of a complex polyploid by disentangling the two key evolutionary mechanisms at play, clarifying whether alleles of diverse origin have been acquired through polyploidization, introgression, or both. Our results suggest that *B. pubescens* is most likely the offspring of a single parental species, the common ancestor of *B. pendula* and *B. platyphylla*, which gave rise to it via WGD, with the newly formed autopolyploid having subsequently undergone extensive introgressive hybridization with *B. pendula*, *B. nana* and *B. humilis*. This assertion is supported by comparisons of real and simulated polyploid frequency pattern data, with the autopolyploidization model providing the best overall goodness-of-fit and closest match to the observed polyploid-paring patterns across all populations. Demographic modeling using fastsimcoal2 also provided strong support for a phylogeny where *B. pubescens* speciation occurred prior to *B. pendula* and *B. platyphylla* evolved as separate species. Pervasive introgression from *B. pendula*, *B. nana* and *B. humilis* would provide a plausible explanation for the high levels of allelic richness and genetic diversity observed in *B. pubescens*, which until now had been interpreted as indirect evidence of allopolyploidization.

Attempts to model the origin of *B. pubescens* based on a bi-parental allopolyploidization model produced contrasting results depending on whether homoeologous replacement was allowed to occur or not. In the absence of homoeologous replacement, allopolyploid models invariably produced scenarios that failed to replicate the polyploid-pairing patterns observed experimentally, leading to a degradation in *L2* values, as they all struggled to accurately predict the genomic contribution of a hypothetical second parental species. When homoeologous replacement was allowed to occur between subgenomes, the performance of allopolyploid models was in line with their autoploid counterparts, but the level of structural dominance required (affecting up to ∼39% of the *B. pubescens* genome) is unheard-of in established allopolyploids. This is discussed in more detail in the next section.

### A Hypothetical Allopolyploid Origin for B. pubescens Would Require Massive Levels of Homoeologous Replacement

Modeling results shown in this paper suggest that a hypothetical allopolyploid origin for *B. pubescens* would be possible − though unlikely − if it had been followed by subsequent homoeologous exchanges resulting in *B. pubescens* (or the ancestral *B. pubescens*/*B. platyphylla* allele) dominating both subgenomes in about 35% of all protein coding regions (with all gene copies from the second subgenome having been lost). While frequently observed in allopolyploid species, homoeologous replacement usually affects a relatively small fraction of genes in naturally formed polyploids. In the *Gossypium hirsutum* ‘TM-1’ cotton cultivar, an allotetraploid estimated to first have emerged 1.5 Ma, Li and colleagues (2015) identified 1,790 out of 35,056 genes (∼5%) present in the At subgenome that were transferred from the Dt subgenome. In the allotriploid ‘FenJiao’ banana cultivar (*Musa* ssp.), homoeologous exchanges from the A to the B subgenome were identified in 39 genomic segments containing 2,579 genes (out of 35,148 ∼ 7%; Wang et al. 2019). In *B. napus*, a recent (7,500-12,500 years BP; Chalhoub et al. 2014) allotetraploid and model organism for the study of genome restructuring and homoeologous exchange in neo-polyploids (Gaeta and Pires 2010; Chalhoub et al. 2014; Sun et al. 2017; Lloyd et al. 2018; Xiong et al. 2021), expression analysis of the ‘Yudal’ cultivar showed that 1,076 (2%) out of the ∼49,000 coding genes present in the C_n_ subgenome have been replaced by their homoeologous A_n_ subgenome duplicates (Lloyd et al. 2018). Similar levels of C_n_ to A_n_ homoeologous replacement (1,032 genes ∼ 2%) have been observed in the ‘ZS11’ *B. napus* cultivar (Sun et al. 2017). In the allotetraploid *Coffee arabica*, homoeologous loss was detected in less than 2% of 9,047 genes analyzed (Lashermes et al. 2016). Levels of homoeologous replacement in the neo-allotetraploid *A. suecica* (∼16,000 years BP; Novikova et al. 2017) are also rather subdued, involving at most a few hundred genes (Burns et al. 2021; Jiang et al. 2021), likely because the two parental species, *A. thaliana* and *A. arenosa*, are divergent enough (∼6 MYA) to prevent homoeologous exchange.

To our knowledge, rampant homoeologous replacement has only been observed in newly resynthesized allopolyploids (Chalhoub et al. 2014; Wu et al. 2021; Xiong et al. 2021) and in (very) recently formed natural allopolyploids such as the *Tragopogon miscellus* salsify, a biennial plant whose origin dates to the early 1900s after both diploid progenitors were introduced in North America from Europe (Chester et al. 2012). Here, we estimated that *B. pubescens* first emerged about 180,000 generation ago (4.5 Ma, assuming a 25-year generation time), therefore ruling out the possibility of it being a polyploid of recent origin. In short, while we cannot fully rule out an allopolyploid origin for *B. pubescens*, such a possibility seems very unlikely as it would require levels of homoeologous replacement that far exceed values so far observed in natural allotetraploid populations.

### Ecogeographical Variation in B. nana and B. humilis Introgression Levels

There is evidence that introgression into *B. pubescens* is both an ancestral and on-going phenomenon, occurring whenever the species involved coexist in the same geographic area. Evidence gathered from genetic, morphological and compatibility studies point to the presence of on-going gene flow from *B. humilis*, *B. nana* and *B. pendula* into *B. pubescens* in Western Eurasia and from *B. ermanii* in East Asia (Jeffers 1971; Anamthawat-Jónsson and Tomasson 1990; Atkinson 1992; Thórsson et al. 2001; Anamthawat-Jónsson and Thórsson 2003; Palmé et al. 2004; Jadwiszczak et al. 2012; Ashburner and McAllister 2013, pp. 68; Wang et al. 2014; Eidesen et al. 2015; Zohren et al. 2016; Tsuda et al. 2017). This is supported by our study showing that, although average introgression levels are very high across the species’ range, the identity of most introgressed alleles varies across individuals, even within the same local population, implying that many of them are likely being acquired stochastically via *de novo* introgression. Concurrently, we identified a set of alleles of *B. nana* and *B. humilis* origin whose presence is widespread across one or more *B. pubescens* populations. These highly retained introgressed alleles are located on different *B. pubescens* chromosomes, and may have been acquired during or soon after speciation, either because they themselves provided a selective advantage and hence were driven to high frequencies by selective sweeps, or alternatively because they are in linkage with other genes under selective pressure, or due to genetic drift acting in a small budding population.

Our results further suggest that, at the present time, *B. pubescens* populations located in two distinct zones, the Arctic and Central Asia, serve as the species’ major reservoirs for alleles of *B. nana* and *B. humilis* origin, respectively, arguably because these are locations where the species involved are abundant and live in sympatry (Ashburner and McAllister 2013, pp. 27-28). However, many of the birch species that hybridize with *B. pubescens* can also hybridize among themselves, as well as with several other birch species (Jeffers 1971; Atkinson 1992; Palmé et al. 2004; Jadwiszczak et al. 2012; Ashburner and McAllister 2013, pp. 62-71; Koropachinskii 2013; Tsuda et al. 2017), implying that each of them can act as a genetic bridge or conduit (McDonald et al. 2008; Grant and Grant 2020) for many of the others. It is therefore possible that some of the introgressed alleles of *B. nana* or *B. humilis* origin found today in *B. pubescens* might have been obtained via a bridge species, and not directly from *B. nana* or *B. humilis*, altogether sidestepping the sympatry requirement.

### B. pubescens Loci Containing Alleles of B. Nana Origin Are Involved In Climate Adaptation in Plants

One of the benefits of introgression is that it expands the pool of alleles available in a population upon which natural selection can act, often leading to phenotypic novelty and potentially facilitating adaptation to new environments (Rieseberg et al. 1999; Whitney et al. 2006; Dasmahapatra et al. 2012; Pardo-Diaz et al. 2012; Rius and Darling 2014; Arnold et al. 2016; Schmickl et al. 2017). Introgression of biotic resistance traits from a congener is believed to have enabled the southward range expansion of *Helianthus annuus*, a sunflower found in North America (Whitney et al. 2006). Interspecific genetic admixture has also been associated with a sudden increase in cold tolerance in the wasp spider *Argiope bruennichi* (Krehenwinkel and Tautz 2013), with the spread of the autotetraploid *A. arenosa* into more extreme habitats characterized by frequent drought and phytotoxic levels of metals (Arnold et al. 2016), and the expansion of the poplar *Populus trichocarpa* into colder and drier climates (Suarez-Gonzalez et al. 2018).

Here, we identified several regulatory genes associated to abiotic stress in plants that contained alleles of *B. nana* origin obtained via introgression. These include homologs of *PWR*, *CAMTA2*, *DIN10, GWD1*, and *KCS11.* While we can only speculate whether such derived alleles contribute to local adaptation and range expansion in *B. pubescens*, their retention by most individuals sampled across the species range suggests that they could be located in genomic regions under selective pressure, warranting further investigation. *PWR* is known to co-regulate both flowering time and leaf senescence in *A. thaliana* (Chen et al. 2016; Kim et al. 2016) and to be part of a histone-modifying complex that epigenetically controls freezing tolerance (Lim et al. 2020), while *CAMTA2, DIN10, GWD1*, and *KCS11* are all involved in the regulation of cellular response to cold in *Arabidopsis* (Yano et al. 2005; Joubès et al. 2008; Maruyama et al. 2009; Kim et al. 2013). In northern Eurasia, *B. pubescens* and *B. nana* have geographic ranges that extend further north than that of *B. pendula* (Atkinson 1992; de Groot et al. 1997), and the acquisition and retention of new alleles associated with cold acclimatization from *B. nana* could be one of the factors contributing to the cold tolerance of northern populations of *B. pubescens*.

We also found a cline in the frequency of the *PhyB* nana allele in *B. pubescens*, with the allele decreasing in frequency from 88% in the Arctic to 20% in Southwestern Europe. This gene is located in a region of low relative genomic diversity in *B. pendula*, which was associated with a selective sweep (Salojärvi et al. 2017). *PhyB* is known to modulate flowering time in *A. thaliana* (Hajdu et al. 2015) and has been shown to regulate bud set and seasonal growth in *Populus* (Ingvarsson et al. 2008; Ding et al. 2021). Changes in the *PhyB* amino acid sequence can lead to differential plant responses to light (Filiault et al. 2008), and it has been speculated that intraspecific diversification among phytochromes might allow a species to adjust its life cycle to different climates (Jung et al. 2016). The presence of introgressed allelic variants of *B. nana* origin showing a clinal dependency, while congruent with an adaptive role, could also be caused by the stochastic loss of neutral alleles diffusing away from the contact zone (Barton and Hewitt 1989), or by changes in the spatial range for any of the species involved (Excoffier et al. 2009, Zohren et al. 2016). While the high prevalence of the *PhyB* nana allele in Arctic regions is a strong indication that it is located in a genomic region under strong selective pressure, further studies are required to elucidate whether changes in allelic frequency for this and other introgressed alleles in birch are the outcome of selection, drift, or both.

### B. pubescens Loci Containing Alleles of B. Humilis Origin Are Known to Regulate Meiotic Stability and Pollen Viability in Plant Species

How new polyploids regulate meiotic chromosomal segregation is one of the key issues hinging over their long-term genomic stability and gamete viability (Comai 2005; Cifuentes et al. 2010; Bomblies et al. 2015; Lloyd and Bomblies 2016; Baduel et al. 2018; Morgan et al. 2021; Bomblies 2023). Recent results suggest that some plant species attain this competence by co-opting adaptive alleles obtained via introgression from one or more third species. There is evidence that bidirectional gene flow between the autotetraploids *A. arenosa* and *A. lyrata* provided the latter with a set of pre-adapted alleles in genes that regulate the synaptonemal complex between homologous chromosomes during meiosis, favoring a reduction in multivalent formation, and which are located in loci that had previously been found to be under strong selection in both species (Yant et al. 2013; Marburger et al. 2019; Seear et al. 2020). Likewise, in the allotetraploid *A. suecica*, which has *A. thaliana* and the autotetraploid *A. arenosa* as parental species, there is evidence that key meiotic genes harbor allelic variants acquired from diploid *A. arenosa* (Nibau et al. 2022).

In this paper, we identified two meiosis genes in the tetraploid *B. pubescens*, *RECQL4A* and *RAD17*, that contain alleles acquired from *B. humilis*. Additionally, we identified another locus associated to gametophyte development, *CALS12*, which appears to be under strong selection in *B. pubescens* as specimens sampled across the species range are all homozygous for the allele of *B. humilis* origin. In *Arabidopsis*, *CALS12* is part of a pathway that synthesizes the tetrad callose wall separating the four microspores during cytokinesis (Enns et al. 2005) and silencing or misregulation of this gene has been shown to negatively impact male fertility in both *Arabidopsis* and rice (*Oryza sativa*) mutant lines (Enns et al. 2005, Shi et al. 2015, Li et al. 2018b). In rice and maize, loss of *RAD17* in pollen mother cells often causes sterility as it leads to non-homologous chromosome entanglement and chromosomal fragmentation (Hu et al. 2018; Zhang et al. 2021). In *A. thaliana*, *RECQL4A* is part of one of the three anti-crossover pathways that strictly limits the number of crossovers during meiotic recombination (Séguéla-Arnaud et al. 2015; Serra et al. 2018), and is required for resolving and dissolving telomeric bridges between non-homologous chromosomes during meiosis (Higgins et al. 2011). Reduction in crossover rates helps to limit multivalent prevalence during meiosis, which may have implications for the long term viability of a species as high rates of multivalent formation are often associated with low fertility and aneuploidy in neo-autopolyploids (Ramsey and Schemske 2002; Bomblies et al. 2015; Lloyd and Bomblies 2016). Bivalent formation with random partner choice (tetrasomic inheritance) is prevalent in *B. pubescens* (Stern 1965; Brown and Al-Dawoody 1979), with the rate of multivalent formation reported to be lower than that observed in diploid *B. pendula* (Brown and Al-Dawoody 1979). While further research will be required to fully elucidate the specific impact of introgressed alleles on development and fitness, the results here presented raise the enticing possibility that the acquisition of new adaptive alleles from *B. humilis* assisted *B. pubescens* in decreasing rates of multivalent formation, improving meiotic stability and gametophytic fertility.

### Solving the Origin-Introgression Riddle in Polyploid Species

In this article, we used genomic polarization and a Bayesian framework to solve an old phylogenetics problem, namely, how to untangle polyploidization from introgression in order to determine the mode of origin and genomic composition of a complex polyploid. The results here presented also showcase the method’s ability to unearth the identity and provenience of introgressed alleles, which in conjugation with known information about their functional role, can be used to chart future research directions. The methodology here described can be easily extended to other polyploid species whose reticulate history remains debatable − often due to the confluence of polyploidization and introgression − in the birch (Wang et al. 2021), *Asteraceae* (Mandel et al. 2017), *Salix* (Wagner et al. 2020), *Senecio* (Kim et al. 2008), and *Cardamine* (Lihová et al. 2007) genera of flowering plants, among other. At the present time, the phasing approach here employed, genomic polarization, has only been used on tetraploids (Leal et al. 2023), and further testing and modeling might be required before it can be applied to higher ploidy species.

## Supplementary Materials

Supplementary scripts and data available from the Dryad Digital Repository: https://doi.org/10.5061/dryad.5tb2rbp9f and the GitHub repository https://github.com/LLN273/Complex-Polyploids. The genomic data underlying this article are available in The European Nucleotide Archive (ENA) at www.ebi.ac.uk and can be accessed with the PRJEB64873 [data held private until publication] and PRJEB14544 project accession codes. See Supplementary Materials 2 for ENA run accession codes for each sample.

## Acknowledgements

The computations were performed on resources provided by the Swedish National Infrastructure for Computing (SNIC) at the Uppsala Multidisciplinary Center for Advanced Computational Science (UPPMAX) under Project # SNIC 2017/7-149 and at the High Performance Computing Center North (HPC2N) under Project # SNIC 2020/9-84.

## References

Albertin W., Marullo P. 2012. Polyploidy in fungi: evolution after whole-genome duplication. Proc. R. Soc. B: Biol. Sci. 279:2497–2509.

Amborella Genome Project. 2013. The Amborella genome and the evolution of flowering plants. Science 342:1241089.

Anamthawat-Jónsson K., Tomasson T. 1990. Cytogenetics of hybrid introgression in Icelandic birch. Hereditas. 112:65–70.

Anamthawat-Jónsson K., Thór Thórsson A. 2003. Natural hybridisation in birch: triploid hybrids between *Betula nana* and *B. pubescens*. Plant Cell, Tissue Organ Cult. 75:99–107.

Arnold B., Kim S.-T., Bomblies K. 2015. Single Geographic Origin of a Widespread Autotetraploid *Arabidopsis arenosa* Lineage Followed by Interploidy Admixture. Mol. Biol. Evol. 32:1382–1395.

Arnold B.J., Lahner B., DaCosta J.M., Weisman C.M., Hollister J.D., Salt D.E., Bomblies K., Yant L. 2016. Borrowed alleles and convergence in serpentine adaptation. Proc. Natl. Acad. Sci. U.S.A. 113:8320–8325.

Ashburner K., McAllister H.A. 2013. The genus Betula: a taxonomic revision of birches:26-28 (London: Kew Publishing).

Atkinson M.D. 1992. *Betula Pendula* Roth (*B*. Verrucosa Ehrh.) and B. Pubescens Ehrh. J. Ecol. 80:837–870.

Aury J.-M., Jaillon O., Duret L., Noel B., Jubin C., Porcel B.M., Ségurens B., Daubin V., Anthouard V., Aiach N., Arnaiz O., Billaut A., Beisson J., Blanc I., Bouhouche K., Câmara F., Duharcourt S., Guigo R., Gogendeau D., Katinka M., Keller A.-M., Kissmehl R., Klotz C., Koll F., Le Mouël A., Lepère G., Malinsky S., Nowacki M., Nowak J.K., Plattner H., Poulain J., Ruiz F., Serrano V., Zagulski M., Dessen P., Bétermier M., Weissenbach J., Scarpelli C., Schächter V., Sperling L., Meyer E., Cohen J., Wincker P. 2006. Global trends of whole-genome duplications revealed by the ciliate *Paramecium tetraurelia*. Nature. 444:171–178.

Baack E., Melo M.C., Rieseberg L.H., Ortiz-Barrientos D. 2015. The origins of reproductive isolation in plants. New Phytol. 207:968–984.

Baduel P., Bray S., Vallejo-Marin M., Kolář F., Yant L. 2018. The “Polyploid Hop”: Shifting Challenges and Opportunities Over the Evolutionary Lifespan of Genome Duplications. Front. Ecol. Evol. 6.

Bagherieh-Najjar M.B., de Vries O.M.H., Hille J., Dijkwel P.P. 2005. Arabidopsis RecQl4A suppresses homologous recombination and modulates DNA damage responses. Plant J. 43:789– 798.

Bao S., Tibbetts R.S., Brumbaugh K.M., Fang Y., Richardson D.A., Ali A., Chen S.M., Abraham R.T., Wang X.-F. 2001. ATR/ATM-mediated phosphorylation of human Rad17 is required for genotoxic stress responses. Nature. 411:969–974.

Barker M.S., Arrigo N., Baniaga A.E., Li Z., Levin D.A. 2016. On the relative abundance of autopolyploids and allopolyploids. New Phytol. 210:391–398.

Barton N.H., Hewitt G.M. 1989. Adaptation, speciation and hybrid zones. Nature. 341:497–503.

Beaumont M.A., Zhang W., Balding D.J. 2002. Approximate Bayesian Computation in Population Genetics. Genetics. 162:2025–2035.

Beaumont M.A. 2019. Approximate Bayesian Computation. Annu. Rev. Stat. Appl. 6:379–403.

Berthelot C., Brunet F., Chalopin D., Juanchich A., Bernard M., Noël B., Bento P., Da Silva C., Labadie K., Alberti A., Aury J.-M., Louis A., Dehais P., Bardou P., Montfort J., Klopp C., Cabau C., Gaspin C., Thorgaard G.H., Boussaha M., Quillet E., Guyomard R., Galiana D., Bobe J., Volff J.-N., Genêt C., Wincker P., Jaillon O., Crollius H.R., Guiguen Y. 2014. The rainbow trout genome provides novel insights into evolution after whole-genome duplication in vertebrates. Nat. Commun. 5:3657.

Bertrand Y.J.K., Scheen A.-C., Marcussen T., Pfeil B.E., de Sousa F., Oxelman B. 2015. Assignment of homoeologs to parental genomes in allopolyploids for species tree inference, with an example from *Fumaria* (Papaveraceae). Syst. Biol. 64:448–471.

Bleeker W., Hurka H. 2001. Introgressive hybridization in *Rorippa* (Brassicaceae): gene flow and its consequences in natural and anthropogenic habitats. Mol. Ecol. 10:2013–2022.

Bleeker W. 2003. Hybridization and *Rorippa austriaca* (Brassicaceae) invasion in Germany. Mol. Ecol. 12:1831–1841.

Bomblies K., Higgins J.D., Yant L. 2015. Meiosis evolves: adaptation to external and internal environments. New Phytol. 208:306–323.

Bomblies K. 2023. Learning to tango with four (or more): the molecular basis of adaptation to polyploid meiosis. Plant Reprod. 36:107–124.

Bowers J.E., Chapman B.A., Rong J., Paterson A.H. 2003. Unravelling angiosperm genome evolution by phylogenetic analysis of chromosomal duplication events. Nature. 422:433–438.

Braasch I., Postlethwait J.H. 2012. Polyploidy in Fish and the Teleost Genome Duplication. In: Soltis P.S., Soltis D.E., editors. Polyploidy and Genome Evolution. Berlin, Heidelberg: Springer. p. 341–383.

Brown I.R., Al-Dawoody D. 1979. Observations on meiosis in three cytotypes of *Betula alba* L. New Phytol. 83:801–811.

Bryant D., Moulton V. 2004. Neighbor-Net: An agglomerative method for the construction of phylogenetic networks. Mol. Biol. Evol. 21:255–265.

Burns R., Mandáková T., Gunis J., Soto-Jiménez L.M., Liu C., Lysak M.A., Novikova P.Y., Nordborg M. 2021. Gradual evolution of allopolyploidy in *Arabidopsis suecica*. Nat Ecol Evol. 5:1367–1381.

Caldwell K.S., Michelmore R.W. 2009. *Arabidopsis thaliana* Genes Encoding Defense Signaling and Recognition Proteins Exhibit Contrasting Evolutionary Dynamics. Genetics. 181:671–684.

Cao H., Bowling S.A., Gordon A.S., Dong X. 1994. Characterization of an Arabidopsis Mutant That Is Nonresponsive to Inducers of Systemic Acquired Resistance. Plant Cell. 6:1583–1592.

Cerca J., Armstrong E.E., Vizueta J., Fernández R., Dimitrov D., Petersen B., Prost S., Rozas J., Petrov D., Gillespie R.G. 2021. The *Tetragnatha kauaiensis* Genome Sheds Light on the Origins of Genomic Novelty in Spiders. Genome Biol. Evol. 13:evab262.

Chalhoub B., Denoeud F., Liu S., Parkin I.A.P., Tang H., Wang X., Chiquet J., Belcram H., Tong C., Samans B., Corréa M., Da Silva C., Just J., Falentin C., Koh C.S., Le Clainche I., Bernard M., Bento P., Noel B., Labadie K., Alberti A., Charles M., Arnaud D., Guo H., Daviaud C., Alamery S., Jabbari K., Zhao M., Edger P.P., Chelaifa H., Tack D., Lassalle G., Mestiri I., Schnel N., Le Paslier M.-C., Fan G., Renault V., Bayer P.E., Golicz A.A., Manoli S., Lee T.-H., Thi V.H.D., Chalabi S., Hu Q., Fan C., Tollenaere R., Lu Y., Battail C., Shen J., Sidebottom C.H.D., Wang X., Canaguier A., Chauveau A., Bérard A., Deniot G., Guan M., Liu Z., Sun F., Lim Y.P., Lyons E., Town C.D., Bancroft I., Wang X., Meng J., Ma J., Pires J.C., King G.J., Brunel D., Delourme R., Renard M., Aury J.-M., Adams K.L., Batley J., Snowdon R.J., Tost J., Edwards D., Zhou Y., Hua W., Sharpe A.G., Paterson A.H., Guan C., Wincker P. 2014. Early allopolyploid evolution in the post-Neolithic *Brassica napus* oilseed genome. Science. 345:950–953.

Chen X., Lu L., Mayer K.S., Scalf M., Qian S., Lomax A., Smith L.M., Zhong X. 2016. POWERDRESS interacts with HISTONE DEACETYLASE 9 to promote aging in Arabidopsis. eLife. 5:e17214.

Chen S., Wang Y., Yu L., Zheng T., Wang S., Yue Z., Jiang J., Kumari S., Zheng C., Tang H., Li J., Li Y., Chen J., Zhang W., Kuang H., Robertson J.S., Zhao P.X., Li H., Shu S., Yordanov Y.S., Huang H., Goodstein D.M., Gai Y., Qi Q., Min J., Xu C., Wang S., Qu G.-Z., Paterson A.H., Sankoff D., Wei H., Liu G., Yang C. 2021. Genome sequence and evolution of *Betula platyphylla*. Hort. Res. 8:37.

Cheng H., Liu J., Wen J., Nie X., Xu L., Chen N., Li Z., Wang Q., Zheng Z., Li M., Cui L., Liu Z., Bian J., Wang Z., Xu S., Yang Q., Appels R., Han D., Song W., Sun Q., Jiang Y. 2019. Frequent intra- and inter-species introgression shapes the landscape of genetic variation in bread wheat. Genome Biol. 20:136.

Chenuil A., Galtier N., Berrebi P. 1999. A test of the hypothesis of an autopolyploid vs. allopolyploid origin for a tetraploid lineage: application to the genus Barbus (Cyprinidae). Heredity. 82:373–380.

Chester M., Gallagher J.P., Symonds V.V., Cruz da Silva A.V., Mavrodiev E.V., Leitch A.R., Soltis P.S., Soltis D.E. 2012. Extensive chromosomal variation in a recently formed natural allopolyploid species, *Tragopogon miscellus* (Asteraceae). Proc. Natl. Acad. Sci. U.S.A. 109:1176–1181.

Cifuentes M., Grandont L., Moore G., Chèvre A.M., Jenczewski E. 2010. Genetic regulation of meiosis in polyploid species: new insights into an old question. New Phytol. 186:29–36.

Cingolani P., Platts A., Wang L.L., Coon M., Nguyen T., Wang L., Land S.J., Lu X., Ruden D.M. 2012. A program for annotating and predicting the effects of single nucleotide polymorphisms, SnpEff. Fly. 6:80–92.

Clark L.V., Stewart J.R., Nishiwaki A., Toma Y., Kjeldsen J.B., Jørgensen U., Zhao H., Peng J., Yoo J.H., Heo K., Yu C.Y., Yamada T., Sacks E.J. 2015. Genetic structure of *Miscanthus sinensis* and *Miscanthus sacchariflorus* in Japan indicates a gradient of bidirectional but asymmetric introgression. J. Exp. Bot. 66:4213–4225.

Comai L. 2005. The advantages and disadvantages of being polyploid. Nat. Rev. Genet. 6:836– 846.

Cornish-Bowden A. 1985. Nomenclature for incompletely specified bases in nucleic acid sequences: recommendations 1984. Nucleic Acids Res. 13:3021–3030.

Danecek P., Bonfield J.K., Liddle J., Marshall J., Ohan V., Pollard M.O., Whitwham A., Keane T., McCarthy S.A., Davies R.M., Li H. 2021. Twelve years of SAMtools and BCFtools. GigaScience. 10.

Dasmahapatra K.K., Walters J.R., Briscoe A.D., Davey J.W., Whibley A., Nadeau N.J., Zimin A.V., Hughes D.S.T., Ferguson L.C., Martin S.H., Salazar C., Lewis J.J., Adler S., Ahn S.-J., Baker D.A., Baxter S.W., Chamberlain N.L., Chauhan R., Counterman B.A., Dalmay T., Gilbert L.E., Gordon K., Heckel D.G., Hines H.M., Hoff K.J., Holland P.W.H., Jacquin-Joly E., Jiggins F.M., Jones R.T., Kapan D.D., Kersey P., Lamas G., Lawson D., Mapleson D., Maroja L.S., Martin A., Moxon S., Palmér W.J., Papa R., Papanicolaou A., Pauchet Y., Ray D.A., Rosser N., Salzberg S.L., Supple M.A., Surridge A., Tenger-Trolander A., Vogel H., Wilkinson P.A., Wilson D., Yorke J.A., Yuan F., Balmuth A.L., Eland C., Gharbi K., Thomson M., Gibbs R.A., Han Y., Jayaseelan J.C., Kovar C., Mathew T., Muzny D.M., Ongeri F., Pu L.-L., Qu J., Thornton R.L., Worley K.C., Wu Y.-Q., Linares M., Blaxter M.L., ffrench-Constant R.H., Joron M., Kronforst M.R., Mullen S.P., Reed R.D., Scherer S.E., Richards S., Mallet J., Owen McMillan W., Jiggins C.D., The Heliconius Genome Consortium. 2012. Butterfly genome reveals promiscuous exchange of mimicry adaptations among species. Nature. 487:94–98.

De Bodt S., Maere S., Van de Peer Y. 2005. Genome duplication and the origin of angiosperms. Trends Ecol. Evol. 20:591–597.

de Groot W.J., Thomas P.A., Wein R.W. 1997. *Betula nana* L. and *Betula glandulosa* Michx. J. Ecol. 85:241–264.

Dehal P., Boore J.L. 2005. Two Rounds of Whole Genome Duplication in the Ancestral Vertebrate. PLOS Biol. 3:e314.

Ding J., Zhang B., Li Y., André D., Nilsson O. 2021. Phytochrome B and PHYTOCHROME INTERACTING FACTOR8 modulate seasonal growth in trees. New Phytol. 232:2339–2352.

Dong M.A., Farré E.M., Thomashow M.F. 2011. CIRCADIAN CLOCK-ASSOCIATED 1 and LATE ELONGATED HYPOCOTYL regulate expression of the C-REPEAT BINDING FACTOR (CBF) pathway in Arabidopsis. Proc. Natl. Acad. Sci. U.S.A. 108:7241–7246.

Dowling T.E., Secor C.L. 1997. The Role of Hybridization and Introgression in the Diversification of Animals. Annu. Rev. Ecol. Syst. 28:593–619.

Eidesen P.B., Alsos I.G., Brochmann C. 2015. Comparative analyses of plastid and AFLP data suggest different colonization history and asymmetric hybridization between *Betula pubescens* and *B. nana*. Mol. Ecol. 24:3993–4009.

Ellstrand N.C. 2014. Is gene flow the most important evolutionary force in plants? Am. J. Bot. 101:737–753.

Enns L.C., Kanaoka M.M., Torii K.U., Comai L., Okada K., Cleland R.E. 2005. Two callose synthases, GSL1 and GSL5, play an essential and redundant role in plant and pollen development and in fertility. Plant Mol. Biol. 58:333–349.

Excoffier L., Foll M., Petit R.J. 2009. Genetic Consequences of Range Expansions. Annu. Rev. Ecol. Evol. Syst. 40:481–501.

Excoffier L., Dupanloup I., Huerta-Sánchez E., Sousa V.C., Foll M. 2013. Robust Demographic Inference from Genomic and SNP Data. PLOS Genet. 9:e1003905.

Felber F. 1991. Establishment of a tetraploid cytotype in a diploid population: Effect of relative fitness of the cytotypes. J. Evol. Biol. 4:195–207.

Filiault D.L., Wessinger C.A., Dinneny J.R., Lutes J., Borevitz J.O., Weigel D., Chory J., Maloof J.N. 2008. Amino acid polymorphisms in *Arabidopsis* phytochrome B cause differential responses to light. Proc. Natl. Acad. Sci. U.S.A. 105:3157–3162.

Fletcher W., Yang Z. 2009. INDELible: a flexible simulator of biological sequence evolution. Mol. Biol. Evol. 26:1879–1888.

Fontaine M.C., Pease J.B., Steele A., Waterhouse R.M., Neafsey D.E., Sharakhov I.V., Jiang X., Hall A.B., Catteruccia F., Kakani E., Mitchell S.N., Wu Y.-C., Smith H.A., Love R.R., Lawniczak M.K., Slotman M.A., Emrich S.J., Hahn M.W., Besansky N.J. 2015. Extensive introgression in a malaria vector species complex revealed by phylogenomics. Science. 347:1258524.

Freyman W.A., Johnson M.G., Rothfels C.J. 2023. homologizer: Phylogenetic phasing of gene copies into polyploid subgenomes. Methods Ecol. Evol. 14:1230–1244.

Fujiwara S., Oda A., Yoshida R., Niinuma K., Miyata K., Tomozoe Y., Tajima T., Nakagawa M., Hayashi K., Coupland G., Mizoguchi T. 2008. Circadian Clock Proteins LHY and CCA1 Regulate SVP Protein Accumulation to Control Flowering in *Arabidopsis*. Plant Cell. 20:2960– 2971.

Gaeta R.T., Chris Pires J. 2010. Homoeologous recombination in allopolyploids: the polyploid ratchet. New Phytol. 186:18–28.

Grant P.R., and Grant B.R. 1992. Hybridization of Bird Species. Science. 256:193–197.

Grant P.R., and Grant B.R. 2020. Triad hybridization via a conduit species. Proc. Natl. Acad. Sci. U.S.A. 117:7888–7896.

Hajdu A., Ádám É., Sheerin D.J., Dobos O., Bernula P., Hiltbrunner A., Kozma-Bognár L., Nagy F. 2015. High-level expression and phosphorylation of phytochrome B modulates flowering time in *Arabidopsis*. Plant J. 83:794–805.

Hardigan M.A., Laimbeer F.P.E., Newton L., Crisovan E., Hamilton J.P., Vaillancourt B., Wiegert-Rininger K., Wood J.C., Douches D.S., Farré E.M., Veilleux R.E., Buell C.R. 2017. Genome diversity of tuber-bearing Solanum uncovers complex evolutionary history and targets of domestication in the cultivated potato. Proc. Natl. Acad. Sci. USA. 114:E9999–E10008.

Hartung F., Suer S., Knoll A., Wurz-Wildersinn R., Puchta H. 2008. Topoisomerase 3α and RMI1 Suppress Somatic Crossovers and Are Essential for Resolution of Meiotic Recombination Intermediates in Arabidopsis thaliana. PLoS Genet. 4:e1000285.

Heitzeberg F., Chen I.-P., Hartung F., Orel N., Angelis K.J., Puchta H. 2004. The Rad17 homologue of Arabidopsis is involved in the regulation of DNA damage repair and homologous recombination. Plant J. 38:954–968.

Higgins J.D., Ferdous M., Osman K., Franklin F.C.H. 2011. The RecQ helicase AtRECQ4A is required to remove inter-chromosomal telomeric connections that arise during meiotic recombination in Arabidopsis. Plant J. 65:492–502.

Hohmann N., Koch M.A. 2017. An Arabidopsis introgression zone studied at high spatio-temporal resolution: interglacial and multiple genetic contact exemplified using whole nuclear and plastid genomes. BMC Genom. 18:1–18.

Howland D.E., Oliver R.P., Davy A.J. 1995. Morphological and molecular variation in natural populations of Betula. New Phytol. 130:117–124.

Hu Q., Zhang C., Xue Z., Ma L., Liu W., Shen Y., Ma B., Cheng Z. 2018. OsRAD17 Is Required for Meiotic Double-Strand Break Repair and Plays a Redundant Role With OsZIP4 in Synaptonemal Complex Assembly. Front. Plant Sci. 9.

Husband B.C. 2004. The role of triploid hybrids in the evolutionary dynamics of mixed-ploidy populations. Biol. J. Linn. Soc. 82:537–546.

Huson D.H., Bryant D. 2006. Application of phylogenetic networks in evolutionary studies. Mol. Biol. Evol. 23:254–267.

Ingvarsson P.K., Garcia M.V., Luquez V., Hall D., Jansson S. 2008. Nucleotide Polymorphism and Phenotypic Associations Within and Around the phytochrome B2 Locus in European Aspen (*Populus tremula*, Salicaceae). Genetics. 178:2217–2226.

Jadwiszczak K.A., Banaszek A., Jabłońska E., Sozinov O.V. 2012. Chloroplast DNA variation of *Betula humilis* Schrk. in Poland and Belarus. Tree Genet. Genomes. 8:1017–1030.

Järvinen P., Palmé A., Morales L.O., Lännenpää M., Keinänen M., Sopanen T., Lascoux M. 2004. Phylogenetic relationships of Betula species (Betulaceae) based on nuclear ADH and chloroplast matK sequences. Am. J. Bot.. 91:1834–1845.

Jeffers, R. M. 1971. Research at the Institute of Forest Genetics, Rhinelander, Wisconsin. USDA Forest Service, Research Paper NC-67, p. 17.

Jiang X., Song Q., Ye W., Chen Z.J. 2021. Concerted genomic and epigenomic changes accompany stabilization of *Arabidopsis* allopolyploids. Nat Ecol Evol. 5:1382–1393.

Johnsson H. 1944. Triploidy in *Betula alba* L. Bot. Kotiser. 97: 85.

Jombart T., Devillard S., Balloux F. 2010. Discriminant analysis of principal components: a new method for the analysis of genetically structured populations. BMC Genet. 11:1–15.

Jones G., Sagitov S., Oxelman B. 2013. Statistical inference of allopolyploid species networks in the presence of incomplete lineage sorting. Syst. Biol. 62:467–478.

Jørgensen M.H., Ehrich D., Schmickl R., Koch M.A., Brysting A.K. 2011. Interspecific and interploidal gene flow in Central European Arabidopsis (Brassicaceae). BMC Evol. Biol. 11:346.

Joubès J., Raffaele S., Bourdenx B., Garcia C., Laroche-Traineau J., Moreau P., Domergue F., Lessire R. 2008. The VLCFA elongase gene family in Arabidopsis thaliana: phylogenetic analysis, 3D modelling and expression profiling. Plant Mol. Biol. 67:547–566.

Jung J.-H., Domijan M., Klose C., Biswas S., Ezer D., Gao M., Khattak A.K., Box M.S., Charoensawan V., Cortijo S., Kumar M., Grant A., Locke J.C.W., Schäfer E., Jaeger K.E., Wigge P.A. 2016. Phytochromes function as thermosensors in *Arabidopsis*. Science. 354:886–889.

Kaczorowski K.A., Quail P.H. 2003. Arabidopsis PSEUDO-RESPONSE REGULATOR7 Is a Signaling Intermediate in Phytochrome-Regulated Seedling Deetiolation and Phasing of the Circadian Clock. Plant Cell. 15:2654–2665.

Keightley P.D., Jackson B.C. 2018. Inferring the Probability of the Derived vs. the Ancestral Allelic State at a Polymorphic Site. Genetics. 209:897–906.

Keller I., Wagner C.E., Greuter L., Mwaiko S., Selz O.M., Sivasundar A., Wittwer S., Seehausen O. 2013. Population genomic signatures of divergent adaptation, gene flow and hybrid speciation in the rapid radiation of Lake Victoria cichlid fishes. Mol. Ecol. 22:2848–2863.

Kellis M., Birren B.W., Lander E.S. 2004. Proof and evolutionary analysis of ancient genome duplication in the yeast *Saccharomyces cerevisiae*. Nature. 428:617–624.

Kim H.-J., Kim Y.-K., Park J.-Y., Kim J. 2002. Light signalling mediated by phytochrome plays an important role in cold-induced gene expression through the C-repeat/dehydration responsive element (C/DRE) in *Arabidopsis thaliana*. Plant J. 29:693–704.

Kim M., Cui M.-L., Cubas P., Gillies A., Lee K., Chapman M.A., Abbott R.J., Coen E. 2008. Regulatory Genes Control a Key Morphological and Ecological Trait Transferred between Species. Science. 322:1116–1119.

Kim Y., Park S., Gilmour S.J., Thomashow M.F. 2013. Roles of CAMTA transcription factors and salicylic acid in configuring the low-temperature transcriptome and freezing tolerance of Arabidopsis. Plant J. 75:364–376.

Kim Y.J., Wang R., Gao L., Li D., Xu C., Mang H., Jeon J., Chen X., Zhong X., Kwak J.M., Mo B., Xiao L., Chen X. 2016. POWERDRESS and HDA9 interact and promote histone H3 deacetylation at specific genomic sites in Arabidopsis. Proc. Natl. Acad. Sci. U.S.A. 113:14858– 14863.

Koropachinskii I.Yu. 2013. Natural hybridization and taxonomy of birches in North Asia. Contemp. Probl. Ecol. 6:350–369.

Krehenwinkel H., Tautz D. 2013. Northern range expansion of European populations of the wasp spider *Argiope bruennichi* is associated with global warming–correlated genetic admixture and population-specific temperature adaptations. Mol. Ecol. 22:2232–2248.

Lafon-Placette C., Johannessen I.M., Hornslien K.S., Ali M.F., Bjerkan K.N., Bramsiepe J., Glöckle B.M., Rebernig C.A., Brysting A.K., Grini P.E., Köhler C. 2017. Endosperm-based hybridization barriers explain the pattern of gene flow between *Arabidopsis lyrat*a and *Arabidopsis arenosa* in Central Europe. Proc. Natl. Acad. Sci. USA. 114:E1027–E1035.

Lamichhaney S., Berglund J., Almén M.S., Maqbool K., Grabherr M., Martinez-Barrio A., Promerová M., Rubin C.-J., Wang C., Zamani N., Grant B.R., Grant P.R., Webster M.T., Andersson L. 2015. Evolution of Darwin’s finches and their beaks revealed by genome sequencing. Nature. 518:371–375.

Lashermes P., Hueber Y., Combes M.-C., Severac D., Dereeper A. 2016. Inter-genomic DNA Exchanges and Homeologous Gene Silencing Shaped the Nascent Allopolyploid Coffee Genome (*Coffea arabica* L.). G3: Genes Genomes Genet. 6:2937–2948.

Lautenschlager U., Wagner F., Oberprieler C. 2020. AllCoPol: inferring allele co-ancestry in polyploids. BMC Bioinf. 21:441.

Leal J.L., Milesi P., Salojärvi J., Lascoux M. 2023. Phylogenetic Analysis of Allotetraploid Species Using Polarized Genomic Sequences. Syst. Biol. 72:372–390.

Legris M., Klose C., Burgie E.S., Rojas C.C.R., Neme M., Hiltbrunner A., Wigge P.A., Schäfer E., Vierstra R.D., Casal J.J. 2016. Phytochrome B integrates light and temperature signals in *Arabidopsis*. Science. 354:897–900.

Li J., Shoup S., Chen Z. 2005. Phylogenetics of *Betula* (*Betulaceae*) inferred from sequences of nuclear ribosomal DNA. Rhodora. 107:69–86.

Li J., Shoup S., Chen Z. 2007. Phylogenetic Relationships of Diploid Species of *Betula* (*Betulaceae*) Inferred from DNA Sequences of Nuclear Nitrate Reductase. Syst. Bot. 32:357– 365.

Li H., Durbin R. 2009. Fast and accurate short read alignment with Burrows–Wheeler transform. Bioinformatics. 25:1754–1760.

Li F., Fan G., Lu C., Xiao G., Zou C., Kohel R.J., Ma Z., Shang H., Ma X., Wu J., Liang X., Huang G., Percy R.G., Liu K., Yang W., Chen W., Du X., Shi C., Yuan Y., Ye W., Liu X., Zhang X., Liu W., Wei H., Wei S., Huang G., Zhang X., Zhu S., Zhang H., Sun F., Wang X., Liang J., Wang J., He Q., Huang L., Wang J., Cui J., Song G., Wang K., Xu X., Yu J.Z., Zhu Y., Yu S. 2015. Genome sequence of cultivated Upland cotton (*Gossypium hirsutum* TM-1) provides insights into genome evolution. Nat. Biotechnol. 33:524–530.

Li Z., Tiley G.P., Galuska S.R., Reardon C.R., Kidder T.I., Rundell R.J., Barker M.S. 2018a. Multiple large-scale gene and genome duplications during the evolution of hexapods. Proc. Natl. Acad. Sci. U.S.A. 115:4713–4718.

Li X., Yu H., Jiao Y., Shahid M.Q., Wu J., Liu X. 2018b. Genome-wide analysis of DNA polymorphisms, the methylome and transcriptome revealed that multiple factors are associated with low pollen fertility in autotetraploid rice. PLoS One. 13:e0201854.

Li L., Milesi P., Tiret M., Chen J., Sendrowski J., Baison J., Chen Z., Zhou L., Karlsson B., Berlin M., Westin J., Garcia-Gil M.R., Wu H.X., Lascoux M. 2022. Teasing apart the joint effect of demography and natural selection in the birth of a contact zone. New Phytol. 236:1976–1987.

Lihová J., Kučera J., Perný M., Marhold K. 2007. Hybridization between Two Polyploid Cardamine (Brassicaceae) Species in North-western Spain: Discordance Between Morphological and Genetic Variation Patterns. Ann. Bot. 99:1083–1096.

Lim C.J., Park J., Shen M., Park H.J., Cheong M.S., Park K.S., Baek D., Bae M.J., Ali A., Jan M., Lee S.Y., Lee B., Kim W.-Y., Pardo J.M., Yun D.-J. 2020. The Histone-Modifying Complex PWR/HOS15/HD2C Epigenetically Regulates Cold Tolerance. Plant Physiol. 184:1097–1111.

Liu Z., Hall J.D., Mount D.W. 2001. Arabidopsis UVH3 gene is a homolog of the *Saccharomyces cerevisiae* RAD2 and human XPG DNA repair genes. Plant J. 26:329–338.

Lloyd A., Bomblies K. 2016. Meiosis in autopolyploid and allopolyploid Arabidopsis. Curr. Opin. Plant Biol. 30:116–122.

Lloyd A., Blary A., Charif D., Charpentier C., Tran J., Balzergue S., Delannoy E., Rigaill G., Jenczewski E. 2018. Homoeologous exchanges cause extensive dosage-dependent gene expression changes in an allopolyploid crop. New Phytol. 217:367–377.

Ma L.-J., Ibrahim A.S., Skory C., Grabherr M.G., Burger G., Butler M., Elias M., Idnurm A., Lang B.F., Sone T., Abe A., Calvo S.E., Corrochano L.M., Engels R., Fu J., Hansberg W., Kim J.-M., Kodira C.D., Koehrsen M.J., Liu B., Miranda-Saavedra D., O’Leary S., Ortiz-Castellanos L., Poulter R., Rodriguez-Romero J., Ruiz-Herrera J., Shen Y.-Q., Zeng Q., Galagan J., Birren B.W., Cuomo C.A., Wickes B.L. 2009. Genomic Analysis of the Basal Lineage Fungus *Rhizopus oryzae* Reveals a Whole-Genome Duplication. PLOS Genet. 5:e1000549.

Macqueen D.J., Johnston I.A. 2014. A well-constrained estimate for the timing of the salmonid whole genome duplication reveals major decoupling from species diversification. Proc. R. Soc. Lond. B. 281:20132881.

Mallet J. 2005. Hybridization as an invasion of the genome. Trends Ecol. Evol. 20:229–237.

Mallet J. 2007. Hybrid speciation. Nature. 446:279–283.

Mallet J., Besansky N., Hahn M.W. 2016. How reticulated are species? BioEssays. 38:140–149.

Mallo D., De Oliveira Martins L., Posada D. 2016. SimPhy: phylogenomic simulation of gene, locus, and species trees. Syst. Biol. 65:334–344.

Mandel J.R., Barker M.S., Bayer R.J., Dikow R.B., Gao T.-G., Jones K.E., Keeley S., Kilian N., Ma H., Siniscalchi C.M., Susanna A., Thapa R., Watson L., Funk V.A. 2017. The Compositae Tree of Life in the age of phylogenomics. J. Syst. Evol. 55:405–410.

Marburger S., Monnahan P., Seear P.J., Martin S.H., Koch J., Paajanen P., Bohutínská M., Higgins J.D., Schmickl R., Yant L. 2019. Interspecific introgression mediates adaptation to whole genome duplication. Nat. Commun. 10:5218.

Marhold K., Lihová J. 2006. Polyploidy, hybridization and reticulate evolution: lessons from the Brassicaceae. Plant Syst. Evol. 259:143–174.

Maruyama K., Takeda M., Kidokoro S., Yamada K., Sakuma Y., Urano K., Fujita M., Yoshiwara K., Matsukura S., Morishita Y., Sasaki R., Suzuki H., Saito K., Shibata D., Shinozaki K., Yamaguchi-Shinozaki K. 2009. Metabolic Pathways Involved in Cold Acclimation Identified by Integrated Analysis of Metabolites and Transcripts Regulated by DREB1A and DREB2A. Plant Physiol. 150:1972–1980.

McDonald D.B., Parchman T.L., Bower M.R., Hubert W.A., Rahel F.J. 2008. An introduced and a native vertebrate hybridize to form a genetic bridge to a second native species. Proc. Natl. Acad. Sci. U.S.A. 105:10837–10842.

McKenna A., Hanna M., Banks E., Sivachenko A., Cibulskis K., Kernytsky A., Garimella K., Altshuler D., Gabriel S., Daly M., DePristo M.A. 2010. The Genome Analysis Toolkit: A MapReduce framework for analyzing next-generation DNA sequencing data. Genome Res. 20:1297–1303.

Meier J.I., Marques D.A., Mwaiko S., Wagner C.E., Excoffier L., Seehausen O. 2017. Ancient hybridization fuels rapid cichlid fish adaptive radiations. Nat. Commun. 8:14363.

Merret R., Descombin J., Juan Y., Favory J.-J., Carpentier M.-C., Chaparro C., Charng Y., Deragon J.-M., Bousquet-Antonelli C. 2013. XRN4 and LARP1 Are Required for a Heat-Triggered mRNA Decay Pathway Involved in Plant Acclimation and Survival during Thermal Stress. Cell Rep. 5:1279–1293.

Milesi P., Kastally C., Dauphin B., Cervantes S., Bagnoli F., Budde K.B., Cavers S., Ojeda D.I., Fady B., Faivre-Rampant P., González-Martínez S.C., Grivet D., Gugerli F., Jorge V., Lesur-Kupin I., Olsson S., Opgenoorth L., Pinosio S., Plomion C., Rellstab C., Rogier O., Scalabrin S., Scotti I., Vendramin G.G., Westergren M., Consortium G., Lascoux M., Pyhäjärvi T. 2023. Synchronous effective population size changes and genetic stability of forest trees through glacial cycles. bioRxiv:2023.01.05.522822.

Minh B.Q., Schmidt H.A., Chernomor O., Schrempf D., Woodhams M.D., von Haeseler A., Lanfear R. 2020. IQ-TREE 2: new models and efficient methods for phylogenetic inference in the genomic era. Mol. Biol. Evol. 37:1530–1534.

Monnahan P., Kolář F., Baduel P., Sailer C., Koch J., Horvath R., Laenen B., Schmickl R., Paajanen P., Šrámková G., Bohutínská M., Arnold B., Weisman C.M., Marhold K., Slotte T., Bomblies K., Yant L. 2019. Pervasive population genomic consequences of genome duplication in *Arabidopsis arenosa*. *Nat*. Ecol. Evol. 3:457–468.

Morgan C., White M.A., Franklin F.C.H., Zickler D., Kleckner N., Bomblies K. 2021. Evolution of crossover interference enables stable autopolyploidy by ensuring pairwise partner connections in *Arabidopsis arenosa*. Curr. Biol. 31:4713–4726.e4.

Muñoz-Rodríguez P., Carruthers T., Wood J.R.I., Williams B.R.M., Weitemier K., Kronmiller B., Ellis D., Anglin N.L., Longway L., Harris S.A., Rausher M.D., Kelly S., Liston A., Scotland R.W. 2018. Reconciling Conflicting Phylogenies in the Origin of Sweet Potato and Dispersal to Polynesia. Curr. Biol. 28:1246–1256.e12.

Nibau C., Gonzalo A., Evans A., Sweet-Jones W., Phillips D., Lloyd A. 2022. Meiosis in allopolyploid *Arabidopsis suecica*. Plant J. 111:1110–1122.

Nieto Feliner G., Casacuberta J., Wendel J.F. 2020. Genomics of Evolutionary Novelty in Hybrids and Polyploids. Front. Genet. 11.

Nossa C.W., Havlak P., Yue J.-X., Lv J., Vincent K.Y., Brockmann H.J., Putnam N.H. 2014. Joint assembly and genetic mapping of the Atlantic horseshoe crab genome reveals ancient whole genome duplication. GigaScience. 3:2047–217X-3–9.

Novikova P.Yu., Tsuchimatsu T., Simon S., Nizhynska V., Voronin V., Burns R., Fedorenko O.M., Holm S., Säll T., Prat E., Marande W., Castric V., Nordborg M. 2017. Genome Sequencing Reveals the Origin of the Allotetraploid *Arabidopsis suecica*. Mol. Biol. Evol. 34:957–968.

Novikova P.Y., Brennan I.G., Booker W., Mahony M., Doughty P., Lemmon A.R., Lemmon E.M., Roberts J.D., Yant L., de Peer Y.V., Keogh J.S., Donnellan S.C. 2020. Polyploidy breaks speciation barriers in Australian burrowing frogs *Neobatrachus*. PLOS Genet. 16:e1008769.

Oberprieler C., Wagner F., Tomasello S., Konowalik K. 2017. A permutation approach for inferring species networks from gene trees in polyploid complexes by minimising deep coalescences. Methods Ecol. Evol. 8:835–849.

Opgenoorth L., Dauphin B., Benavides R., Heer K., Alizoti P., Martínez-Sancho E., Alía R., Ambrosio O., Audrey A., Auñón F., Avanzi C., Avramidou E., Bagnoli F., Barbas E., Bastias C.C., Bastien C., Ballesteros E., Beffa G., Bernier F., Bignalet H., Bodineau G., Bouic D., Brodbeck S., Brunetto W., Buchovska J., Buy M., Cabanillas-Saldaña A.M., Carvalho B., Cheval N., Climent J.M., Correard M., Cremer E., Danusevičius D., Del Caño F., Denou J.-L., di Gerardi N., Dokhelar B., Ducousso A., Eskild Nilsen A., Farsakoglou A.-M., Fonti P., Ganopoulos I., García del Barrio J.M., Gilg O., González-Martínez S.C., Graf R., Gray A., Grivet D., Gugerli F., Hartleitner C., Hollenbach E., Hurel A., Issehut B., Jean F., Jorge V., Jouineau A., Kappner J.-P., Kärkkäinen K., Kesälahti R., Knutzen F., Kujala S.T., Kumpula T.A., Labriola M., Lalanne C., Lambertz J., Lascoux M., Lejeune V., Le-Provost G., Levillain J., Liesebach M., López-Quiroga D., Meier B., Malliarou E., Marchon J., Mariotte N., Mas A., Matesanz S., Meischner H., Michotey C., Milesi P., Morganti S., Nievergelt D., Notivol E., Ostreng G., Pakull B., Perry A., Piotti A., Plomion C., Poinot N., Pringarbe M., Puzos L., Pyhäjärvi T., Raffin A., Ramírez-Valiente J.A., Rellstab C., Remi D., Richter S., Robledo-Arnuncio J.J., San Segundo S., Savolainen O., Schueler S., Schneck V., Scotti I., Semerikov V., Slámová L., Sønstebø J.H., Spanu I., Thevenet J., Tollefsrud M.M., Turion N., Vendramin G.G., Villar M., von Arx G., Westin J., Fady B., Myking T., Valladares F., Aravanopoulos F.A., Cavers S. 2021. The GenTree Platform: growth traits and tree-level environmental data in 12 European forest tree species. GigaScience. 10:giab010.

Oxelman B., Brysting A.K., Jones G.R., Marcussen T., Oberprieler C., Pfeil B.E. 2017. Phylogenetics of allopolyploids. Annu. Rev. Ecol. Evol. Syst. 48:543–557.

Palmé A.E., Su Q., Palsson S., Lascoux M. 2004. Extensive sharing of chloroplast haplotypes among European birches indicates hybridization among *Betula pendula*, B. pubescens and B. nana. Mol. Ecol. 13:167–178.

Pardo-Diaz C., Salazar C., Baxter S.W., Merot C., Figueiredo-Ready W., Joron M., McMillan W.O., Jiggins C.D. 2012. Adaptive Introgression across Species Boundaries in *Heliconius* Butterflies. PLoS Genet. 8:e1002752.

Pickrell J.K., Pritchard J.K. 2012. Inference of Population Splits and Mixtures from Genome-Wide Allele Frequency Data. PLoS Genet. 8:e1002967.

Pritchard J.K., Stephens M., Donnelly P. 2000. Inference of Population Structure Using Multilocus Genotype Data. Genetics. 155:945–959.

Ramsey J., Schemske D.W. 1998. Pathways, Mechanisms, and Rates of Polyploid Formation in Flowering Plants. Annu Rev Ecol Evol Syst. 29:467–501.

Ramsey J., Schemske D.W. 2002. Neopolyploidy in Flowering Plants. Annu. Rev. Ecol. Evol. Syst. 33:589–639.

Renard J., Niñoles R., Martínez-Almonacid I., Gayubas B., Mateos-Fernández R., Bissoli G., Bueso E., Serrano R., Gadea J. 2020. Identification of novel seed longevity genes related to oxidative stress and seed coat by genome-wide association studies and reverse genetics. Plant Cell Environ. 43:2523–2539.

Rieseberg L.H., Archer M.A., Wayne R.K. 1999. Transgressive segregation, adaptation and speciation. Heredity. 83:363–372.

Rieseberg L.H., Willis J.H. 2007. Plant speciation. Science. 317:910–914.

Rius M., Darling J.A. 2014. How important is intraspecific genetic admixture to the success of colonising populations? Trends Ecol. Evol. 29:233–242.

Robertson F.M., Gundappa M.K., Grammes F., Hvidsten T.R., Redmond A.K., Lien S., Martin S.A.M., Holland P.W.H., Sandve S.R., Macqueen D.J. 2017. Lineage-specific rediploidization is a mechanism to explain time-lags between genome duplication and evolutionary diversification. Genome Biol. 18:111.

Rothfels C.J. 2021. Polyploid phylogenetics. New Phytol. 230:66–72.

Roux C., Pannell J.R. 2015. Inferring the mode of origin of polyploid species from next-generation sequence data. Mol. Ecol. 24:1047–1059.

Salojärvi J., Smolander O.-P., Nieminen K., Rajaraman S., Safronov O., Safdari P., Lamminmäki A., Immanen J., Lan T., Tanskanen J., Rastas P., Amiryousefi A., Jayaprakash B., Kammonen J.I., Hagqvist R., Eswaran G., Ahonen V.H., Serra J.A., Asiegbu F.O., de Dios Barajas-Lopez J., Blande D., Blokhina O., Blomster T., Broholm S., Brosché M., Cui F., Dardick C., Ehonen S.E., Elomaa P., Escamez S., Fagerstedt K.V., Fujii H., Gauthier A., Gollan P.J., Halimaa P., Heino P.I., Himanen K., Hollender C., Kangasjärvi S., Kauppinen L., Kelleher C.T., Kontunen-Soppela S., Koskinen J.P., Kovalchuk A., Kärenlampi S.O., Kärkönen A.K., Lim K.-J., Leppälä J., Macpherson L., Mikola J., Mouhu K., Mähönen A.P., Niinemets Ü., Oksanen E., Overmyer K., Palva E.T., Pazouki L., Pennanen V., Puhakainen T., Poczai P., Possen B.J.H.M., Punkkinen M., Rahikainen M.M., Rousi M., Ruonala R., van der Schoot C., Shapiguzov A., Sierla M., Sipilä T.P., Sutela S., Teeri T.H., Tervahauta A.I., Vaattovaara A., Vahala J., Vetchinnikova L., Welling A., Wrzaczek M., Xu E., Paulin L.G., Schulman A.H., Lascoux M., Albert V.A., Auvinen P., Helariutta Y., Kangasjärvi J. 2017. Genome sequencing and population genomic analyses provide insights into the adaptive landscape of silver birch. Nat. Genet. 49:904–912.

Schenk M.F., Thienpont C.-N., Koopman W.J.M., Gilissen L.J.W.J., Smulders M.J.M. 2008. Phylogenetic relationships in *Betula* (*Betulaceae*) based on AFLP markers. Tree Genet. Genomes. 4:911–924.

Schmickl R., Marburger S., Bray S., Yant L. 2017. Hybrids and horizontal transfer: introgression allows adaptive allele discovery. J. Exp. Bot. 68:5453–5470.

Schmickl R., Yant L. 2021. Adaptive introgression: how polyploidy reshapes gene flow landscapes. New Phytol. 230:457–461.

Schwager E.E., Sharma P.P., Clarke T., Leite D.J., Wierschin T., Pechmann M., Akiyama-Oda Y., Esposito L., Bechsgaard J., Bilde T., Buffry A.D., Chao H., Dinh H., Doddapaneni H., Dugan S., Eibner C., Extavour C.G., Funch P., Garb J., Gonzalez L.B., Gonzalez V.L., Griffiths-Jones S., Han Y., Hayashi C., Hilbrant M., Hughes D.S.T., Janssen R., Lee S.L., Maeso I., Murali S.C., Muzny D.M., Nunes da Fonseca R., Paese C.L.B., Qu J., Ronshaugen M., Schomburg C., Schönauer A., Stollewerk A., Torres-Oliva M., Turetzek N., Vanthournout B., Werren J.H., Wolff C., Worley K.C., Bucher G., Gibbs R.A., Coddington J., Oda H., Stanke M., Ayoub N.A., Prpic N.-M., Flot J.-F., Posnien N., Richards S., McGregor A.P. 2017. The house spider genome reveals an ancient whole-genome duplication during arachnid evolution. BMC Biol. 15:62.

Seear P.J., France M.G., Gregory C.L., Heavens D., Schmickl R., Yant L., Higgins J.D. 2020. A novel allele of ASY3 is associated with greater meiotic stability in autotetraploid *Arabidopsis lyrata*. PLoS Genet. 16:e1008900.

Seehausen O. 2006. African cichlid fish: a model system in adaptive radiation research. Proc. R. Soc. B: Biol. Sci. 273:1987–1998.

Séguéla-Arnaud M., Crismani W., Larchevêque C., Mazel J., Froger N., Choinard S., Lemhemdi A., Macaisne N., Van Leene J., Gevaert K., De Jaeger G., Chelysheva L., Mercier R. 2015. Multiple mechanisms limit meiotic crossovers: TOP3α and two BLM homologs antagonize crossovers in parallel to FANCM. Proc. Natl. Acad. Sci. U.S.A. 112:4713–4718.

Seo M., Hanada A., Kuwahara A., Endo A., Okamoto M., Yamauchi Y., North H., Marion-Poll A., Sun T., Koshiba T., Kamiya Y., Yamaguchi S., Nambara E. 2006. Regulation of hormone metabolism in *Arabidopsis* seeds: phytochrome regulation of abscisic acid metabolism and abscisic acid regulation of gibberellin metabolism. Plant J. 48:354–366.

Serra H., Lambing C., Griffin C.H., Topp S.D., Nageswaran D.C., Underwood C.J., Ziolkowski P.A., Séguéla-Arnaud M., Fernandes J.B., Mercier R., Henderson I.R. 2018. Massive crossover elevation via combination of *HEI10* and *recq4a recq4b* during *Arabidopsis* meiosis. Proc. Natl. Acad. Sci. U.S.A. 115:2437–2442.

Shen H., Luong P., Huq E. 2007. The F-Box Protein MAX2 Functions as a Positive Regulator of Photomorphogenesis in Arabidopsis. Plant Physiol. 145:1471–1483.

Shi X., Sun X., Zhang Z., Feng D., Zhang Q., Han L., Wu J., Lu T. 2015. GLUCAN SYNTHASE-LIKE 5 (GSL5) Plays an Essential Role in Male Fertility by Regulating Callose Metabolism During Microsporogenesis in Rice. Plant Cell Physiol. 56:497–509.

Sicard A., Kappel C., Josephs E.B., Lee Y.W., Marona C., Stinchcombe J.R., Wright S.I., Lenhard M. 2015. Divergent sorting of a balanced ancestral polymorphism underlies the establishment of gene-flow barriers in Capsella. Nat. Commun. 6:7960.

Slotte T., Huang H., Lascoux M., Ceplitis A. 2008. Polyploid Speciation Did Not Confer Instant Reproductive Isolation in Capsella (Brassicaceae). Mol. Biol. Evol. 25:1472–1481.

Solís-Lemus C., Bastide P., Ané C. 2017. PhyloNetworks: a package for phylogenetic networks. Mol. Biol. Evol. 34:3292–3298.

Soltis P.S., Soltis D.E. 2009. The Role of Hybridization in Plant Speciation. Annu. Rev. Plant Biol. 60: 561–588

Somers D.E., Devlin P.F., Kay S.A. 1998. Phytochromes and Cryptochromes in the Entrainment of the *Arabidopsis* Circadian Clock. Science. 282:1488–1490.

Spoelhof J.P., Soltis P.S., Soltis D.E. 2017. Pure polyploidy: Closing the gaps in autopolyploid research. J. Syst. Evol. 55:340–352.

Stebbins G.L. 1956. Cytogenetics and Evolution of the Grass Family. Am. J. Bot. 43:890–905. Stebbins G. 1971. Chromosomal Evolution in Higher Plants. London: Edward Arnold Ltd.

Stern K. 1965. Tetrasome spaltung bei *Betula pubescens*. Silvae Genet. 14:56–57.

Suarez-Gonzalez A., Hefer C.A., Lexer C., Douglas C.J., Cronk Q.C.B. 2018. Introgression from *Populus balsamifera* underlies adaptively significant variation and range boundaries in *P. trichocarpa*. New Phytol. 217:416–427.

Sun F., Fan G., Hu Q., Zhou Y., Guan M., Tong C., Li J., Du D., Qi C., Jiang L., Liu W., Huang S., Chen W., Yu J., Mei D., Meng J., Zeng P., Shi J., Liu K., Wang X., Wang X., Long Y., Liang X., Hu Z., Huang G., Dong C., Zhang H., Li J., Zhang Y., Li L., Shi C., Wang J., Lee S.M.-Y., Guan C., Xu X., Liu S., Liu X., Chalhoub B., Hua W., Wang H. 2017. The high-quality genome of *Brassica napus* cultivar ‘ZS11’ reveals the introgression history in semi-winter morphotype. Plant J. 92:452–468.

Tavaré S., Balding D.J., Griffiths R.C., Donnelly P. 1997. Inferring Coalescence Times From DNA Sequence Data. Genetics. 145:505–518.

Than C., Ruths D., Nakhleh L. 2008. PhyloNet: a software package for analyzing and reconstructing reticulate evolutionary relationships. BMC Bioinf. 9:322.

Thórsson ÆH.Th., Salmela E., Anamthawat-Jónsson K. 2001. Morphological, cytogenetic, and molecular evidence for introgressive hybridization in birch. J. Hered. 92:404–408.

Truong C., Palmé A.E., Felber F. 2007. Recent invasion of the mountain birch *Betula pubescens* ssp. tortuosa above the treeline due to climate change: genetic and ecological study in northern Sweden. J. Evol. Biol. 20:369–380.

Tsuda Y., Semerikov V., Sebastiani F., Vendramin G.G., Lascoux M. 2017. Multispecies genetic structure and hybridization in the Betula genus across Eurasia. Mol. Ecol. 26:589–605.

Van de Peer Y., Mizrachi E., Marchal K. 2017. The evolutionary significance of polyploidy. Nat. Rev. Genet. 18:411–424.

Wagner N.D., He L., Hörandl E. 2020. Phylogenomic Relationships and Evolution of Polyploid Salix Species Revealed by RAD Sequencing Data. Front. Plant Sci. 11:1077.

Walters S.M. 1968. Betula L. in Britain. Pro. Bot. Soc. Br. Isl. 7: 179–180.

Wang N., Borrell J.S., Bodles W.J.A., Kuttapitiya A., Nichols R.A., Buggs R.J.A. 2014. Molecular footprints of the Holocene retreat of dwarf birch in Britain. Mol. Ecol. 23:2771–2782.

Wang N., McAllister H.A., Bartlett P.R., Buggs R.J.A. 2016. Molecular phylogeny and genome size evolution of the genus *Betula* (Betulaceae). Ann Bot. 117:1023–1035.

Wang Z., Miao H., Liu J., Xu B., Yao X., Xu C., Zhao S., Fang X., Jia C., Wang J., Zhang J., Li J., Xu Y., Wang J., Ma W., Wu Z., Yu L., Yang Y., Liu C., Guo Y., Sun S., Baurens F.-C., Martin G., Salmon F., Garsmeur O., Yahiaoui N., Hervouet C., Rouard M., Laboureau N., Habas R., Ricci S., Peng M., Guo A., Xie J., Li Y., Ding Z., Yan Y., Tie W., D’Hont A., Hu W., Jin Z. 2019. *Musa balbisiana* genome reveals subgenome evolution and functional divergence. Nat. Plants. 5:810–821.

Wang N., Kelly L.J., McAllister H.A., Zohren J., Buggs R.J.A. 2021. Resolving phylogeny and polyploid parentage using genus-wide genome-wide sequence data from birch trees. Mol. Phylogenet. Evol. 160:107126.

Wen D., Yu Y., Zhu J., Nakhleh L. 2018. Inferring phylogenetic networks using PhyloNet. Syst. Biol. 67:735–740.

Whitney K.D., Randell R.A., Rieseberg L.H. 2006. Adaptive Introgression of Herbivore Resistance Traits in the Weedy Sunflower *Helianthus annuus*. Am. Nat. 167:794–807.

Winkler M., Escobar García P., Gattringer A., Sonnleitner M., Hülber K., Schönswetter P., Schneeweiss G.M. 2017. A novel method to infer the origin of polyploids from Amplified Fragment Length Polymorphism data reveals that the alpine polyploid complex of *Senecio carniolicus* (Asteraceae) evolved mainly via autopolyploidy. Mol. Ecol. Resour. 17:877–892.

Wolfe K.H., Shields D.C. 1997. Molecular evidence for an ancient duplication of the entire yeast genome. Nature. 387:708–713.

Wood T.E., Takebayashi N., Barker M.S., Mayrose I., Greenspoon P.B., Rieseberg L.H. 2009. The frequency of polyploid speciation in vascular plants. Proc. Natl. Acad. Sci. U.S.A. 106:13875– 13879.

Wu Y., Lin F., Zhou Y., Wang J., Sun S., Wang B., Zhang Z., Li G., Lin X., Wang X., Sun Y., Dong Q., Xu C., Gong L., Wendel J.F., Zhang Z., Liu B. 2021. Genomic mosaicism due to homoeologous exchange generates extensive phenotypic diversity in nascent allopolyploids. Natl. Sci. Rev. 8:nwaa277.

Xiong Z., Gaeta R.T., Edger P.P., Cao Y., Zhao K., Zhang S., Pires J.C. 2021. Chromosome inheritance and meiotic stability in allopolyploid *Brassica napus*. G3: Genes Genomes Genet. 11:jkaa011.

Yan Z., Cao Z., Liu Y., Ogilvie H.A., Nakhleh L. 2022. Maximum Parsimony Inference of Phylogenetic Networks in the Presence of Polyploid Complexes. Syst. Biol. 71:706–720.

Yano R., Nakamura M., Yoneyama T., Nishida I. 2005. Starch-Related α-Glucan/Water Dikinase Is Involved in the Cold-Induced Development of Freezing Tolerance in Arabidopsis. Plant Physiol. 138:837–846.

Yant L., Hollister J.D., Wright K.M., Arnold B.J., Higgins J.D., Franklin F.C.H., Bomblies K. 2013. Meiotic Adaptation to Genome Duplication in *Arabidopsis arenosa*. Curr. Biol. 23:2151– 2156.

Yoshida K., Schuenemann V.J., Cano L.M., Pais M., Mishra B., Sharma R., Lanz C., Martin F.N., Kamoun S., Krause J., Thines M., Weigel D., Burbano H.A. 2013. The rise and fall of the *Phytophthora infestans* lineage that triggered the Irish potato famine. eLife. 2:e00731.

Zhang C., Ogilvie H.A., Drummond A.J., Stadler T. 2018. Bayesian Inference of Species Networks from Multilocus Sequence Data. Mol. Biol. Evol. 35:504–517.

Zhang T., Jing J.-L., Liu L., He Y. 2021. ZmRAD17 Is Required for Accurate Double-Strand Break Repair During Maize Male Meiosis. Front. Plant Sci. 12.

Zhou B.-B.S., Elledge S.J. 2000. The DNA damage response: putting checkpoints in perspective. Nature. 408:433–439.

Zohren J., Wang N., Kardailsky I., Borrell J.S., Joecker A., Nichols R.A., Buggs R.J.A. 2016. Unidirectional diploid–tetraploid introgression among British birch trees with shifting ranges shown by restriction site-associated markers. Mol. Ecol. 25:2413–2426.

