## Supplementary Materials 1 for "Complex Polyploids: Origins, Genomic Composition, and Role of Introgressed Alleles"

**DNA Isolation, Library Preparation and Sequencing: Detailed Description**

For the newly collected samples (samples not included in previous studies), total DNA was extracted from birch bud and leaf tissues using a DNeasy Plant Mini Kit (Qiagen AG, Düsseldorf, Germany) following the instructions provided by the manufacturer, apart from the incubation during lysis which was extended to 2 hours. Individual libraries were generated using the SeqCap EZ HyperCapWorkflow (Roche AG, Basel, Switzerland) using custom probes designed for targeted exome capture (TEC) and sequenced on paired-end mode (150 bp) on an Illumina NovaSeq 6000 platform (Illumina, San Diego, CA). Target resequencing was based on a custom probe design covering 1,609 nuclear genes with a 3.2M bp cumulative target size and reduced sequence redundancy. The probe set was identical to the one used in TEC sequencing of the extra 64 TEC accessions included in this paper and which were part of a previous study (Milesi et al. 2023). Nucleotide sequence similarity was checked with USEARCH, v. 11.0.667 (Edgar 2010), using a greedy clustering algorithm (-cluster_fast) and a 0.95 identity threshold.

Results from the clustering analysis show that, of the 1609 nucleotide sequences, 1599 (99.4%) are unique.

Eight additional birch accessions were obtained from The European Nucleotide Archive (ENA, https://www.ebi.ac.uk/ena). These include Illumina whole genome sequencing (WGS) libraries for *B. lenta*, *B. nana*, *B. occidentalis*, *B. pendula*, *B. platyphylla*, *B. populifolia*, and *B. pubescens* specimens, which were part of a previous study (Salojärvi et al., 2017). Two alder (*Alnus*) accessions, part of the same study, were also included as an outgroup. These accessions were used in the phylogenetic and/or population structure analyses. Information about each accession can be found in Supplementary Table S1 and *Supplementary Materials 2*.

Finally, TEC libraries for twenty *B. pendula* specimens, part of a previous study (Milesi et al. 2023), were downloaded from the ENA database. These samples were used in the population structure analysis. TEC was performed using the same probe set described above. ENA accession run codes are listed in *Supplementary Materials 2*.

**Read Mapping and Variant Calling: Detailed Description**

The pipeline used for read mapping and variant calling generally followed the Broad Institute best practices workflow (DePristo et al. 2011, Poplin et al. 2018). Sequenced reads were soft-clipped for Illumina adapters using MarkIlluminaAdapters from the Picard suite, v. 2.10.3 (https://broadinstitute.github.io/picard/), and mapped to the *B. pendula* genome assembly, v. Bpev01 (Salojärvi et al. 2017), using BWA-MEM v 0.7.17 (Li and Durbin 2009) and PCR and optical duplicates were flagged using MarkDuplicates from the PICARD. Chromosomal assignment of loci was performed on the basis of the *B. pendula* chromosomal map, v. 1.4c (https://genomevolution.org/CoGe/GenomeInfo.pl?gid=35080, accessed 2023-01-26). For samples sequenced using TEC, more than 94% (mean = 99.1%) of reads mapped to the *B.*

*pendula* genome (*Supplementary Materials 3)*. For WGS birch samples, at least 83% (mean = 92.8%) of sequenced reads mapped to the *B. pendula* reference assembly. On average, about 52% of reads associated to the two alder libraries mapped to the *B. pendula* genome.

Genotyping was performed with GATK 4.2.0.0 (McKenna et al., 2010) followed by site-level hard filtering (https://gatk.broadinstitute.org/hc/en-us/articles/360035890471-Hard-filtering-germline-short-variants, accessed 2022-07-14) using the filtering conditions described previously by Leal et al. (2023). For all birch and alder accessions, including those obtained through WGS, only sites located within the 1,603 nuclear loci covered by the TEC exome probes were considered (six of the targeted 1,609 loci had very low coverage and were excluded). Sites located outside the targeted regions were excluded during downstream analysis. Non-variant sites located within the targeted regions were also included in the output VCF file when preparing the datasets used in the phylogenetic analysis, but excluded when generating the datasets used in the analysis of population structure. See *Supplementary Materials 3* for further details.

**Classification of Polyploid Specimens: Detailed Description**

We classified each sample according to its ploidy level (2X, 3X, or 4X) by estimating the distribution of read counts at biallelic variant sites (Yoshida et al. 2013; Zohren et al. 2016). For each sample, the ratio between the number of reads supporting the reference and alternate alleles was computed for each biallelic heterozygous site using CollectAllelicCounts from the GATK suite. If the sample is diploid (2X), the reference and alternate alleles are expected to have similar read support levels, leading to a unimodal distribution of read frequencies centered on 0.5 (Supplementary Fig. S2b). For tetraploid samples (4X), the distribution of read counts is expected to show three modes – at 0.25, 0.5, and 0.75 – while triploid samples (3X) are foreseen to have two modes, at 0.33 and 0.66 (Supplementary Figs. S2a and S2c). The allelic frequency associated to the dominant mode, as well as the kurtosis value associated to the distribution, are shown for each sample in *Supplementary Materials 3*.

Tetraploid samples were further split into *B. pubescens* and 4X *B. pendula* by carrying out a phylogenetic analysis of each individual using the genomic polarization framework described in this paper. *B. pubescens* specimens contain genomic components of *B. nana* and *B. humilis* origin (see pairing profiles shown in Supplementary Figure S2a), while in *B. pendula* autotetraploids these are absent (Supplementary Fig. S2c). The same procedure can be used to distinguish between *B. pendula* triploids and *B. pendula* x *B. pubescens* triploid hybrids (Supplementary Figs. S2d and S2e). Ploidal and taxonomic classification of each of the samples included in this study, based on the approach just described, is shown in *Supplementary Materials 3*.

**Generation of Consensus Sequences and Multiple Sequence Alignments (MSA): Detailed Description**

Consensus sequences were generated for each accession using bcftools consensus v 1.12 (Danecek et al. 2021) by applying single-nucleotide polymorphisms (SNPs) specific to each individual to the *B. pendula* reference assembly. Heterozygotic sites were coded following the IUPAC nomenclature (Cornish-Bowden 1985). Deletions and failed variants were subsequently masked using maskfasta from bedtools v 2.29.2 (Quinlan and Hall 2010). Only sites located in the 1,603 targeted loci were included in the consensus sequence. As only invariant sites and point mutations were used when producing each consensus sequence, consensus sequences associated to different samples are aligned by default: they all have the same length and retain identical start positions for each gene, and therefore require no further alignment (Leal et al. 2023).

Separate MSAs were produced for each of the 1,603 loci included in the analysis by splitting each consensus sequence using getfasta from the bedtools suite, based on the gff3 annotation file associated to the *B. pendula* reference assembly (Salojärvi et al. 2017). Columns in the MSA with more than two masked sites were removed using trimAl v 1.4.1 (Capella-Gutiérrez et al. 2009). Each MSA contains eight fixed birch species (*B. albosinensis, B. humilis, B. lenta*, *B. nana*, *B. occidentalis*, *B. pendula*, *B. platyphylla*, and *B. populifolia*) and two alder species (*A. glutinosa* and *A. incana*). Aside from *B. albosinensis* and *B. humili*s, WGS accessions were always selected when generating the MSAs. Additionally, each MSA contains a nucleotide sequence associated to one of the 269 *B. pubescens* samples included in this study, with each *pubescens* specimen therefore being the subject of a separate phylogenetic analysis. In total, 431,207 (1,603 loci x 269 samples) MSAs were generated. We also prepared MSAs where the *pubescens* nucleotide sequence was replaced by one of the 25 tetraploid *B. pendula* accessions or one of the two triploid hybrids.

**Phylogenetic Analysis: Detailed Description**

Phylogenetic reconstruction of individual gene families was performed using maximum-likelihood as implemented in IQ-TREE2 v 2.0-rc2-omp-mpi (Minh et al. 2020) with 1000 ultrafast bootstrap replicates (Hoang et al. 2018) and substitution model selection executed using ModelFinder (Kalyaanamoorthy et al. 2017). MSAs were discarded and subsequently omitted during downstream analysis if IQ-TREE2 failed to converge, or if the MSA contained less than 100 parsimony-informative sites, over 50% of gaps or other ambiguities, or contained two or more identical nucleotide sequences.

Species trees were inferred using ASTRAL v 5.7.3 (Zhang et al. 2018), a super-tree inference method statistically consistent under the multi-species coalescent model, based on the phylogenies estimated previously for each individual gene family using IQ-TREE2. Options used while running IQ-TREE2 and ASTRAL were identical to those described in a previous paper (Leal et al. 2023).

Because phylogenetic inference was based on genomic data obtained using different sequencing approaches (either WGS or exome capture), a preliminary phylogenetic analysis was carried out for a special set of MSAs that included both WGS and TEC sequences for four birch species: *B. nana*, *B. pendula*, *B. platyphylla*, and *B. pubescens*. This was done as a control in order to ascertain whether the inclusion of libraries produced using disparate technologies is likely to induce biases or artifacts during phylogenetic reconstruction. The inferred species tree shows that WGS and TEC sequences associated to the same species always pair together, thus suggesting that no major drawbacks can be inferred from mixing WGS and TEC sequences during phylogenetic reconstruction (Supplementary Fig. S4). A similar tree topology was obtained when the analysis was rerun for different North-Scandinavian *B. pubescens* TEC sequences (data not shown).

**Population Structure Analysis: Detailed Description**

Bayesian statistical inference of admixture levels between *B. pubescens* populations and different birch species was performed with STRUCTURE v 2.3.4 (Pritchard et al. 2000), a model-based method suitable for the analysis of mixed ploidy datasets (Dufresne et al. 2014, Meirmans et al. 2018, Stift et al. 2019). As recommended by the authors, diploid samples were coded as if tetraploid but with the two extra alleles flagged as missing data. Structure and admixture inference were based on the analysis of 10,000 variant loci in 126 birch accessions. Sixteen independent runs were performed for each K value (number of clusters), with K ranging from 2 to 9. Parallelization was implemented using Structure_threader v. 1.3.10 (Pina-Martins et al. 2017). For each run, the Monte Carlo Markov Chain was run for 250,000 iterations after a burn-in period of 100,000 iterations. STRUCTURE was run using the following options: 'admixture model', 'correlated allele frequencies', and the 'LOCPRIOR model'. The latter allows sampling location to assist in the clustering process and its use is recommended when assessing the presence of structure among weakly divergent populations (Hubisz et al. 2009). Individual values for α, the relative admixture parameter, were inferred for each population. When in the presence of unbalanced sampling, allowing α values to vary across populations improves the accuracy in the estimation of the optimal K value during downstream analysis, as long as the number of clusters is not too large (< 20; Wang 2017). In order to further minimize the effect of unbalanced population sizes, the number of *B. pubescens* samples of Scandinavian origin was also downsampled. Finally, the λ value, the Dirichlet prior parameter used to model the distribution of allele frequencies, was also optimized as the (fixed) default value is often not appropriate for analyses using SNP data (Porras-Hurtado et al. 2013). The most likely number of ancestral populations was determined using both Evanno's ΔK statistic (Evanno et al. 2005) and the ln Pr(X|K) metric, the approximated likelihood of the data given K (Pritchard et al. 2000), averaged across replicates. Output results for the 16 replicates were combined for each K using StructureSelector (Li and Liu 2018) and Clumpp v. 1.1 (Jakobsson and Rosenberg 2007).

The presence of population genetic structure was also investigated using discriminant analysis of principal components (DAPC; Jombart et al. 2010) with *adegenet* v 2.1.7, an R package designed to perform multivariate analysis of genetic markers and which can handle mixed-ploidy datasets (Jombart 2008; Dufresne et al. 2014; Meirmans et al. 2018). In order to minimize within-cluster variance, DAPC first executes a data reduction step using principal component analysis (PCA) and then performs a discriminant analysis on the retained principal components (Jombart 2008). DAPC was performed on 50,000 variant loci belonging to 49 birch accessions. In order to minimize bias on PCA projections due to uneven sampling (McVean

2009), two to five samples were selected for each species/ploidy/location. The optimal number and membership of the clusters present in the dataset was determined using the 'find.clusters' function in *adegenet*, which performs a k-means clustering on the PCA transformed data. Results show that diploid and autotetraploid *B. pendula* samples cluster together (Fig. S11a), suggesting that both originate from a single gene pool (Meirmans et al. 2018). Samples obtained through WGS and TEC cluster together in all four species (*B. nana*, *B. pubescens*, *B. platyphylla* and *B. pendula*) for which both types of samples were available (Fig. S11a). Together with the phylogenetic tree obtained when both types of samples are included (Supplementary Fig. S4), these results provide further evidence that mixing libraries obtained using different sequencing approaches did not give rise to major biases or artifacts during downstream analyses.

Prior to running STRUCTURE and *adegenet*, private alleles were removed using the VariantFiltration tool in the GATK suite. Sites in linkage disequilibrium (LD) were removed using a modified version of scripts by Weitz and colleagues used to perform LD pruning in polyploid species (Weitz et al. 2021). Only variant sites within intronic regions (putatively neutral SNPs) and genotyped in all samples were included in the analysis.

**Coalescent Analysis: Detailed Description**

The approach used to determine the best evolutionary model using fastsimcoal2 (Excoffier et al. 2013), perform model optimization, and carry out final estimation of the confidence intervals (CI), involved three separate steps (Arnold et al. 2016).

I) First we built simple demographic models for all possible phylogenetic relationships between the three species included (3 models in total) and fitted them against the observed SFS data using fastsimcoal2. Each model was fit to SNP data using 50 independent runs of *fastsimcoal2*. At this stage, the only migration events allowed were the presence of gene flow from *B. pendula* to *B. pubescen*s. Comparison of model likelihoods, with the aim of determining which model has the highest probability of being correct given the candidate set of models, was performed using the Akaike Information Criterion (AIC), as described in Johnson and Omland (2004) and Arnold et al. (2016). In short, we started by computing the AIC value for model *i*, containing $d_i$ parameters and maximum estimated likelihood $MEL_i$, using:

$$AIC_i = 2d_i - 2ln(MEL_i) \qquad \text{(S1)}$$

These values are then used to estimate normalized Akaike weights (*w*) for each model using:

$$w_i = \left[\frac{e^{-0.5\Delta_i}}{\sum_k e^{-0.5\Delta_k}}\right] \qquad \text{(S2)}$$

with

$$\Delta_i = AIC_i - \min(\{AIC_1, \dots, AIC_N\}) \qquad \text{(S3)}$$

Akaike weights can be interpreted as the probability that model *i* is the one that best fits the data among the N models considered (Johnson and Omland 2004). Results are shown in Supplementary Table S3.

II) The best evolutionary model determined above was then further optimized by independently testing all possible migration combinations across species (129 models; 50 independent runs per model). As before, comparisons of model likelihoods using the AIC was used to determine the best overall model. Results are shown in *Supplementary Materials 5*.

III) Following the guidelines suggested by the authors (fastsimcoal2 v. 2.7 manual, http://cmpg.unibe.ch/software/fastsimcoal2/man/fastsimcoal27.pdf, last accessed 2023-05-09), estimation of confidence intervals was performed using parametric bootstrap. This is done by first simulating DNA sequences (and associated SFS) based on the parameters estimated for the best, optimized model. This process is repeated multiple times to create 100 replicates, and then 50 independent fastsimcoal2 runs are performed for each replicate. Finally, confidence intervals are computed based on the distributions obtained. Results are shown in Supplementary Table S4.

Next, examples of fastsimcoal2 input templates are provided for the model shown in Figure 5. The mutation rate used was taken from Salojärvi et al. 2017.

**fastsimcoal2 tpl input file:**

```
//Number of population samples (demes)
3 samples to simulate :
//Population effective sizes (number of genes)
PUBpopsize
PENDpopsize
PLATYpopsize
//Sample sizes
16
16
16
//Growth rates	: negative growth implies population expansion
0
0
0
//Number of migration matrices : 0 implies no migration between demes
3
//Migration matrix 0
0.0000  MIG0_pubpen  MIG0_pubpla
MIG0_penpub  0.0000  MIG0_penpla
MIG0_plapub  MIG0_plapen  0.0000
//Migration matrix 1
0.0000  MIG1_pubpen  0.0000
MIG1_penpub  0.0000  0.0000
0.0000  0.0000  0.0000
//Migration matrix 2
0.0000  0.0000  0.0000
0.0000  0.0000  0.0000
0.0000  0.0000  0.0000
//historical event: time, source, sink, migrants, new size, new growth rate, migr. matrix
2  historical event
TDIV1 2 1 1 ResizeTime1 0 1
TDIV2 0 1 1 ResizeTime2 0 2
//Number of independent loci [chromosome]
1 0
//Per chromosome: Number of linkage blocks
1
//per Block: data type, num loci, rec. rate and mut rate + optional parameters
FREQ 1 0 9.5e-9 OUTEXP
```

**fastsimcoal2 est input file:**

```
// Priors and rules file
// *********************

[PARAMETERS]
//#isInt? #name #dist.#min #max
//all N are in number of haploid individuals
1 PUBpopsize unif 1000 1e6 output
1 PENDpopsize unif 1000 1e6 output
1 PLATYpopsize unif 1000 1e6 output
1 ANCpopsize1 unif 1e4 5e6 output
1 ANCpopsize2 unif 1e4 5e6 output
1 TDIV1 unif  600 100000 output
1 TIMEextra unif  1 1000000 hide
0 MIG0_pubpen logunif 1e-7 1e-4 output
0 MIG0_pubpla logunif 1e-7 1e-4 output
0 MIG0_penpub logunif 0 0 output
0 MIG0_penpla logunif 1e-7 1e-4 output
0 MIG0_plapen logunif 1e-7 1e-4 output
0 MIG0_plapub logunif 1e-7 1e-4 output
0 MIG1_pubpen logunif 0 0 output
0 MIG1_penpub logunif 0 0 output

[COMPLEX PARAMETERS]
1 TDIV2 = TDIV1+TIMEextra output
0 ResizeTime1 = ANCpopsize1/PENDpopsize hide
0 ResizeTime2 = ANCpopsize2/ANCpopsize1 hide
0 Nm_pubpen0 = MIG0_pubpen*PUBpopsize output
0 Nm_pubpla0 = MIG0_pubpla*PUBpopsize output
0 Nm_penpub0 = MIG0_penpub*PENDpopsize output
0 Nm_penpla0 = MIG0_penpla*PENDpopsize output
0 Nm_plapen0 = MIG0_plapen*PLAYTpopsize output
0 Nm_plapub0 = MIG0_plapub*PLAYTpopsize output
0 Nm_pubpen1 = MIG1_pubpen*PUBpopsize output
0 Nm_penpub1 = MIG1_penpub*PENDpopsize output
```

### Geographic Distribution and Functional Analysis of Alleles of *B. nana* and *B. humilis* Ancestry: Detailed Description

K-means cluster analysis

K-means cluster analysis was used to partition *B. pubescens* loci containing alleles of *B. nana* or *B. humilis* ancestry in groups with a similar abundance profile across geographic locations. Clustering analysis was performed using the kmeans algorithm as implemented in MATLAB, v. 2019a (Mathworks), using correlation as the distance metric. The analysis was executed 50 times for each *k*, with different random initializations, and the within cluster dissimilarity value ($W_K$, the total sum of distances to the cluster centroids) was computed at each iteration and for each *k*. The optimal number of clusters, *k**, was inferred by determining the *k* at which the change in the cluster dissimilarity value across consecutive *k*-values ($W_K - W_{K+1}$) levels off (Hastie et al. 2009).

### References


Arnold B.J., Lahner B., DaCosta J.M., Weisman C.M., Hollister J.D., Salt D.E., Bomblies K., Yant L. 2016. Borrowed alleles and convergence in serpentine adaptation. *Proc. Natl. Acad. Sci. U.S.A*. 113:8320–8325.

Ashburner K., McAllister H.A. 2013. The genus Betula: a taxonomic revision of birches:26-28 (London: Kew Publishing).

Atkinson M.D. 1992. *Betula Pendula* Roth (*B. Verrucosa* Ehrh.) and *B. Pubescens* Ehrh. *J. Ecol.* 80:837–870.

Bauer J., Chen K., Hiltbunner A., Wehrli E., Eugster M., Schnell D., Kessler F. 2000. The major protein import receptor of plastids is essential for chloroplast biogenesis. *Nature*. 403:203–207.

Capella-Gutiérrez S., Silla-Martínez J.M., Gabaldón T. 2009. trimAl: a tool for automated alignment trimming in large-scale phylogenetic analyses. *Bioinformatics*. 25:1972–1973.

Caudullo G., Welk E., San-Miguel-Ayanz J. 2017. Chorological maps for the main European woody species. *Data Brief*. 12:662–666.

Cornish-Bowden A. 1985. Nomenclature for incompletely specified bases in nucleic acid sequences: recommendations 1984. *Nucleic Acids Res*. 13:3021–3030.

Danecek P., Bonfield J.K., Liddle J., Marshall J., Ohan V., Pollard M.O., Whitwham A., Keane T., McCarthy S.A., Davies R.M., Li H. 2021. Twelve years of SAMtools and BCFtools. *GigaScience*. 10.

DePristo M.A., Banks E., Poplin R., Garimella K.V., Maguire J.R., Hartl C., Philippakis A.A., del Angel G., Rivas M.A., Hanna M., McKenna A., Fennell T.J., Kernytsky A.M., Sivachenko A.Y., Cibulskis K., Gabriel S.B., Altshuler D., Daly M.J. 2011. A framework for variation discovery and genotyping using next-generation DNA sequencing data. *Nat. Genet.* 43:491–498.

Dufresne F., Stift M., Vergilino R., Mable B.K. 2014. Recent progress and challenges in population genetics of polyploid organisms: an overview of current state-of-the-art molecular and statistical tools. *Mol. Ecol*. 23:40–69.

Edgar R.C. 2010. Search and clustering orders of magnitude faster than BLAST. *Bioinformatics*. 26:2460–2461.

Evanno G., Regnaut S., Goudet J. 2005. Detecting the number of clusters of individuals using the software structure: a simulation study. *Mol. Ecol.* 14:2611–2620.

Excoffier L., Dupanloup I., Huerta-Sánchez E., Sousa V.C., Foll M. 2013. Robust Demographic Inference from Genomic and SNP Data. *PLOS Genet*. 9:e1003905.

Hastie T., Tibshirani R., Friedman, J.H. 2009. *The elements of statistical learning: Data mining, inference, and prediction*, 2nd ed. (New York: Springer).

Hoang D.T., Chernomor O., von Haeseler A., Minh B.Q., Vinh L.S. 2018. UFBoot2: Improving the Ultrafast Bootstrap Approximation. *Mol. Biol. Evol.* 35:518–522.

Hu Y.-N., Zhao L., Buggs R.J.A., Zhang X.-M., Li J., Wang N. 2019. Population structure of *Betula albosinensis* and *Betula platyphylla*: evidence for hybridization and a cryptic lineage. *Ann. Bot.* 123:1179–1189.

Hubisz M.J., Falush D., Stephens M., Pritchard J.K. 2009. Inferring weak population structure with the assistance of sample group information. *Mol. Ecol. Resour*. 9:1322–1332.

Jakobsson M., Rosenberg N.A. 2007. CLUMPP: a cluster matching and permutation program for dealing with label switching and multimodality in analysis of population structure. *Bioinformatics*. 23:1801–1806.

Johnson J.B., Omland K.S. 2004. Model selection in ecology and evolution. *Trends Ecol. Evol.* 19:101–108.

Jombart T. 2008. adegenet: a R package for the multivariate analysis of genetic markers. *Bioinformatics*. 24:1403–1405.

Jombart T., Devillard S., Balloux F. 2010. Discriminant analysis of principal components: a new method for the analysis of genetically structured populations. *BMC Genet.* 11:1–15.

Kalyaanamoorthy S., Minh B.Q., Wong T.K.F., von Haeseler A., Jermiin L.S. 2017. ModelFinder: fast model selection for accurate phylogenetic estimates. *Nat. Methods.* 14:587–589.

Leal J.L., Milesi P., Salojärvi J., Lascoux M. 2023. Phylogenetic Analysis of Allotetraploid Species Using Polarized Genomic Sequences. *Syst. Biol.* 72:372–390.

Li H., Durbin R. 2009. Fast and accurate short read alignment with Burrows–Wheeler transform. *Bioinformatics*. 25:1754–1760.

Li Y.-L., Liu J.-X. 2018. StructureSelector: A web-based software to select and visualize the optimal number of clusters using multiple methods. *Mol. Ecol. Resour.* 18:176–177.

McKenna A., Hanna M., Banks E., Sivachenko A., Cibulskis K., Kernytsky A., Garimella K., Altshuler D., Gabriel S., Daly M., DePristo M.A. 2010. The Genome Analysis Toolkit: A MapReduce framework for analyzing next-generation DNA sequencing data. *Genome Res.* 20:1297–1303.

McVean G. 2009. A Genealogical Interpretation of Principal Components Analysis. *PLoS Genet.* 5:e1000686.

Meirmans P.G., Liu S., van Tienderen P.H. 2018. The Analysis of Polyploid Genetic Data. *J. Hered.* 109:283–296.

Milesi P., Kastally C., Dauphin B., Cervantes S., Bagnoli F., Budde K.B., Cavers S., Ojeda D.I., Fady B., Faivre-Rampant P., González-Martínez S.C., Grivet D., Gugerli F., Jorge V., Lesur-Kupin I., Olsson S., Opgenoorth L., Pinosio S., Plomion C., Rellstab C., Rogier O., Scalabrin S., Scotti I., Vendramin G.G., Westergren M., Consortium G., Lascoux M., Pyhäjärvi T. 2023.

Synchronous effective population size changes and genetic stability of forest trees through glacial cycles. bioRxiv:2023.01.05.522822.

Minh B.Q., Schmidt H.A., Chernomor O., Schrempf D., Woodhams M.D., von Haeseler A., Lanfear R. 2020. IQ-TREE 2: new models and efficient methods for phylogenetic inference in the genomic era. *Mol. Biol. Evol.* 37:1530–1534.

Pina-Martins F., Silva D.N., Fino J., Paulo O.S. 2017. Structure_threader: An improved method for automation and parallelization of programs structure, fastStructure and MavericK on multicore CPU systems. *Mol. Ecol. Resour.* 17:e268–e274.

Poplin R., Ruano-Rubio V., DePristo M.A., Fennell T.J., Carneiro M.O., Auwera G.A.V. der, Kling D.E., Gauthier L.D., Levy-Moonshine A., Roazen D., Shakir K., Thibault J., Chandran S., Whelan C., Lek M., Gabriel S., Daly M.J., Neale B., MacArthur D.G., Banks E. 2018. Scaling accurate genetic variant discovery to tens of thousands of samples. bioRxiv.:201178.

Porras-Hurtado L., Ruiz Y., Santos C., Phillips C., Carracedo Á., Lareu M. 2013. An overview of STRUCTURE: applications, parameter settings, and supporting software. *Front. Genet.* 4.

Pritchard J.K., Stephens M., Donnelly P. 2000. Inference of Population Structure Using Multilocus Genotype Data. *Genetics.* 155:945–959.

Quinlan A.R., Hall I.M. 2010. BEDTools: a flexible suite of utilities for comparing genomic features. *Bioinformatics.* 26:841–842.

Salojärvi J., Smolander O.-P., Nieminen K., Rajaraman S., Safronov O., Safdari P., Lamminmäki A., Immanen J., Lan T., Tanskanen J., Rastas P., Amiryousefi A., Jayaprakash B., Kammonen J.I., Hagqvist R., Eswaran G., Ahonen V.H., Serra J.A., Asiegbu F.O., de Dios Barajas-Lopez J., Blande D., Blokhina O., Blomster T., Broholm S., Brosché M., Cui F., Dardick C., Ehonen S.E., Elomaa P., Escamez S., Fagerstedt K.V., Fujii H., Gauthier A., Gollan P.J., Halimaa P., Heino P.I., Himanen K., Hollender C., Kangasjärvi S., Kauppinen L., Kelleher C.T., Kontunen-Soppela S., Koskinen J.P., Kovalchuk A., Kärenlampi S.O., Kärkönen A.K., Lim K.-J., Leppälä J., Macpherson L., Mikola J., Mouhu K., Mähönen A.P., Niinemets Ü., Oksanen E., Overmyer K., Palva E.T., Pazouki L., Pennanen V., Puhakainen T., Poczai P., Possen B.J.H.M., Punkkinen M., Rahikainen M.M., Rousi M., Ruonala R., van der Schoot C., Shapiguzov A., Sierla M., Sipilä T.P., Sutela S., Teeri T.H., Tervahauta A.I., Vaattovaara A., Vahala J., Vetchinnikova L., Welling A., Wrzaczek M., Xu E., Paulin L.G., Schulman A.H., Lascoux M., Albert V.A., Auvinen P., Helariutta Y., Kangasjärvi J. 2017. Genome sequencing and population genomic analyses provide insights into the adaptive landscape of silver birch. *Nat. Genet.* 49:904–912. https://doi.org/10.1038/ng.3862

Stift M., Kolář F., Meirmans P.G. 2019. STRUCTURE is more robust than other clustering methods in simulated mixed-ploidy populations. *Heredity*. 123:429–441.

Wang J. 2017. The computer program structure for assigning individuals to populations: easy to use but easier to misuse. *Mol. Ecol. Resour.* 17:981–990.

Weitz A.P., Dukic M., Zeitler L., Bomblies K. 2021. Male meiotic recombination rate varies with seasonal temperature fluctuations in wild populations of autotetraploid *Arabidopsis arenosa*. *Mol. Ecol*. 30:4630–4641.

Yoshida K., Schuenemann V.J., Cano L.M., Pais M., Mishra B., Sharma R., Lanz C., Martin F.N., Kamoun S., Krause J., Thines M., Weigel D., Burbano H.A. 2013. The rise and fall of the *Phytophthora infestans* lineage that triggered the Irish potato famine. *eLife*. 2:e00731.

Zhang C., Rabiee M., Sayyari E., Mirarab S. 2018. ASTRAL-III: polynomial time species tree reconstruction from partially resolved gene trees. *BMC Bioinf.* 19:153.

Zohren J., Wang N., Kardailsky I., Borrell J.S., Joecker A., Nichols R.A., Buggs R.J.A. 2016. Unidirectional diploid–tetraploid introgression among British birch trees with shifting ranges shown by restriction site-associated markers. *Mol. Ecol*. 25:2413–2426.

Table S1 | **Sample information (*Betula* and *Alnus*)**. Specie's name, number of individuals sequenced, sequencing technology, ploidy, number of chromosomes found in somatic cells, and geographic distribution. Betula genus's base chromosome number is x=14. See *Supplementary Materials 2* for ENA accession number for each sample.

| Genus | Species | Number of individuals | Sequencing | Ploidy | Chromosome No. | Distribution | Source[a] |
|---|---|---|---|---|---|---|---|
| *Betula* | *B. albosinensis* Burk. | 1 | exome capture | tetraploid | 2n=4x=56[b] | Asia | BBG |
| | *B. humilis* Schrk. | 2 | exome capture | diploid | 2n=2x=28 | Asia, Europe | NP |
| | *B. lenta* L. | 1 | WGS | diploid | 2n=2x=28 | North America | HBG (JS) |
| | *B. nana* L. | 1 | WGS | diploid | 2n=2x=28 | Asia, Europe, N. America | FFR (JS) |
| | | 3 | exome capture | diploid | 2n=2x=28 | Asia, Europe, N. America | NP |
| | *B. occidentalis* Hook | 1 | WGS | diploid | 2n=2x=28 | North America | HBG (JS) |
| | *B. pendula* Roth. | 2 | WGS | diploid | 2n=2x=28 | Asia, Europe | NP (JS) |
| | | 20 | exome capture | diploid | 2n=2x=28 | Asia, Europe | NP (PM) |
| | *B. pendula* (4X) | 25 | exome capture | tetraploid | 2n=4x=56 | | NP (PM) |
| | *B. pendula* (3X) | 1 | exome capture | triploid | 2n=3x=42 | | NP (PM) |
| | *B. platyphylla* Suk. | 1 | WGS | diploid | 2n=2x=28 | Asia | HBG (JS) |
| | | 15 | exome capture | diploid | 2n=2x=28 | Asia | NP |
| | *B. populifolia* Marsh. | 1 | WGS | diploid | 2n=2x=28 | North America | HBG (JS) |
| | *B. pubescens* Ehrh. | 1 | WGS | tetraploid | 2n=4x=56 | Asia, Europe | HBG (JS) |
| | | 269 | exome capture | tetraploid | 2n=4x=56 | Asia, Europe | NP |
| | *B. pendula* x *B. pubescens* | 1 | exome capture | triploid | 2n=3x=42 | | NP |
| *Alnus* (outgroup) | *A. glutinosa* | 1 | WGS | | | | NP (JS) |
| | *A. incana* | 1 | WGS | | | | NP (JS) |

[a] BBG = Bergius Botanic Garden (Sweden); FFR = The Finnish Research Institute (Finland); HBG = Helsinki Botanical Gardens (Finland); JS = Salojärvi et al. (2017); NP = natural population in Europe; PM = Milesi et al. (2023)
[b] Both diploid and tetraploid specimens have been documented (Hu et al. 2019)

Table S2 | **Optimized genomic composition in the presence of homoeologous replacement –** *B. pubescens* genomic composition as predicted by each evolutionary model, in the absence (**a**) or presence (**b**) of homoeologous replacement. P, N, H, $P_y$, and *A* indicate the hypothetical evolutionary origin of each subgenomic component in the tetraploid (*B. pendula*, *B. nana*, *B. humilis*, *B. platyphylla*, and common ancestor of *B. pendula* and *B. platyphylla*, respectively). Modeling data summarizes optimized results [top 5% simulation runs (50 out of 1000 runs)] obtained using approximate Bayesian computation having as a reference the values observed experimentally for a population in Southern Sweden. "1+ copies": percentage of loci with at least one gene copy originating from, respectively, *B. nana*, *B. humilis*, *B. pendula*, or the common ancestor of *B. pendula* and *B. platyphylla*. "4 copies": percentage of loci where all four gene copies originate from one of the subgenomes via homoeologous replacement. Weighted *L2* norm distance, averaged over the top 5% simulation run, is also shown for each model. Values for AAAA model (best model in the absence of homoeologous replacement) are highlighted in green. Total levels of homoeologous replacement are highlighted in lilac (includes only values for gene copies originating from *B. pendula* or from the ancestor of *B. pendula* and *B. platyphylla*).

a)

| MODEL | No Homoeologous Replacement | | | | | | | |
|---|---|---|---|---|---|---|---|---|
| | L2 norm | 1+ copies | | | | All 4 copies | | |
| | | nana | humilis | pendula | ancestral | pendula | ancestral | Total |
| 1 PPPP | 0.74 ± 0.01 | 0.42 ± 0.03 | 0.36 ± 0.02 | 0.97 ± 0.01 | - | 0.35 ± 0.03 | - | 0.35 |
| 2 AAAA | **0.49 ± 0.01** | 0.41 ± 0.03 | 0.34 ± 0.02 | 0.79 ± 0.02 | 0.82 ± 0.02 | 0.13 ± 0.02 | 0.17 ± 0.02 | **0.30** |
| 3 PPNN | 1.58 ± 0.01 | 0.53 ± 0.02 | 0.50 ± 0.03 | 0.98 ± 0.01 | - | - | - | - |
| 4 PPHH | 1.58 ± 0.01 | 0.56 ± 0.03 | 0.48 ± 0.02 | 0.98 ± 0.01 | - | - | - | - |
| 5 PPPyPy | 1.07 ± 0.02 | 0.49 ± 0.04 | 0.40 ± 0.03 | 0.95 ± 0.01 | - | - | - | - |
| 6 PPNH | 1.59 ± 0.01 | 0.60 ± 0.03 | 0.48 ± 0.02 | 0.91 ± 0.03 | - | - | - | - |
| 7 AANN | 1.50 ± 0.01 | 0.53 ± 0.02 | 0.51 ± 0.02 | 0.88 ± 0.02 | 0.98 ± 0.01 | - | - | - |
| 8 AAHH | 1.55 ± 0.01 | 0.55 ± 0.03 | 0.48 ± 0.02 | 0.81 ± 0.02 | 0.98 ± 0.01 | - | - | - |
| 9 AANH | 1.58 ± 0.01 | 0.50 ± 0.03 | 0.54 ± 0.02 | 0.81 ± 0.02 | 0.89 ± 0.01 | - | - | - |

b)

| MODEL | With Homoeologous Replacement (allopolyploid models only) | | | | | | | | | | |
|---|---|---|---|---|---|---|---|---|---|---|---|
| | L2 norm | 1+ copies | | | | All 4 copies (homoeologous replacement) | | | | | |
| | | nana | humilis | pendula | ancestral | nana | humilis | platyphylla | pendula | ancestral | Total |
| 1 PPPP | - | - | - | - | - | - | - | - | - | - | - |
| 2 AAAA | - | - | - | - | - | - | - | - | - | - | - |
| 3 PPNN | **0.76 ± 0.02** | 0.41 ± 0.04 | 0.35 ± 0.02 | 0.95 ± 0.02 | - | 0.01 ± 0.01 | - | - | 0.37 ± 0.02 | - | 0.37 |
| 4 PPHH | **0.76 ± 0.02** | 0.58 ± 0.05 | 0.38 ± 0.02 | 0.97 ± 0.01 | - | - | 0.01 ± 0.01 | - | 0.34 ± 0.05 | - | 0.34 |
| 5 PPPyPy | **0.66 ± 0.02** | 0.40 ± 0.03 | 0.33 ± 0.02 | 0.92 ± 0.03 | - | - | - | 0.02 ± 0.01 | 0.29 ± 0.02 | - | 0.29 |
| 6 PPNH | **0.75 ± 0.02** | 0.35 ± 0.04 | 0.31 ± 0.03 | 0.94 ± 0.05 | - | 0.01 ± 0.01 | 0.03 ± 0.01 | - | 0.39 ± 0.03 | - | 0.39 |
| 7 AANN | **0.55 ± 0.02** | 0.35 ± 0.03 | 0.36 ± 0.02 | 0.80 ± 0.02 | 0.80 ± 0.02 | 0.02 ± 0.01 | - | - | 0.17 ± 0.02 | 0.17 ± 0.02 | 0.34 |
| 8 AAHH | **0.53 ± 0.02** | 0.39 ± 0.04 | 0.31 ± 0.03 | 0.77 ± 0.02 | 0.78 ± 0.03 | - | 0.02 ± 0.01 | - | 0.18 ± 0.02 | 0.18 ± 0.02 | 0.36 |
| 9 AANH | **0.57 ± 0.02** | 0.41 ± 0.04 | 0.34 ± 0.03 | 0.74 ± 0.03 | 0.76 ± 0.03 | 0.01 ± 0.01 | 0.02 ± 0.01 | - | 0.17 ± 0.02 | 0.18 ± 0.02 | 0.35 |

Table S3 | **Likelihood analysis of possible phylogenies**. Likelihood (*Lhood*) analysis for different phylogenetic topologies and *B. pubescens* populations estimated using *fastsimcoal2*. Each model was fit to SNP data using 50 runs of *fastsimcoal2*. Possibility of gene flow from *B. pendula* to *B. pubescen*s is included in all models. Akaike information criteria (AIC), AIC variation versus best model (Δ), and Akaike weights (**w**) also shown. AIC analysis is performed independently for each *B. pubescens* population. Akaike weights show probability that model *i* is the best model for the observed data, given the candidate set of models.

| Population | Phylogeny (model) | Max $\log_{10}(\text{Lhood}_i)$ | $AIC_i$ | $\Delta_i$ | $\boldsymbol{w}_i$ | Notes |
|---|---|---|---|---|---|---|
| Central Europe | ((PEN, PUB), PLA) | -28527.1 | 65702.07521 | 698.6043172 | 1.9953E-152 | |
| | **((PEN, PLA), PUB)** | **-28223.7** | **65003.47089** | **0** | **1** | **best model** |
| | (PEN, (PUB, PLA)) | -28435.7 | 65491.61893 | 488.1480397 | 1.0000E-106 | |
| Southern Sweden | ((PEN, PUB), PLA) | -31814.8 | 73272.28422 | 351.8350022 | 3.98107E-77 | |
| | **((PEN, PLA), PUB)** | **-31662** | **72920.44921** | **0** | **1** | **best model** |
| | (PEN, (PUB, PLA)) | -31810.8 | 73263.07388 | 342.6246618 | 3.98107E-75 | |
| Central Asia | ((PEN, PUB), PLA) | -34656.7 | 79816.00079 | 705.281814 | 7.0795E-154 | |
| | **((PEN, PLA), PUB)** | **-34350.4** | **79110.71898** | **0** | **1** | **best model** |
| | (PEN, (PUB, PLA)) | -34670.7 | 79848.23698 | 737.5180053 | 7.0795E-161 | |
| Spain | ((PEN, PUB), PLA) | -49845 | 114788.354 | 372.7885266 | 1.12202E-81 | |
| | **((PEN, PLA), PUB)** | **-49683.1** | **114415.5654** | **0** | **1** | **best model** |
| | (PEN, (PUB, PLA)) | -49867 | 114839.0108 | 423.4453986 | 1.12202E-92 | |

Table S4 | **Maximum likelihood estimates (MLE) for parameters associated to evolutionary model with highest Akaike weight**. Parameters estimated using *fastsimcoal2* for the ((*B. pendula*, *B. platyphylla*), *B. pubescens*) phylogenetic topology, with the *B. pubescens* population containing individuals sampled from Central Europe. Upper and lower confidence intervals (95%) were estimated using parametric bootstrap based on 100 replicates, and then performing inference on each replicate based on 50 runs with *fastsimcoal2*. Population sizes are $2N_e$ or $4N_e$ haploid numbers for diploid and tetraploid species, respectively. Divergence times are shown in number of generations. The mean number of individuals in the sink population that migrated from the source population each generation was computed by multiplying the population size by the migration probability. Only *B. pubescens* loci containing four gene copies of *B. pendula/B. platyphylla* ancestry were included in the analysis (loci containing one or more gene copies originating from *B. nana* or *B. humilis* were excluded).

| | MLEs | Lower 95% CI | Upper 95% CI | Notes |
|---|---|---|---|---|
| PUBpopsize | 628,719 | 614,818 | 642,620 | *B. pubescens* population size ($4N_e$) |
| PENDpopsize | 178,164 | 174,431 | 181,896 | *B. pendula* population size ($2N_e$) |
| PLATYpopsize | 95,847 | 93,734 | 97,959 | *B. platyphylla* population size ($2N_e$) |
| ANCpopsize1 | 381,235 | 356,410 | 406,061 | Ancestral population size of common ancestor of *B. pendula* and *B. platyphylla* |
| ANCpopsize2 | 132,634 | 129,624 | 135,645 | Ancestral population size of common ancestor of all three species |
| TDIV1 | 100,120 | 97,821 | 102,419 | *B. pendula* / *B. platyphylla* divergence time |
| **TDIV2** | **183,594** | **178,659** | **188,529** | **Divergence time between *B. pubescens* and common ancestor of *B. pendula* and *B. platyphylla*** |
| MIG0_pubpen | 1.16E-05 | 1.13E-05 | 1.19E-05 | Backwards migration probability from B. *pubescens* to *B. pendula* |
| MIG0_pubpla | 7.72E-07 | 7.08E-07 | 8.36E-07 | Backwards migration probability from *B. pubescens* to *B. platyphylla* |
| MIG0_penpub | set to zero | | | Backwards migration probability from *B. pendula* to *B. pubescens* |
| MIG0_penpla | 1.24E-06 | 1.12E-06 | 1.35E-06 | Backwards migration probability from *B. pendula* to *B. platyphylla* |
| MIG0_plapen | 3.38E-06 | 3.07E-06 | 3.68E-06 | Backwards migration probability from *B. platyphylla to B. pendula* |
| MIG0_plapub | 5.90E-06 | 5.44E-06 | 6.36E-06 | Backwards migration probability from *B. platyphylla* to *B. pubescens* |
| MIG1_pubpen | set to zero | | | Backwards migration probability from *B. pubescens* to common ancestor of *B. pendula/B. platyphylla* |
| MIG1_penpub | set to zero | | | Backwards migration probability from common ancestor of *B. pendula/B. platyphylla* to *B. pubescens* |
| Nm_pubpen0 | 6.40079 | 6.16533 | 6.63625 | MIG0_pubpen * PUBpopsize |
| Nm_pubpla0 | 0.420072 | 0.387006 | 0.453139 | MIG0_pubpla * PUBpopsize |
| Nm_penpub0 | set to zero | | | MIG0_penpub * PENDpopsize |
| Nm_penpla0 | 0.185597 | 0.169749 | 0.201445 | MIG0_penpla * PENDpopsize |
| Nm_plapen0 | 0.206294 | 0.190474 | 0.222115 | MIG0_plapen * PLATYpopsize |
| Nm_plapub0 | 0.345092 | 0.32922 | 0.360963 | MIG0_plapub * PLATYpopsize |
| Nm_pubpen1 | set to zero | | | MIG1_pubpen * PUBpopsize |
| Nm_penpub1 | set to zero | | | MIG1_penpub * ANCpopsize1 |

Table S5 | ***B. pubescens*** **loci containing exclusively alleles inherited from** ***B. pendula*** **or from the common ancestor of** ***B. pendula*** **and** ***B. platyphylla***.

| Locus | *A. thaliana* homolog | Function |
|---|---|---|
| **Loci containing exclusively alleles inherited from the common ancestor of *B. pendula* and *B. platyphylla*** | | |
| Bpev01.c0752.g0012 | TOC159 Translocase of chloroplast 159 | Required for chloroplast biogenesis (Bauer et al. 2000) |
| **Loci containing exclusively alleles inherited from *B. pendula*** | | |
| Bpev01.c0223.g0009 | At2g37230 Pentatricopeptide repeat-containing protein | chloroplast protein |
| Bpev01.c0017.g0022 | At1g15240 | uncharacterized protein |

Figure S1 | ***B. pubescens* sampling locations** – Sampling locations (●) and *B. pubescens* natural distribution range. Species range adapted from Caudullo et al. (2017), Atkinson (1992), and Ashburner and McAllister (2013).

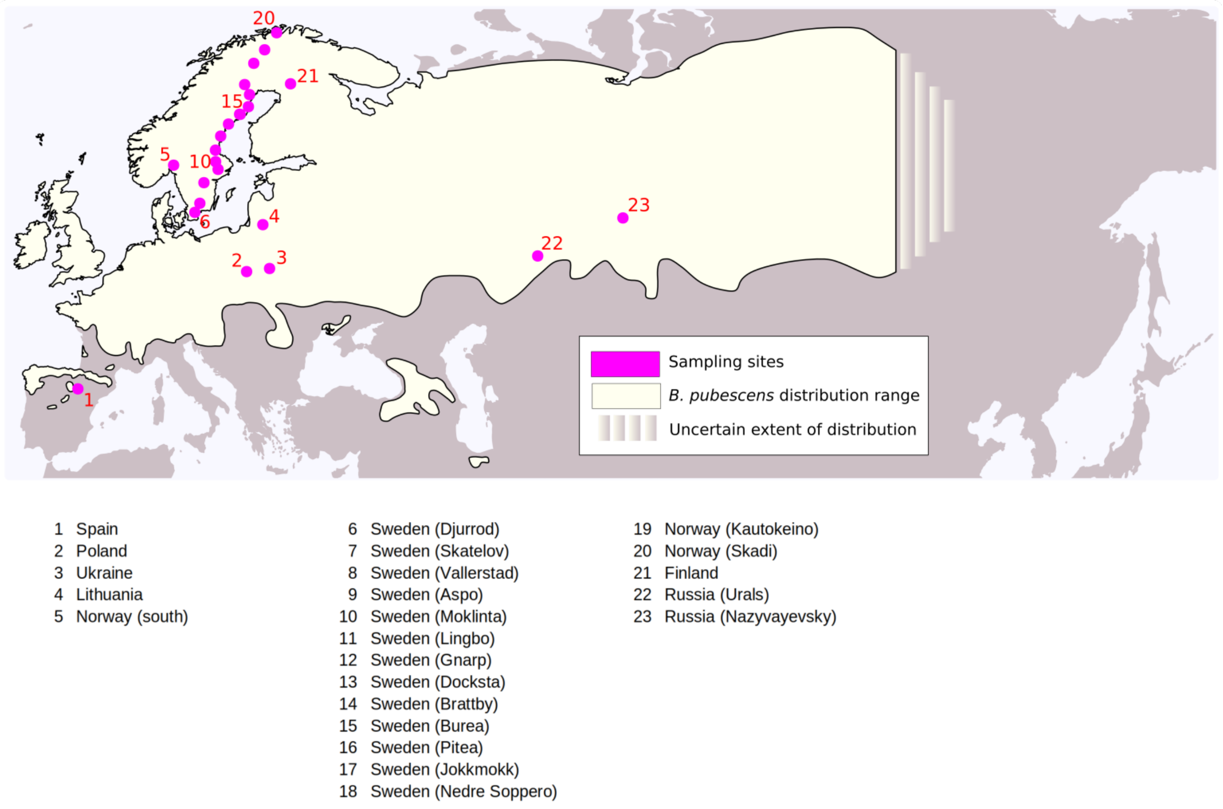

Figure S2 | **Classification of birch specimens** – Read frequency distributions for biallelic variants (left-most column) and pairing patterns for focal species (two right-most columns) for five birch samples: **(a)** *B. pubescens*; **(b)** diploid *B. pendula*; **(c)** autotetraploid *B. pendula*; **(d)** triploid *B. pendula*; and **(e)** *B. pendula* x *B. pubescens* triploid hybrid. **Barplots** on the right side show frequency with which the focal birch specimen pairs with other species and is based on phylogenetic analysis of individual gene families using IQ-TREE2. Identity of the reference sequence used during genomic polarization of focal birch species shown atop each of the two right-most columns.

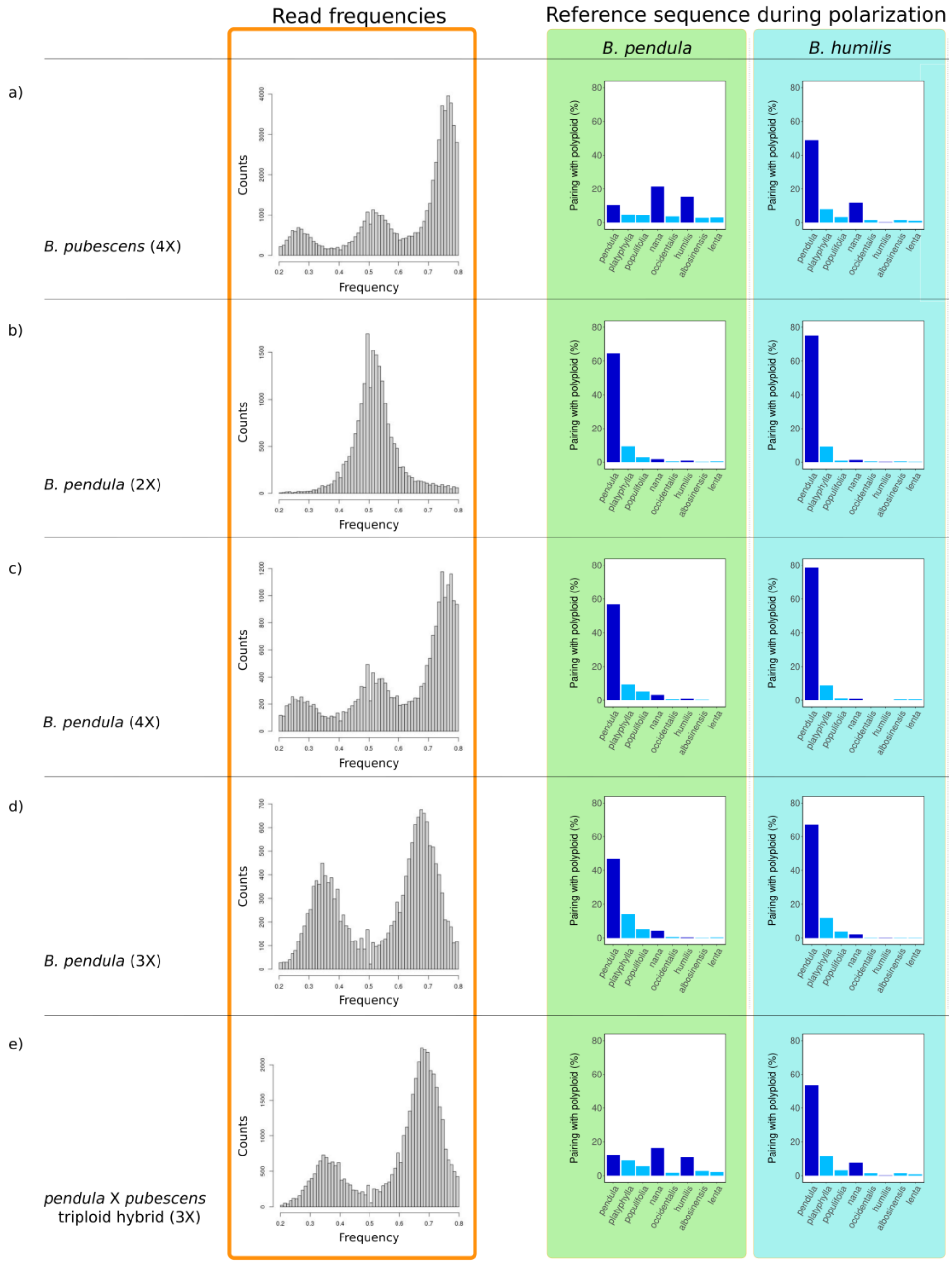

Figure S3 | **Bioinformatics pipeline** – Synoptic diagram displaying pipeline developed to produced observed pairing profiles per population and polarizing geometry (left panel) and perform polyploidization model optimization (right panel). Polarization geometry refers to reference sequence used during polarization (one of *B. pendula*, *B. platyphylla*, *B. nana*, or *B. humilis*). During polyploidization model optimization, comparison between simulated and observed pairing profiles (using the L2 norm distance) is used to find the best model parameters and estimate introgression levels. Initial model optimization is performed using simulated annealing. Final model optimization performed using an approximate Bayesian computation (ABC) rejection algorithm with priors sampled from distribution centered on model parameters estimated using simulated annealing. Optimization was done separately for each polyploidization model and *B. pubescens* population. Nine different polyploidization models were tested: PPPP, AAAA, PPNN, PPHH, PPPyPy, PPNH, AANN, AAHH, and AANH. *B. pubescens* populations modeled: Arctic, Northern Sweden, Southern Sweden, Central Asia, Lithuania, Ukraine, and Southwestern Europe.

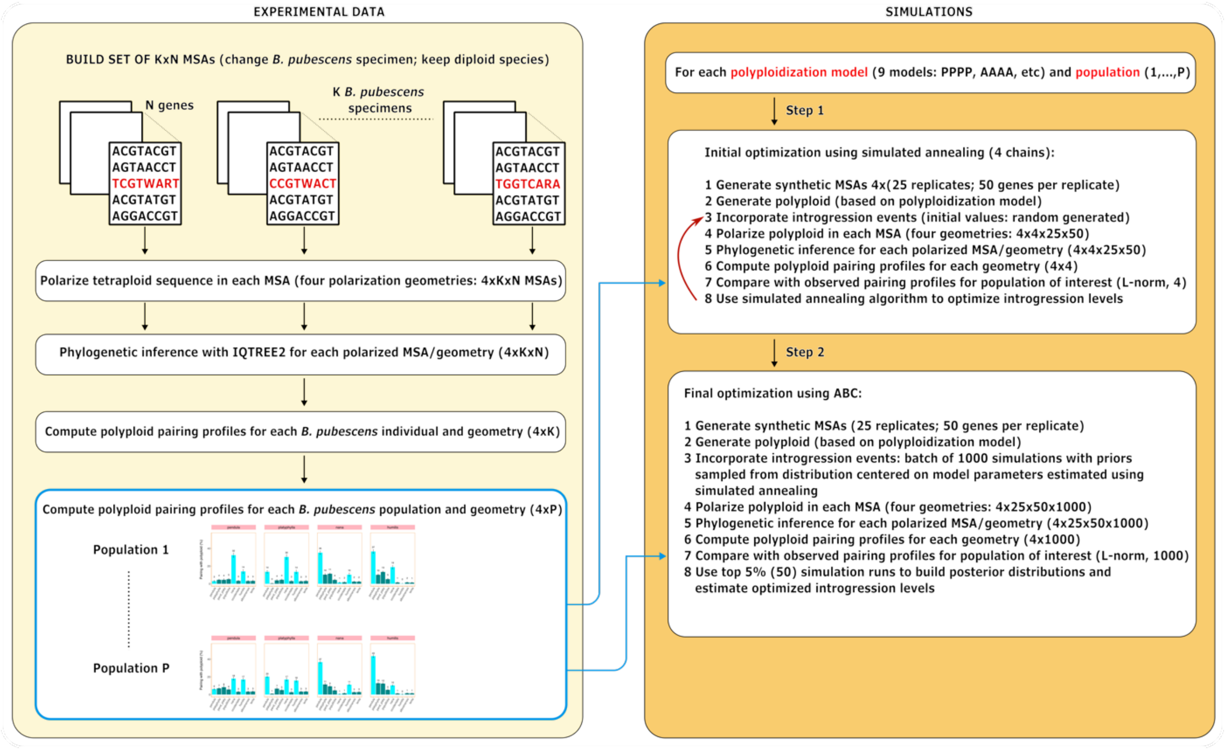

Figure S4 | **Birch phylogeny based on WGS and TEC sequences** – Species tree cladogram for 11 Betulaceae species. Four birch species (*B. nana*, *B. pendula*, *B. platyphylla*, and *B. pubescens*) are represented by both whole genome sequencing (WGS) and targeted exome capture (TEC) accessions. WGS and TEC sequences pair together in all four cases. Phylogenies obtained using ASTRAL based on 490 gene families (exonic sequences only). Pie charts show quartet support for each branch. Precise support values are shown for the four focal clades. Both *B. pubescens* specimens are from northern Finland (see *Supplementary Materials 2*). *B. pubescens* sequences were polarized using *B. humilis* as the reference sequence (●). Only the *B. pubescens* sequences are polarized. Sample ID provided after species name.

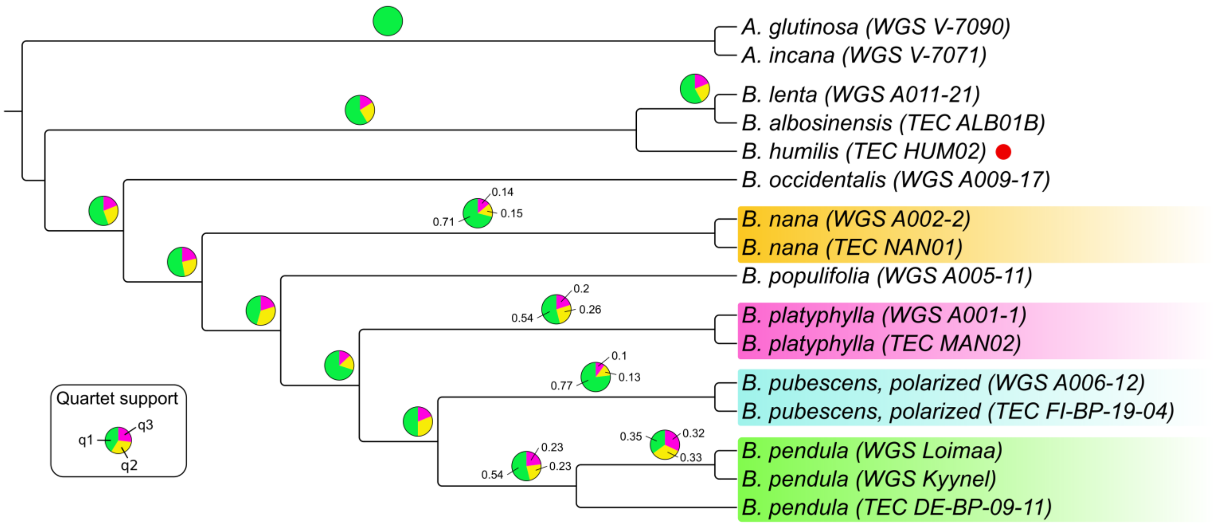

Figure S5 | **SYMPHY input phylogeny used in the simulation of the birch genus.** Example of input tree, mimicking the *Betulaceae* phylogeny, used for testing different evolutionary models for *B. pubescens*, based on different parental species (●) and levels of introgressive hybridization between *B. pubescens* and a third species (○). Phylogenetic position of parental species and hybridizing species vary according to the model. *Note*: SIMPHY requires starting trees to be ultrametric, that is, the distance from the root must be the same for every tree tip.

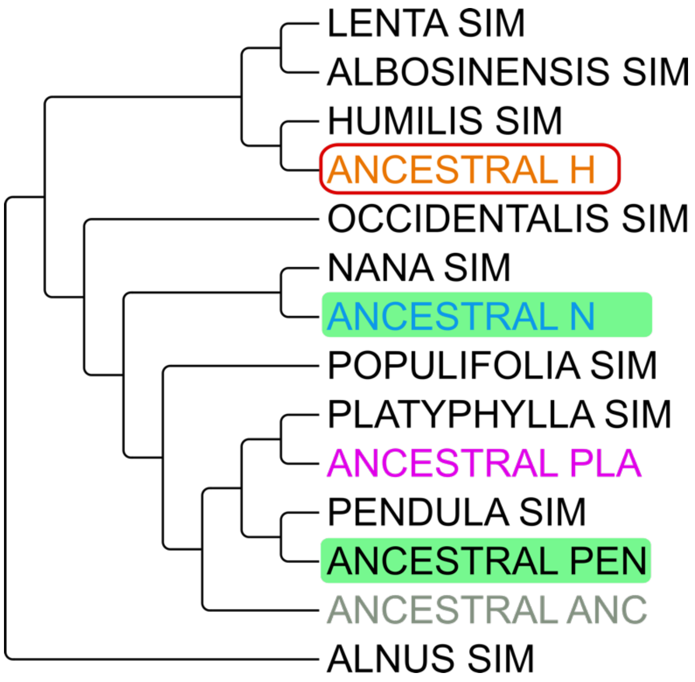

Figure S6 | ***B. pubescens* evolutionary models** – Schematic representation of the nine polyploidization models considered. For each model, different levels of hybridization with one or more third species are hypothesized. Each $H_i$ value represents a fraction of the genome with a specific evolutionary history. For example, parameter $H_2$ in Model 1 represents the fraction of the *B. pubescens* genome that contains alleles of both *B. humilis* and *B. pendula* origin, with the humilis alleles having been acquired via introgression. Simulated annealing and approximate Bayesian computation are used to optimize the $H_i$ values for each model and sampling location. Two extra parameters (not shown in the diagram below) are used to model genus-wide ILS levels, and specific ILS levels between *B. pendula* and *B. platyphylla*. *P*, *N*, *H*, $P_y$ and *A* indicate the hypothetical evolutionary origin of each subgenomic component in the tetraploid (*B. pendula*, *B. nana*, *B. humilis*, *B. platyphylla*, and the common ancestor of *B. pendula*/*B. platyphylla*, respectively). **Insert**: Notice that while the polarization protocol is sensitive to the evolutionary origin of each locus in the polyploid, in general it cannot be used to estimate allele dosage (copy number of each allele).

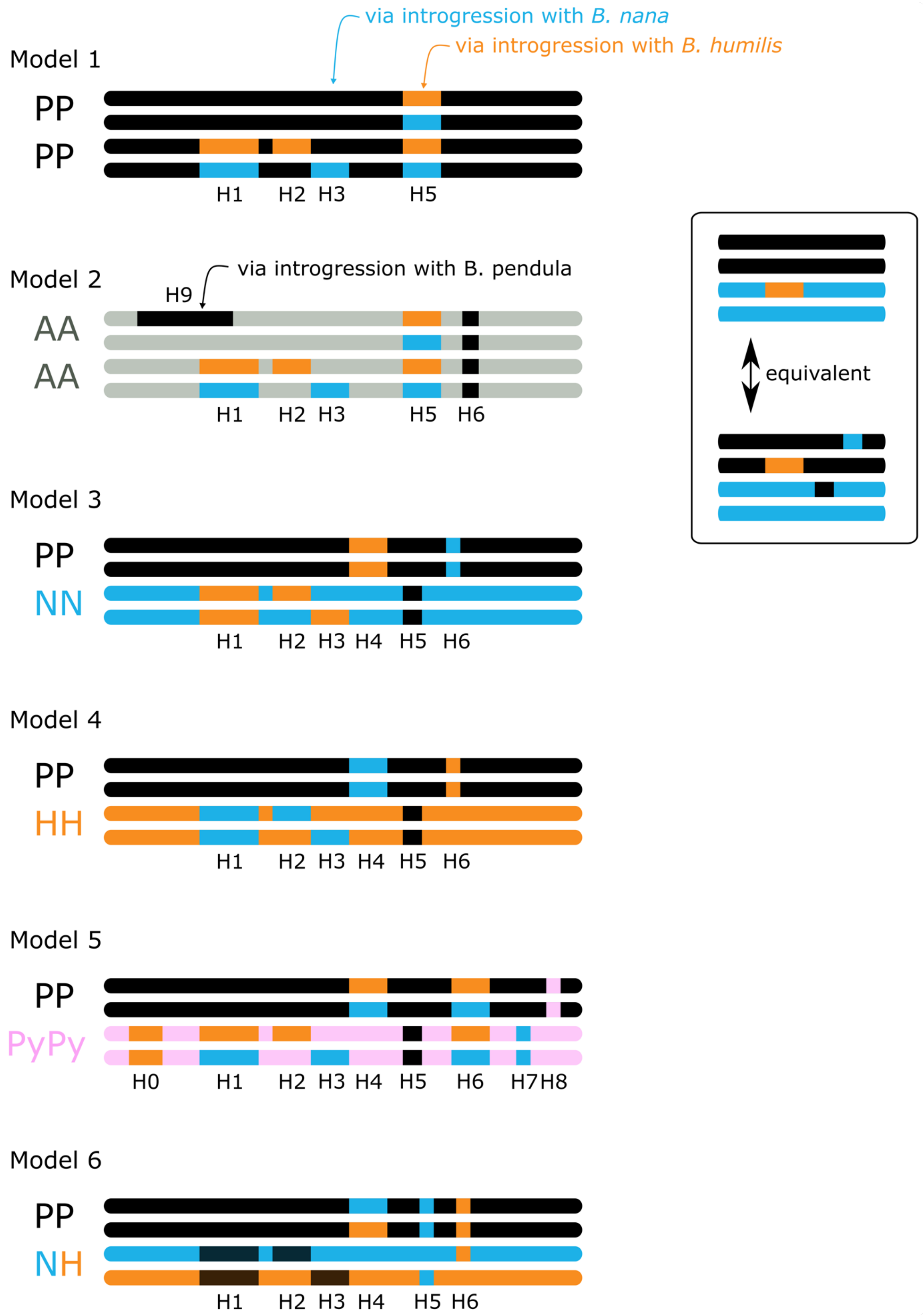
via introgression with B. nana
via introgression with B. humilis
Model 1
PP
PP
H1 H2 H3 H5
Model 2
via introgression with B. pendula
H9
AA
AA
H1 H2 H3 H5 H6
Model 3
PP
NN
H1 H2 H3 H4 H5 H6
Model 4
PP
HH
H1 H2 H3 H4 H5 H6
Model 5
PP
PyPy
H0 H1 H2 H3 H4 H5 H6 H7 H8
Model 6
PP
NH
H1 H2 H3 H4 H5 H6
equivalent

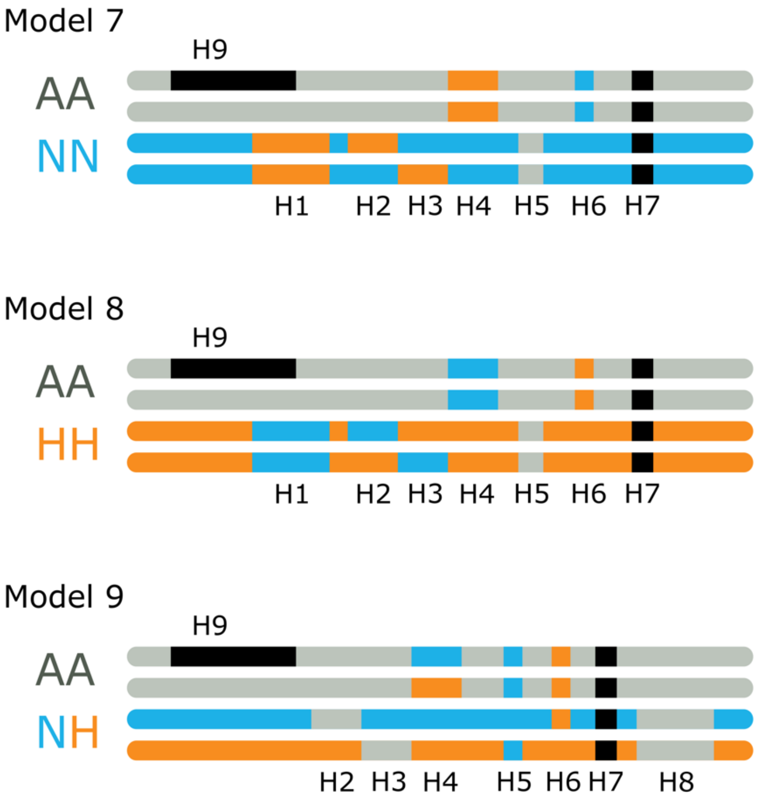
Model 7
H9
AA
NN
H1 H2 H3 H4 H5 H6 H7
Model 8
H9
AA
HH
H1 H2 H3 H4 H5 H6 H7
Model 9
H9
AA
NH
H2 H3 H4 H5 H6 H7 H8

Figure S7 | ***B. pubescens* polyploid-pairing frequencies –** Barplots show the frequency with which *B. pubescens* pairs with other birch species, for different populations and polarization settings, based on phylogenetic analysis of individual gene families using IQ-TREE2. Population numbers correspond to locations shown in Supplementary Fig. S1. The number of specimens per population (N) is also provided. In the x-axis, 'basal' indicates instances when the polyploid is an outgroup to all birch species included in the analysis, 'pend-platy' indicates that the polyploid is an outgroup to the *pendula-platyphylla* clade, 'pop_out' indicates that the polyploid is an outgroup to the *pendula-platyphylla-populifolia* clade, and so on.

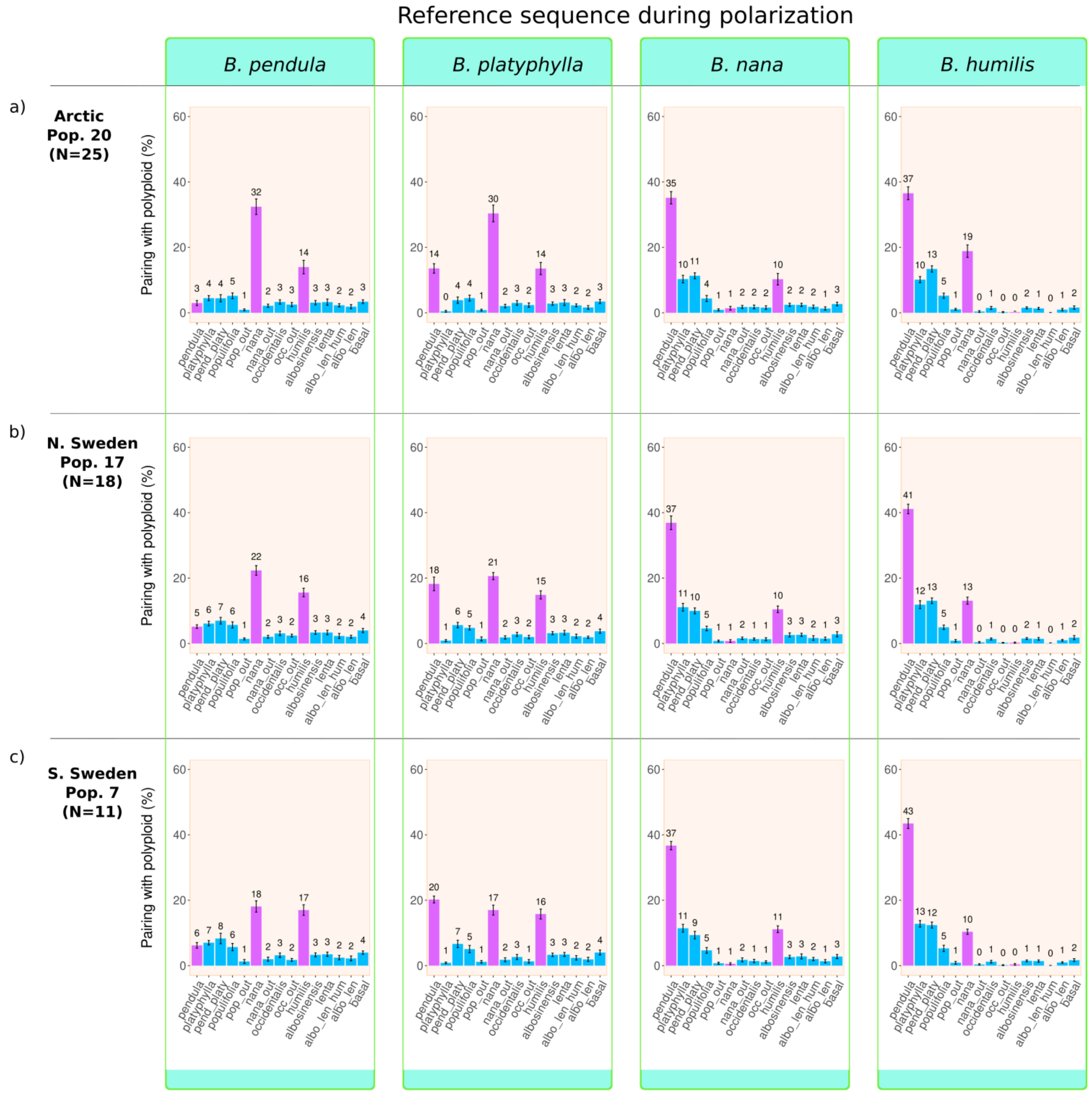

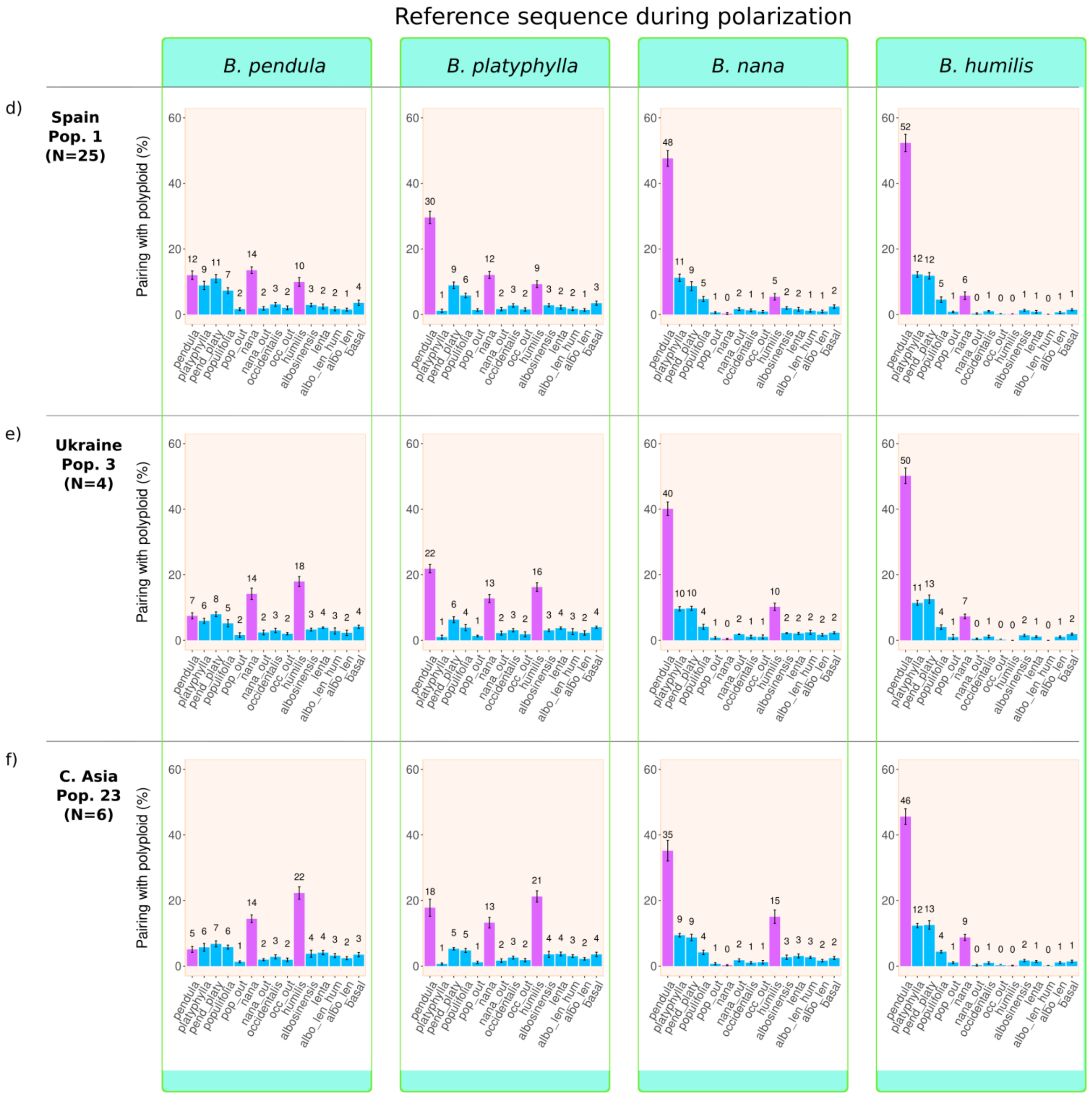
Reference sequence during polarization
B. pendula
B. platyphylla
B. nana
B. humilis
d)
Spain
Pop. 1
(N=25)
e)
Ukraine
Pop. 3
(N=4)
f)
C. Asia
Pop. 23
(N=6)
Pairing with polyploid (%)
0
20
40
60
pendula
platyphylla
pend_platy
populifolia
pop_out
nana
nana_out
occidentalis
occ_out
humilis
albosinensis
lenta
albo_len_hum
albo_len
basal

Figure S8 | **4X *B. pendula* polyploid-pairing frequencies** – Barplots show the frequency with which the autotetraploid *B. pendula* pairs with other birch species, for different polarization settings, and is based on phylogenetic analysis of individual gene families using IQ-TREE2.

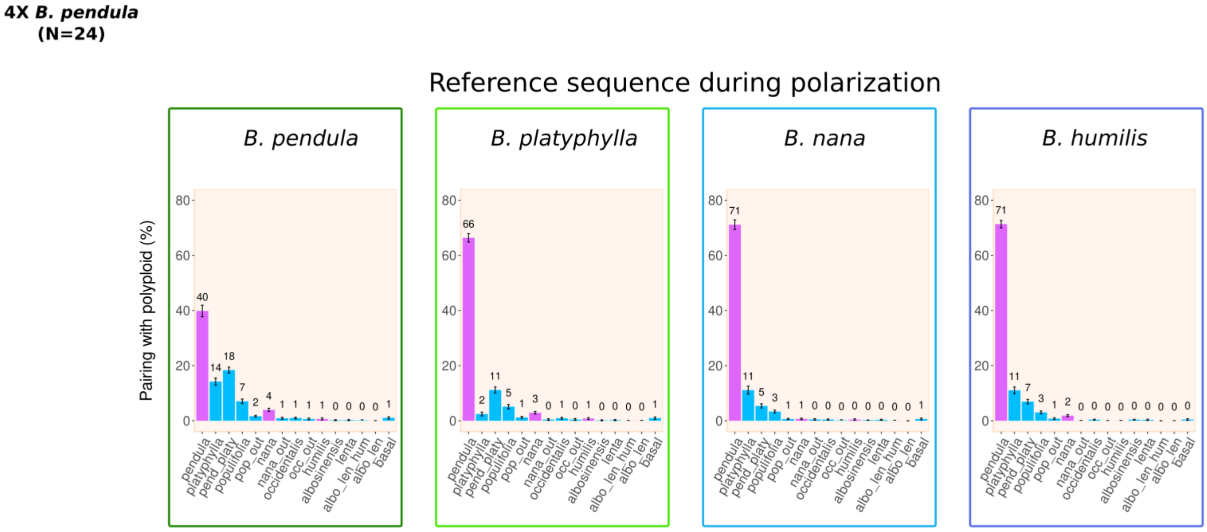

Figure S9 | **STRUCTURE profiles for representative *Betula* populations –** Estimated genetic admixture of 126 *Betula* samples according to the STRUCTURE analysis based on 10,000 variant loci, for K=2 to K= 9. Each individual is represented by a vertical column. Data for diploid (2X) and tetraploid samples (4X) shown separately to facilitate interpretation. Data for tetraploid samples is sorted by latitude from left (south) to right (north). Population location and identifiers (based on Figure S1): ES (1. Spain), PL (2. Poland), UA (3. Ukraine), CA-west (22. Urals, Russia), LT (4. Lithuania), CA-east (23. Nazyvayevsky, Russia), SV-south (7. Skatelov, Sweden), NO (5. southern Norway), FI (21. northern Finland), Arctic (20. Skadi, Norway).

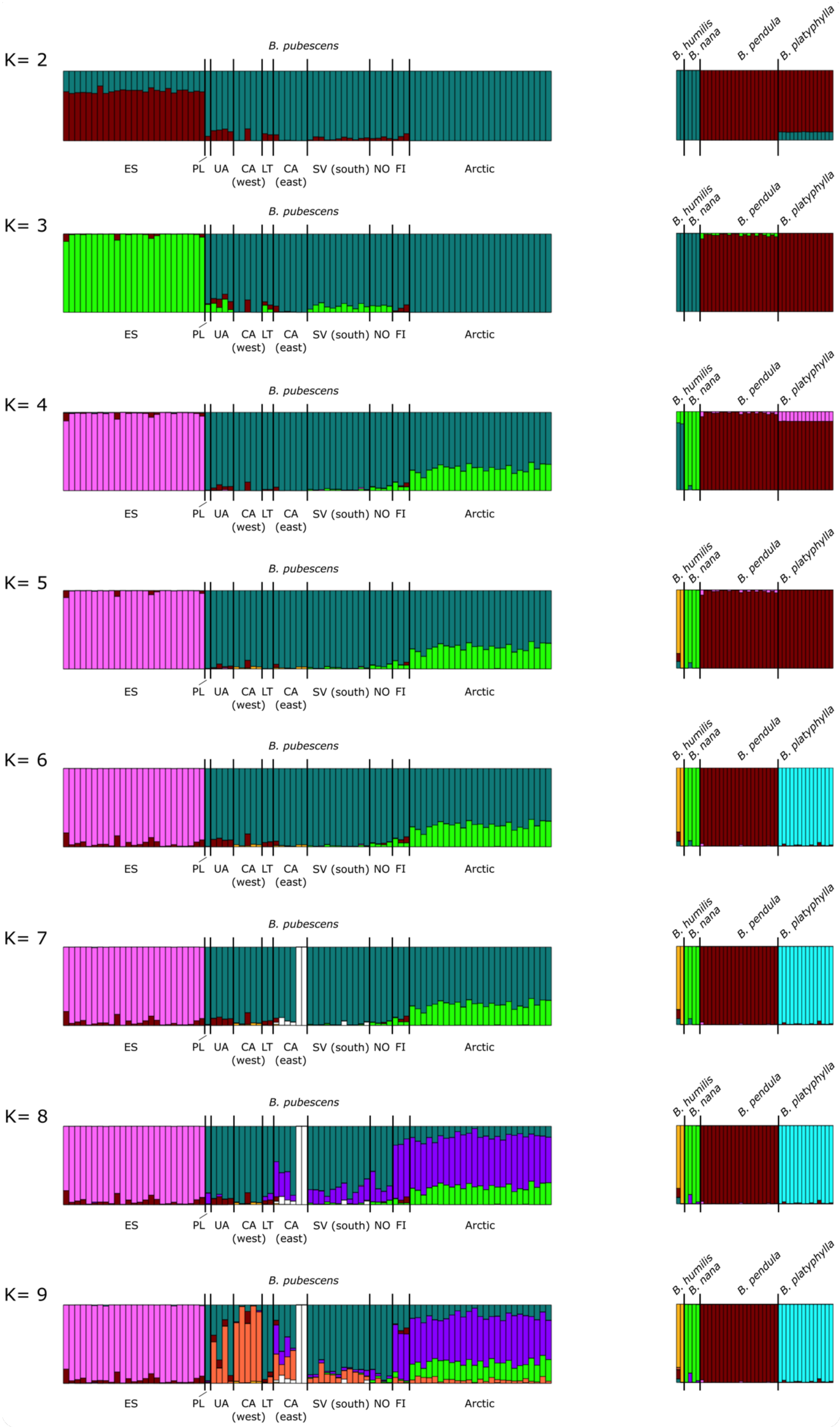
K= 2
K= 3
K= 4
K= 5
K= 6
K= 7
K= 8
K= 9
B. pubescens
ES
PL
UA
CA (west)
LT
CA (east)
SV (south)
NO
FI
Arctic
B. humilis
B. nana
B. pendula
B. platyphylla

Figure S10 | **Clustering analysis based on STRUCTURE results** – Number of clusters determined using the **(a)** Evvano method (Evanno et al. 2005) and **(b)** the ln Pr(X|K) statistic (Pritchard et al. 2000) averaged across replicates.

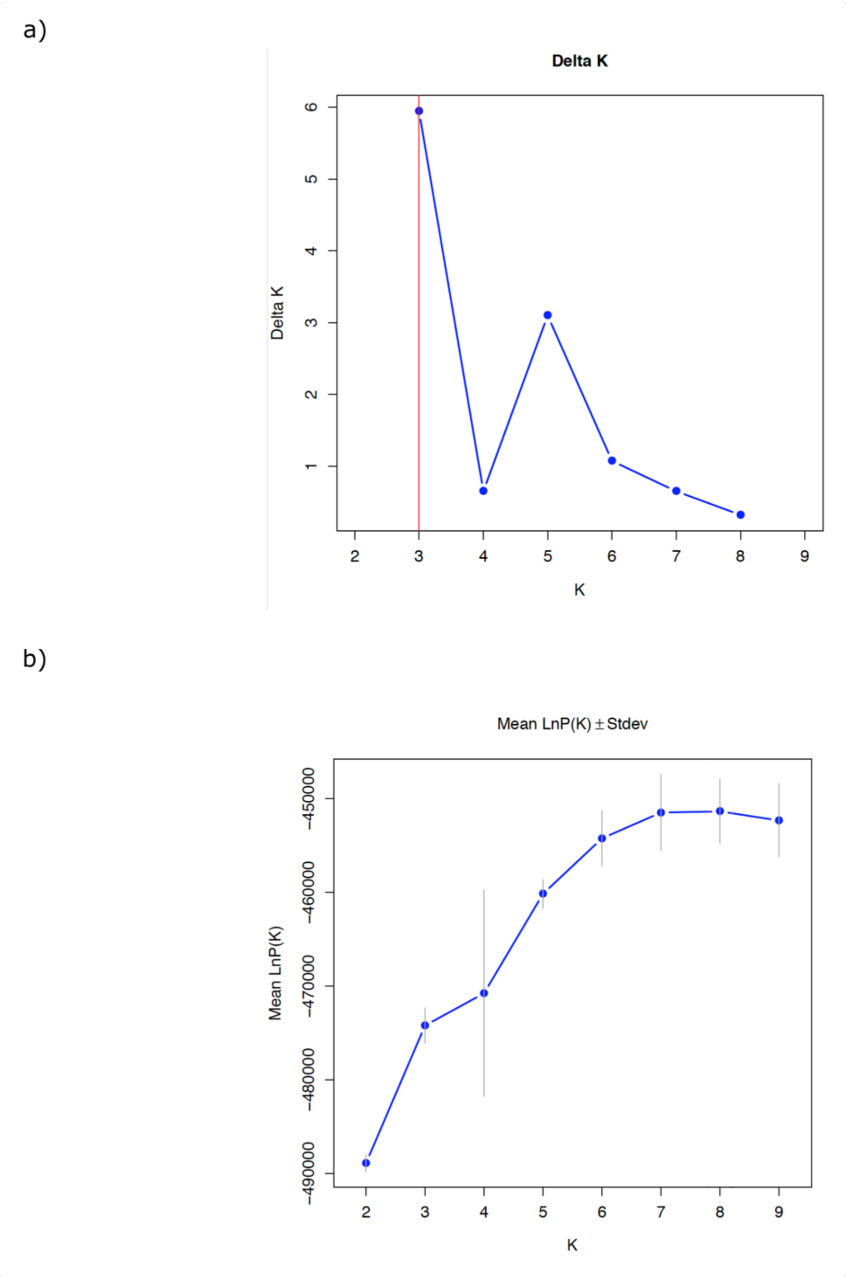

Figure S11 | **DAPC clustering analysis – (a)** DAPC plot for 49 birch accessions based on 50,000 variant loci. Four birch species (*B. nana*, *B. pendula*, *B. platyphylla* and *B. pubescens*) are represented by both whole genome sequencing (WGS) and targeted exome capture (TEC) accessions. *B. pendula* is represented by both diploid (2X) and tetraploid (4X) specimens. TEC accessions: each species/ploidy/population is represented by 5 samples, apart from *B. humilis* (2) and *B. nana* (3). WGS accessions: each species is represented by one sample, apart from *B. pendula* which is represented by 2 samples. *B. pubescens* population location and identifiers (based on Figure S1): Southwestern Europe (1.Spain), Central Europe (2.Poland, 3.Ukraine, and 4.Lithuania), Central Scandinavia (17-Jokkmokk-Sweden), Arctic (20.Skadi-Norway), Central Asia (22-23.Russia). **(b)** Results from k-means clustering analysis of PCA transformed data, based on same dataset described above and obtained using the 'find.clusters' function in *adegenet*, confirmed the presence of five clusters, with cluster membership being organized along species lines. Red dashed line indicates optimal number of clusters based on the Bayesian Information Criterion (BIC).

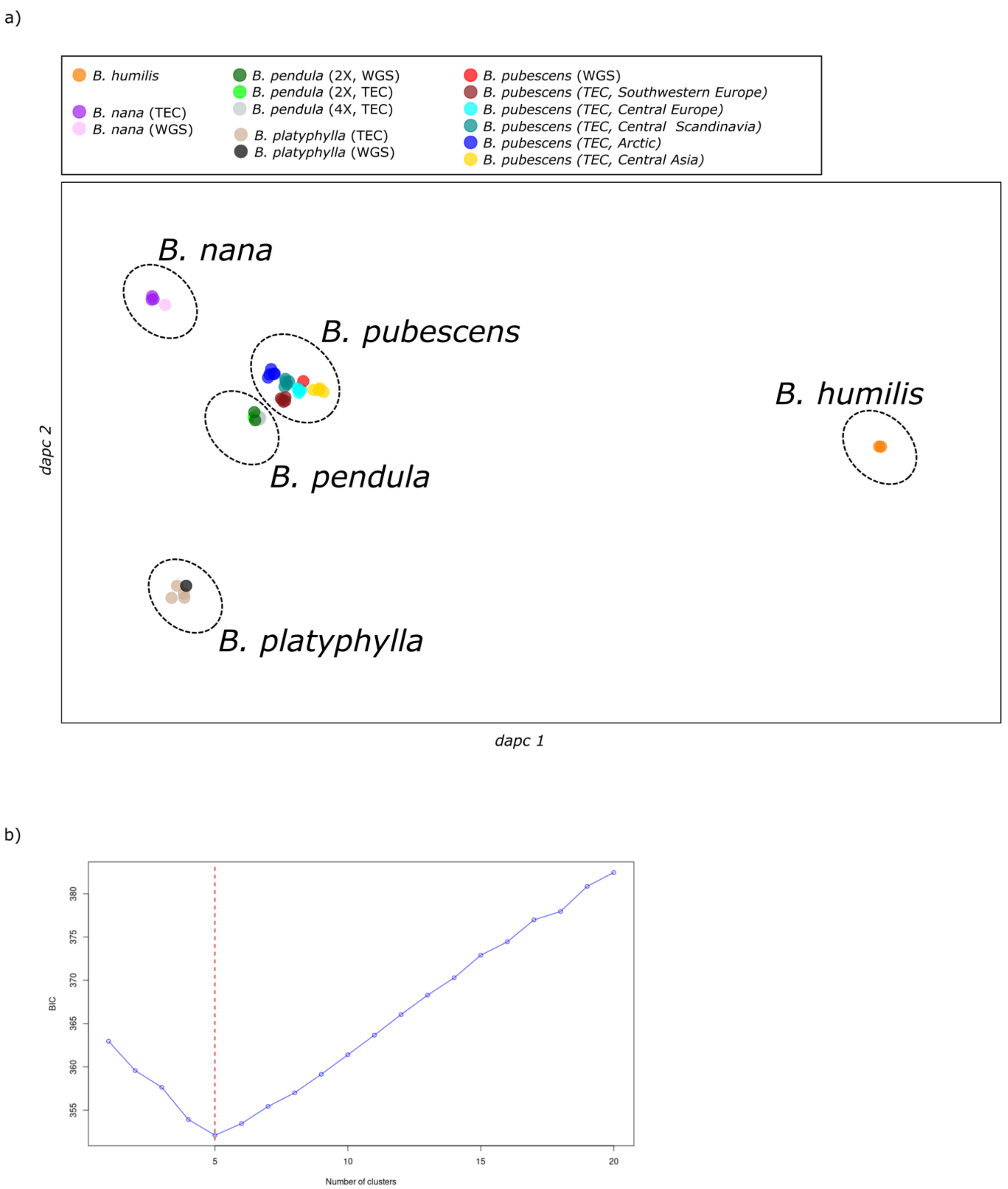
a)
B. humilis
B. nana (TEC)
B. nana (WGS)
B. pendula (2X, WGS)
B. pendula (2X, TEC)
B. pendula (4X, TEC)
B. platyphylla (TEC)
B. platyphylla (WGS)
B. pubescens (WGS)
B. pubescens (TEC, Southwestern Europe)
B. pubescens (TEC, Central Europe)
B. pubescens (TEC, Central Scandinavia)
B. pubescens (TEC, Arctic)
B. pubescens (TEC, Central Asia)
B. nana
B. pubescens
B. humilis
B. pendula
B. platyphylla
dapc 2
dapc 1
b)
BIC
380
375
370
365
360
355
5
10
15
20
Number of clusters

Figure S12 | **Comparison of *B. pubescens* evolutionary models** – This is an extended version of Figure 3 that shows pairing frequencies for all nine models and also for multi-species clades. In the x-axis, 'basal' indicates instances when the polyploid is an outgroup to all birch species included in the analysis, 'pend-platy' indicates that the polyploid is an outgroup to the *pendula-platyphylla* clade, 'pop_out' indicates that the polyploid is an outgroup to the *pendula-platyphylla-populifolia* clade, and so on.

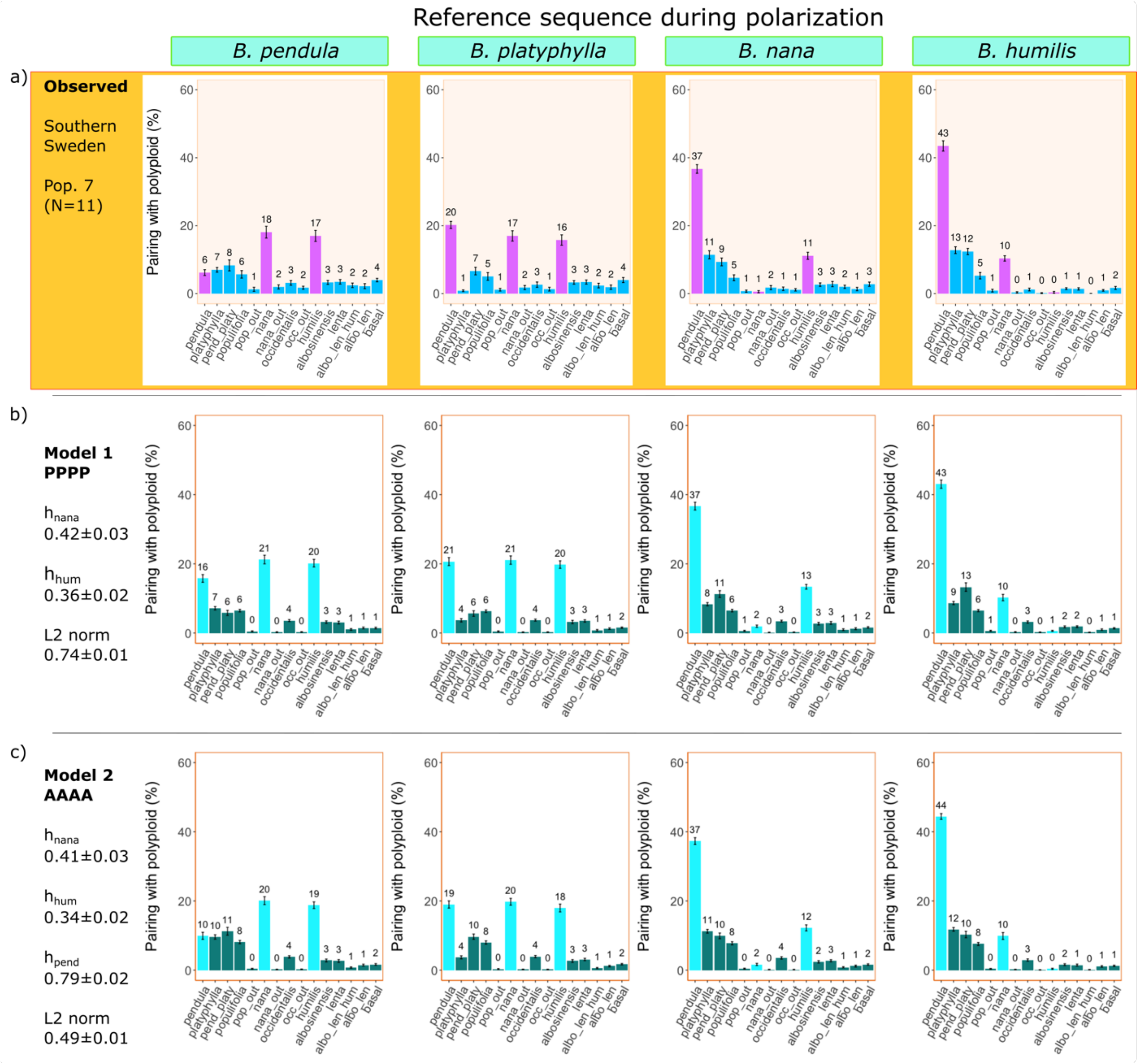

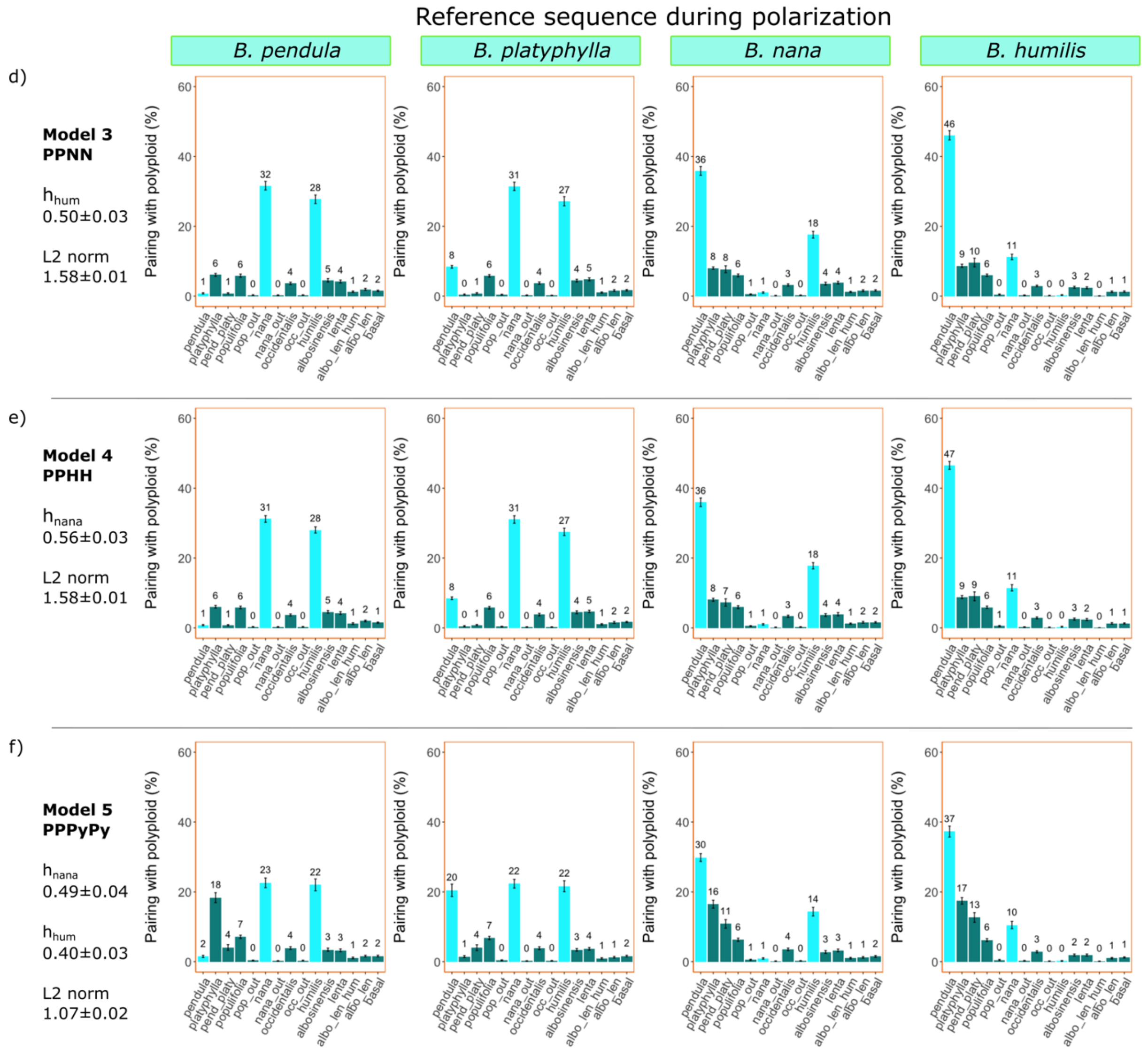
Reference sequence during polarization
B. pendula
B. platyphylla
B. nana
B. humilis
d)
Model 3
PPNN
h_hum
0.50±0.03
L2 norm
1.58±0.01
e)
Model 4
PPHH
h_nana
0.56±0.03
L2 norm
1.58±0.01
f)
Model 5
PPPyPy
h_nana
0.49±0.04
h_hum
0.40±0.03
L2 norm
1.07±0.02
Pairing with polyploid (%)
pendula
platyphylla
pend_platy
populifolia
pop_out
nana
nana_out
occidentalis
occ_out
humilis
albosinensis
lenta
albo_len_hum
albo_len
basal

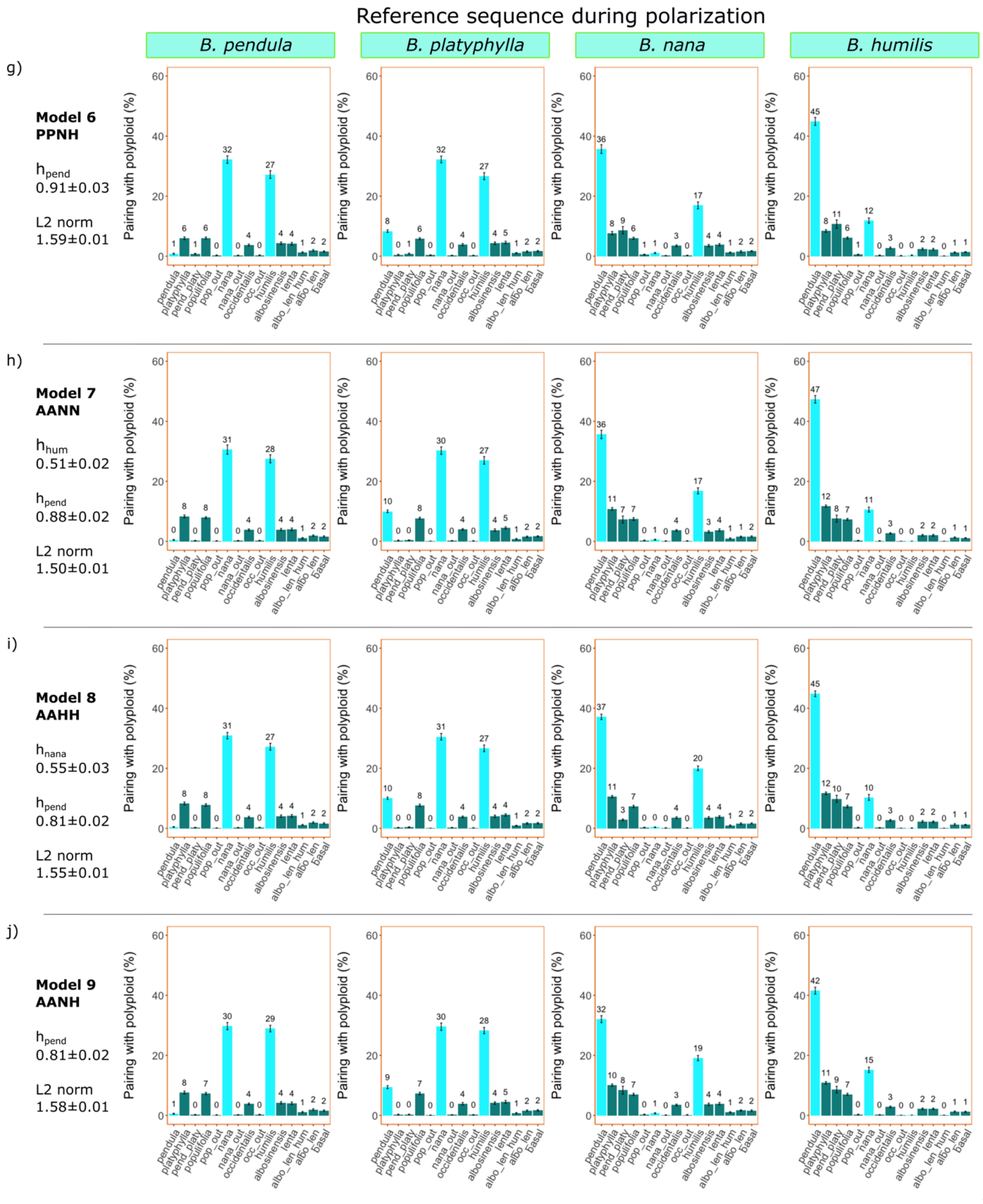
Reference sequence during polarization
B. pendula
B. platyphylla
B. nana
B. humilis
g)
Model 6
PPNH
h_pend
0.91±0.03
L2 norm
1.59±0.01
h)
Model 7
AANN
h_hum
0.51±0.02
h_pend
0.88±0.02
L2 norm
1.50±0.01
i)
Model 8
AAHH
h_nana
0.55±0.03
h_pend
0.81±0.02
L2 norm
1.55±0.01
j)
Model 9
AANH
h_pend
0.81±0.02
L2 norm
1.58±0.01
Pairing with polyploid (%)
pendula
platyphylla
pend_platy
populifolia
pop_out
nana
nana_out
occidentalis
occ_out
humilis
albosinensis
lenta
albo_len_hum
albo_len
basal

Figure S13 | **AAAA autopolyploid model predictions** – Barplots show the frequency with which the polarized *B. pubescens* sequence pairs with other birch species, as predicted by the AAAA evolutionary model and for different polarization settings, based on phylogenetic analysis of individual gene families using IQ-TREE2. "AA" indicates the hypothetical evolutionary origin of each subgenomic component in the tetraploid (common ancestor of *B. pendula* and *B. platyphylla*). Modeling data summarizes optimized results [top 5% simulation runs (50 out of 1000 runs)], obtained using approximate Bayesian computation having as a reference the values observed experimentally (shown on top panel for each population). Each simulation run averages results over 25 independent replicates. Cyan bars indicate level of polyploid-pairing for the three key birch species. Bars in red indicate discrepancies between predicted and experimental results higher than 5 percentage points. The weighted *L2* norm distance, averaged over the top 5% simulation runs, measures the goodness of fit and includes data for all four polarizations (lower *L2* values denote a better fit). *pend_platy* indicates instances when the polarized polyploid sequence is an outgroup to the pendula-platyphylla clade.

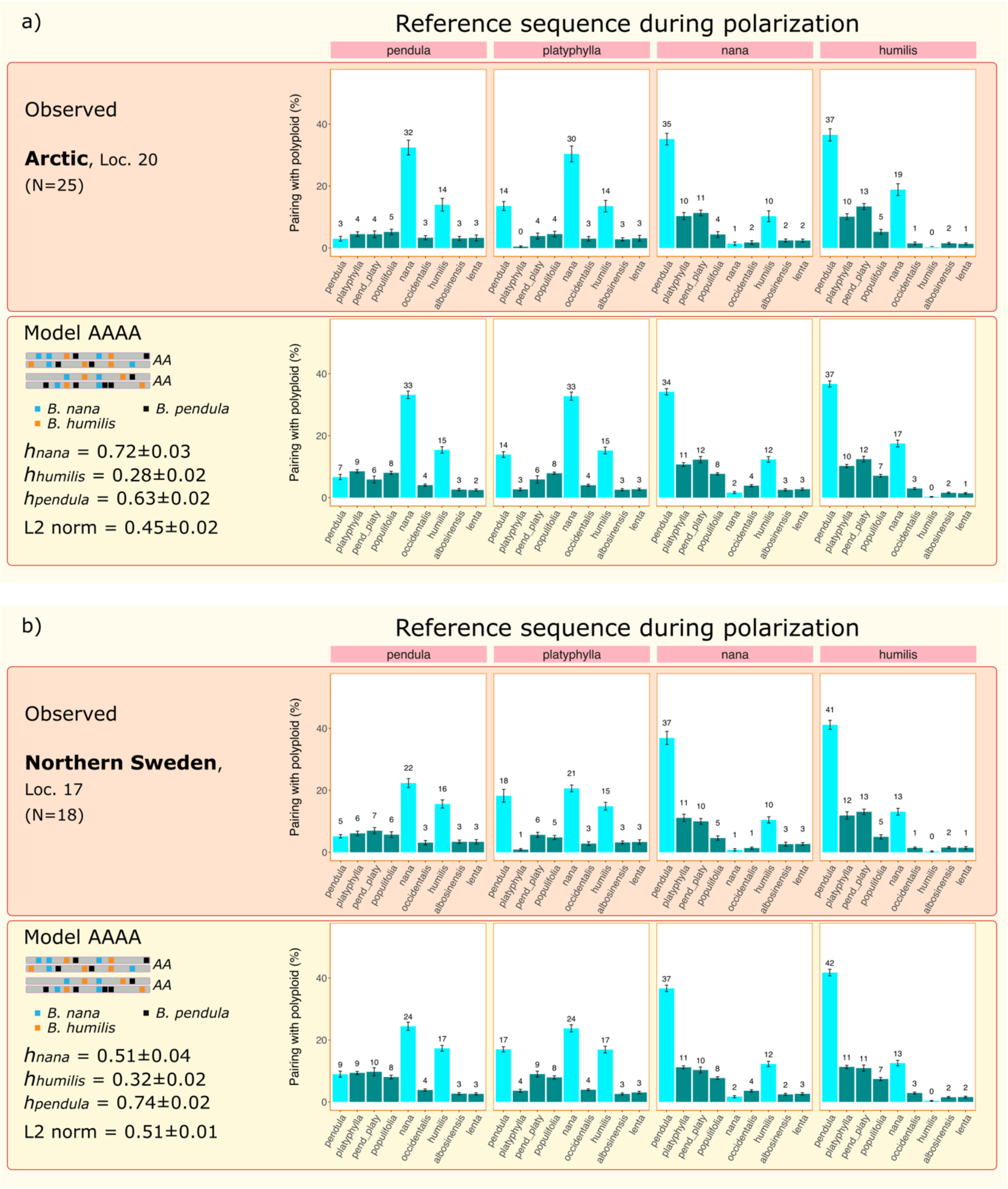
a)
Reference sequence during polarization
pendula
platyphylla
nana
humilis
Observed
Arctic, Loc. 20
(N=25)
Pairing with polyploid (%)
Model AAAA
AA
AA
B. nana
B. pendula
B. humilis
hnana = 0.72±0.03
hhumilis = 0.28±0.02
hpendula = 0.63±0.02
L2 norm = 0.45±0.02
pendula platyphylla pend_platy populifolia nana occidentalis humilis albosinensis lenta
b)
Reference sequence during polarization
Observed
Northern Sweden,
Loc. 17
(N=18)
Model AAAA
hnana = 0.51±0.04
hhumilis = 0.32±0.02
hpendula = 0.74±0.02
L2 norm = 0.51±0.01

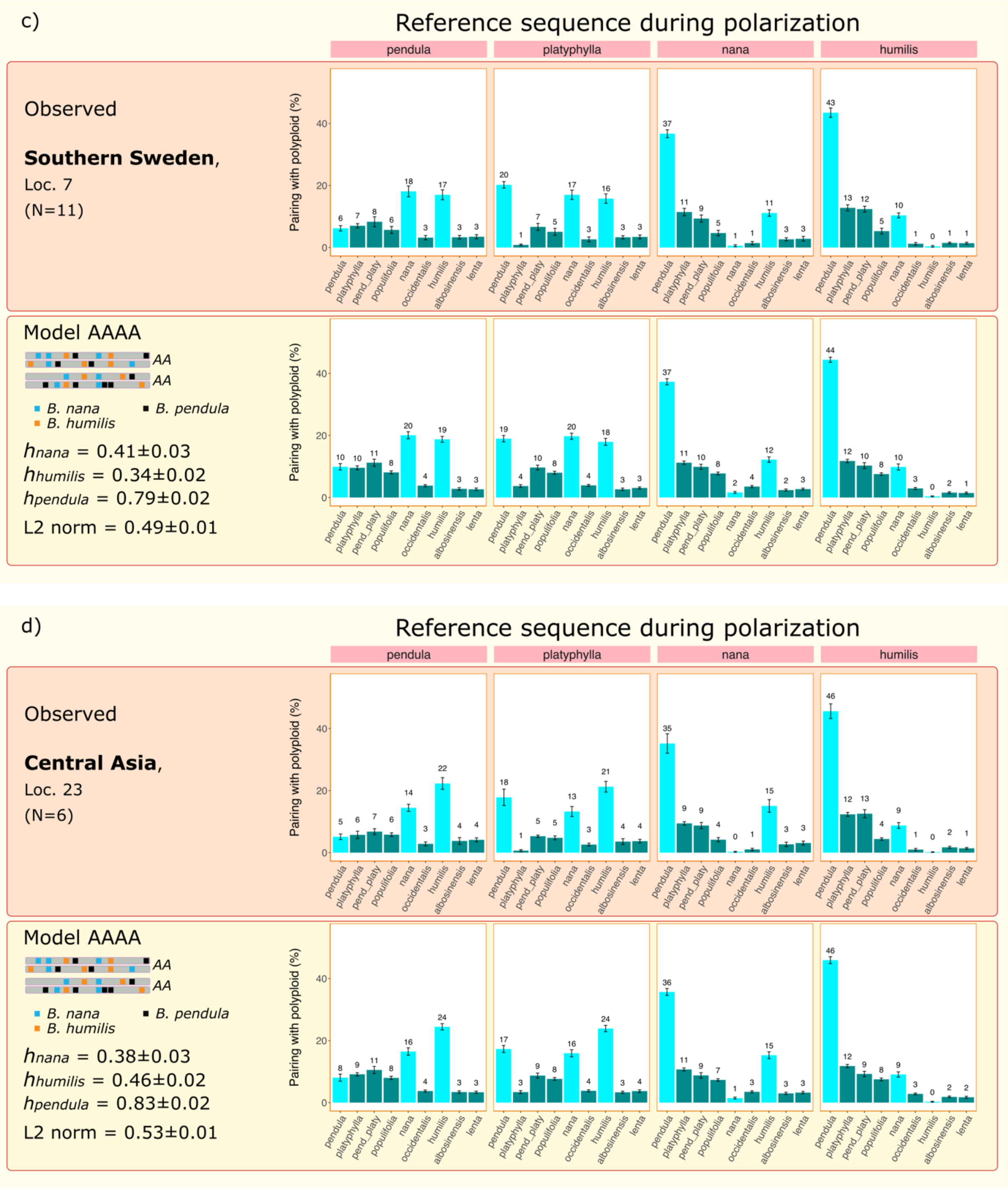
c)
Reference sequence during polarization
pendula
platyphylla
nana
humilis
Observed
Southern Sweden,
Loc. 7
(N=11)
Pairing with polyploid (%)
pendula
platyphylla
pend_platy
populifolia
nana
occidentalis
humilis
albosinensis
lenta
Model AAAA
AA
AA
B. nana
B. pendula
B. humilis
hnana = 0.41±0.03
hhumilis = 0.34±0.02
hpendula = 0.79±0.02
L2 norm = 0.49±0.01
d)
Reference sequence during polarization
Observed
Central Asia,
Loc. 23
(N=6)
Model AAAA
hnana = 0.38±0.03
hhumilis = 0.46±0.02
hpendula = 0.83±0.02
L2 norm = 0.53±0.01

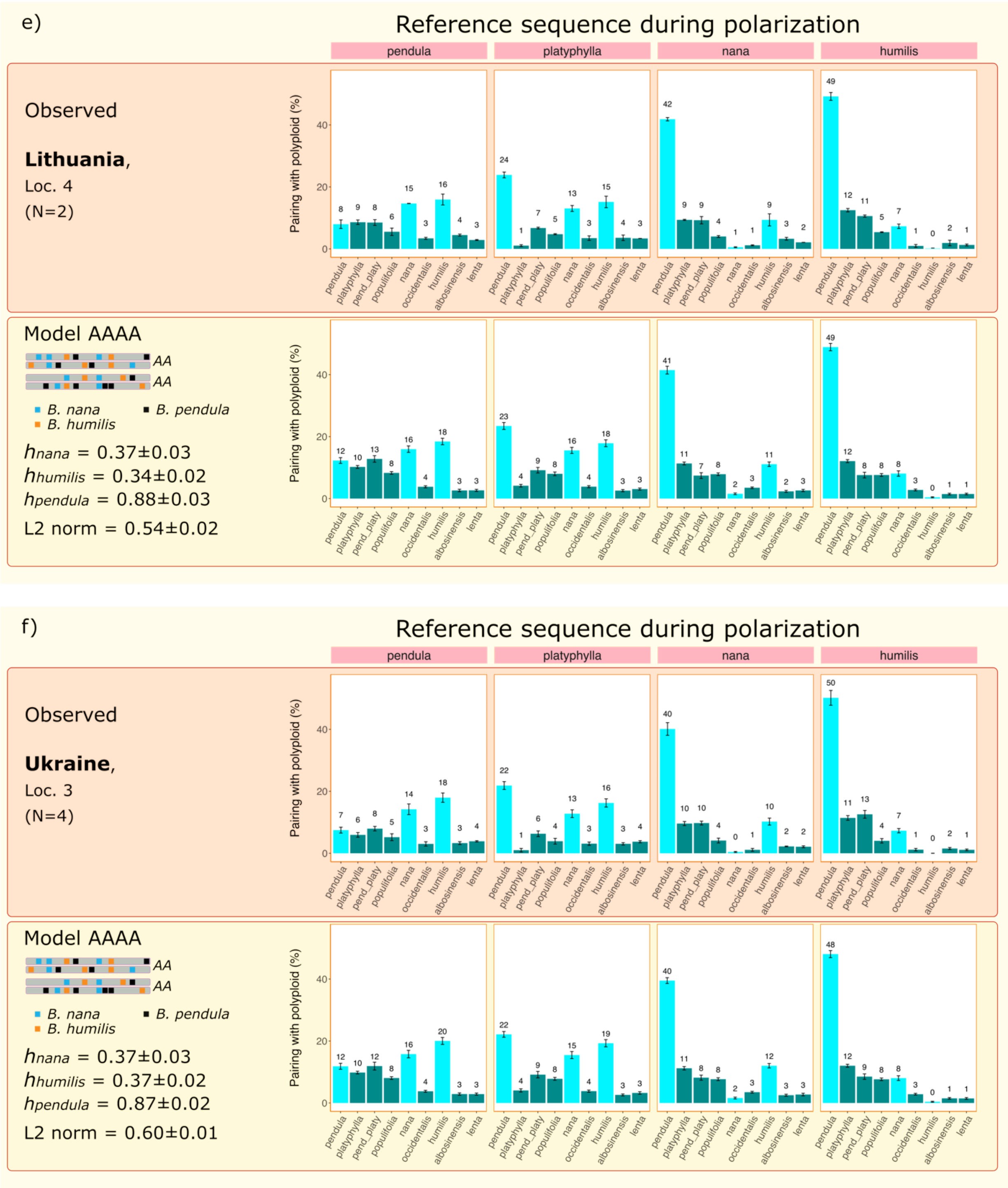
e)
Reference sequence during polarization
pendula
platyphylla
nana
humilis
Observed
Lithuania,
Loc. 4
(N=2)
Pairing with polyploid (%)
pendula
platyphylla
pend_platy
populifolia
nana
occidentalis
humilis
albosinensis
lenta
Model AAAA
AA
AA
B. nana
B. pendula
B. humilis
hnana = 0.37±0.03
hhumilis = 0.34±0.02
hpendula = 0.88±0.03
L2 norm = 0.54±0.02
f)
Reference sequence during polarization
Observed
Ukraine,
Loc. 3
(N=4)
Model AAAA
hnana = 0.37±0.03
hhumilis = 0.37±0.02
hpendula = 0.87±0.02
L2 norm = 0.60±0.01

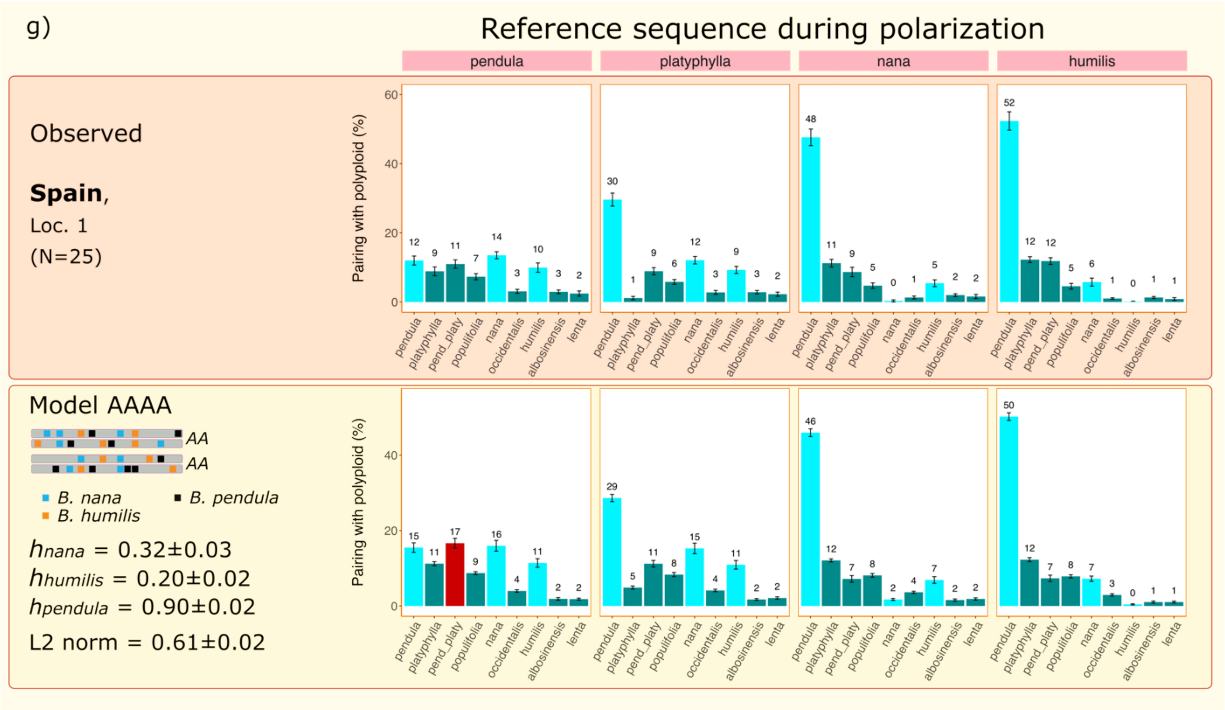
g)
Reference sequence during polarization
pendula
platyphylla
nana
humilis
Observed
**Spain**,
Loc. 1
(N=25)
Pairing with polyploid (%)
pendula platyphylla pend_platy populifolia nana occidentalis humilis albosinensis lenta
Model AAAA
AA
AA
B. nana
B. pendula
B. humilis
$h_{nana}$ = 0.32±0.03
$h_{humilis}$ = 0.20±0.02
$h_{pendula}$ = 0.90±0.02
L2 norm = 0.61±0.02
